# selscan 2.0: scanning for sweeps in unphased data

**DOI:** 10.1101/2021.10.22.465497

**Authors:** Zachary A. Szpiech

## Abstract

**Summary:** Several popular haplotype-based statistics for identifying recent or ongoing positive selection in genomes require knowledge of haplotype phase. Here we provide an update to selscan which implements a re-definition of these statistics for use in unphased data.

**Availability and Implementation:** Source code and binaries freely available at https://github.com/szpiech/selscan, implemented in C/C++ and supported on Linux, Windows, and MacOS.

**Supplemental Information:** Online supplemental information available

## 1 Introduction

Haplotype-based summary statistics—such as iHS (Voight, et al., 2006), nSL (Ferrer-Admetlla, et al., 2014), XP-EHH (Sabeti, et al., 2007), and XP-nSL (Szpiech, et al., 2021)—have become commonplace in evolutionary genomics studies to identify recent and ongoing positive selection in populations (e.g.,Colonna, et al., 2014; Crawford, et al., 2017; Lu, et al., 2019; Meier, et al., 2018; Nedelec, et al., 2016; Salmon, et al., 2021; Zhang, et al., 2020; Zoledziewska, et al., 2015). When an adaptive allele sweeps through a population, it leaves a characteristic pattern of long high-frequency haplotypes and low genetic diversity in the vicinity of the allele.

These statistics aim to capture these signals by summarizing the decay of haplotype homozygosity as a function of distance from a putatively selected region, either within a single population (iHS and nSL) or between two populations (XP-EHH and XP-nSL).

These haplotype-based statistics are powerful for detecting recent positive selection (Colonna, et al., 2014; Crawford, et al., 2017; Lu, et al., 2019; Meier, et al., 2018; Nedelec, et al., 2016; Salmon, et al., 2021; Zhang, et al., 2020; Zoledziewska, et al., 2015), and the two-population versions can even out-perform pairwise Fst scans on a large swath of the parameter space (Szpiech, et al., 2021). Furthermore, haplotype-based methods have also been shown to be robust to background selection (Fagny, et al., 2014; Schrider, 2020). However, each of these statistics presumes that haplotype phase is known or well-estimated.

As the generation of genomic sequencing data for non-model organisms is becoming routine (Ellegren, 2014), there are many great opportunities for studying recent adaptation across the tree of life (e.g., Campagna and Toews (2022)). However, often these organisms/populations do not have a well-characterized demographic history or recombination rate map, two pieces of information which are important inputs for statistical phasing methods (Browning, et al., 2021; Delaneau, et al., 2013).

Recent work has shown that haplotype-based statistics can be adapted for use on unphased data (Klassmann and Gautier, 2022) and that converting haplotype data into “multi-locus genotype” data is an effective approach for using haplotype-based selection statistics such as G12, LASSI, and saltiLASSI (DeGiorgio and Szpiech, 2022; Harris and DeGiorgio, 2020; Harris, et al., 2018) in unphased data. Recognizing this, we have reformulated the iHS, nSL, XP-EHH, and XP-nSL statistics to use multi-locus genotypes and provided an easy-to-use implementation in selscan 2.0 (Szpiech and Hernandez, 2014). We evaluate the performance of these unphased statistics under various generic demographic models and compare against the original statistics applied to simulated datasets when phase is either known or unknown.

## 2 Methods

When the --unphased flag is set in selscan v2.0+, biallelic genotype data is collapsed into multi-locus genotype data by representing the genotype as either 0, 1, or 2—the number of derived alleles observed. In this case, selscan v2.0+ will then compute iHS, nSL, XP-EHH, and XP-nSL as described below. We follow the notation conventions of Szpiech and Hernandez (2014).

### 2.1 Extended Haplotype Homozygosity

In a sample of n diploid individuals, let 𝒞 denote the set of all possible genotypes at locus *x*_0_. For multi-locus genotypes, 𝒞 ≔ {0,1,2}, representing the total counts of a derived allele. Let 𝒞 (*x*_*i*_) be the set of all unique haplotypes extending from site *x*_0_ to site *x*_*i*_ either upstream or downstream of *x*_0_. If *x*_1_ is a site immediately adjacent to *x*_0_, then 𝒞 (*x*_1_) ≔ {00,01,02,10,11,12,20,21,22}, representing all possible two-site multi-locus genotypes. We can then compute the extended haplotype homozygosity (EHH) of a set of multi-locus genotypes as

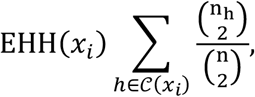

where *n*_*h*_ is the number of observed haplotypes of type *h*.

If we wish to compute the EHH of a subset of observed haplotypes that all contain the same ‘core’ multi-locus genotype, let ℋ _*c*_(*x*_*i*_) be the partition of 𝒞 (*x*_*i*_) containing genotype c ∈ 𝒞 at *x*_0_. For example, choosing a homozygous derived genotype (c = 2) as the core, ℋ _2_ ≔ {20,21,22}. Thus, we can compute the EHH of all individuals carrying a given genotype at site *x*_0_ extending out to site *x*_*i*_ as

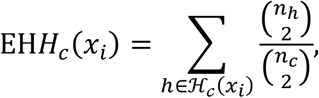

where *n*_*h*_ is the number of observed haplotypes of type *h* and *n*_*c*_ is the number of observed multi-locus genotypes with core genotype of c. Finally, we can compute the complement EHH of a sample of multi-locus genotypes as

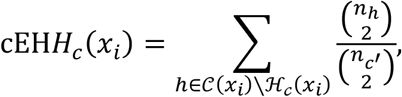

where *n*_c’_ is the number of observed multi-locus genotypes with a core genotype of not *c*.

### 2.2 iHS and nSL

Unphased iHS and nSL are calculated using the equations above. First, we compute the integrated haplotype homozygosity (iHH) for the homozygous ancestral (*c* = 0) and derived (*c* = 2) core genotypes as

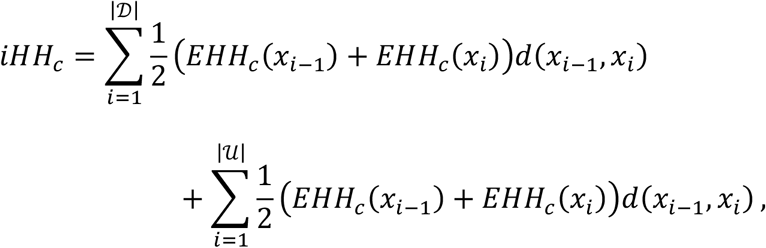

where 𝒟 is the set of downstream sites from the core locus and 𝒰 is the set of upstream sites. *d*(*x*_*i* −1_, *x*_*i*_) is a measure of genomic distance between to markers and is the genetic distance in centimorgans or physical distance in basepairs for iHS (Voight, et al., 2006) or the number of sites observed for nSL (Ferrer-Admetlla, et al., 2014). We similarly compute the complement integrated haplotype homozygosity (ciHH) for both homozygous core genotypes as

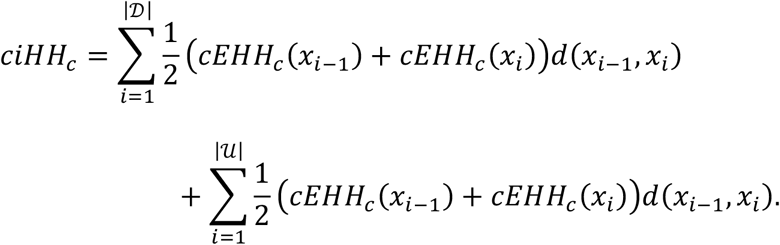

The (unstandardized) unphased iHS is then calculated as

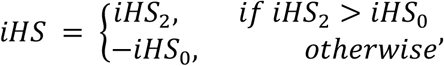

where 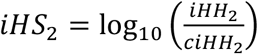 and 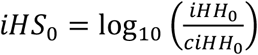. Conceptually, this is nearly identical to the phased version of iHS, where the log ratio of the integrated haplotype homozygosity is computed between all haplotypes carrying the ancestral allele at the core locus versus all haplotypes carrying the derived allele at the core locus. In this case, however, we compare the iHH of the haplotypes containing homozygous genotypes of one allele at the core locus to the iHH of the haplotypes containing all other genotypes at the core locus. Doing this for both homozygous derived and homozygous ancestral haplotypes separately, we then choose the most extreme value. We assign a positive sign for long low-diversity haplotypes containing the derived homozygous genotype at the core locus, and we assign a negative sign for long low-diversity haplotypes containing the ancestral homozygous genotype at the core locus.

Unstandardized iHS scores are then normalized in frequency bins, as previously described (Ferrer-Admetlla, et al., 2014; Voight, et al., 2006). Unstandardized unphased nSL is computed similarly with the appropriate distance measure (see Ferrer-Admetlla, et al. (2014) where they show that nSL can be reformulated as iHS with a different distance measure). Large positive scores indicate long high-frequency haplotypes with a homozygous derived core genotype, and large negative scores indicate long high-frequency haplotypes with a homozygous ancestral core genotype. Clusters of extreme scores in both directions indicate evidence for a sweep.

### 2.3 XP-EHH and XP-nSL

Unphased XP-EHH and XP-nSL are calculated by comparing the iHH between populations *A* and *B*, using the entire sample in each population. iHH in a population P is computed as

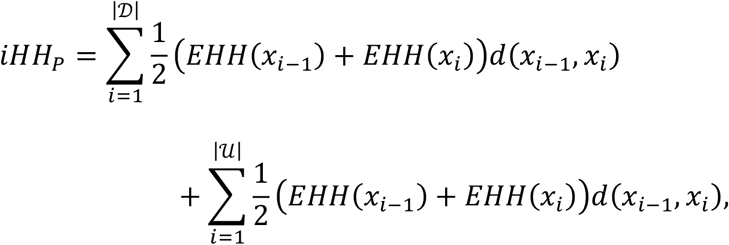

where the distance measure is given as centimorgans or basepairs for XP-EHH (Sabeti, et al., 2007) and number of sites observed for XP-nSL (Szpiech, et al., 2021). The XP statistics between population *A* and *B* are then computed as 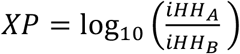 and are normalized genome wide. Large positive scores indicate long high-frequency haplotypes in population *A*, and large negative scores indicate long high-frequency haplotypes in population *B*. Clusters of extreme scores in one direction indicate evidence for a sweep in that population.

## 3 Results

We find that the unphased versions of iHS and nSL generally have good power at large sample sizes (Figures 1A, 1B, S1, S7, and S8) to detect selection prior to fixation of the allele, with nSL generally outperforming iHS. In smaller populations (Figures S1C and S1D), power does suffer relative to larger populations (Figures S1A, S1B, S1E, S1F). We note that these statistics struggle to identify soft sweeps when the population is undergoing exponential growth (Figures S1E and S1F). Each of these statistics also have low false positive rates hovering around 1% (Tables S2-S5). These single-population statistics only perform well for relatively large samples (Figures 1A, 1B, and Figures S19, S25, S26, S31, S32, S37, S43, S44, S55, S61, S62).

**Figure 1.**
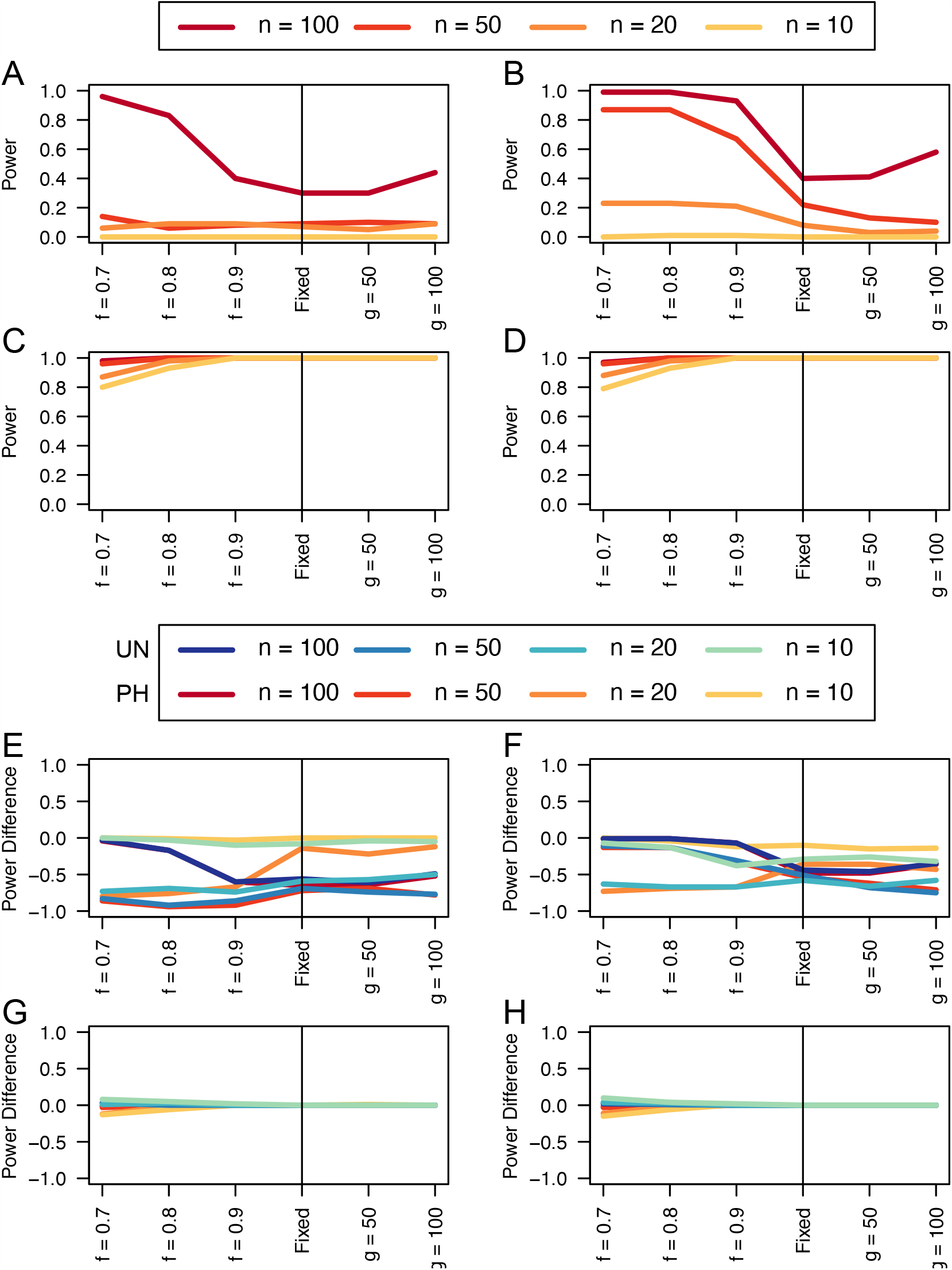
Power curves for unphased implementations of iHS (A), nSL (B), XP-EHH (C), and XP-nSL (D), and power difference between unphased implementations of iHS (E), nSL (F), XP-EHH (G), and XP-nSL (H) and phased implementations. Blue curves represent the power difference between the unphased and phased statistics when applied to unphased data (UN). Red curves represent the power difference between the unphased and phased statistics when applied to perfectly phased data (PH). Values greater than 0 indicate the unphased statistic had higher power. All panels represent analyses with demographic history Demo 1 and n = 100, 50, 20, or 10 diploid samples. For these plots the selection coefficient is set at *s* = 0.01, the frequency at which selection began is set at *e* = 0 (i.e., a hard sweep), and the divergence time in generations is set at *t*_*d*_ = 2000. *f* is the frequency of the adaptive allele at time of sampling, *g* is the number of generations at time of sampling since fixation.

Similarly, we find that the unphased versions of XP-EHH and XP-nSL have good power as well even for relatively low sample sizes (Figures 1C, 1D, 1G, 1H and Figures S2, S3, S9-S12, S20, S21, S27-30, S38, S39, S45-48, S56, S57, S63-S66). When the sweep takes place in the smaller of the two populations (Figures S2C, S2D, S20C, S20D, S38D, S38C, S56C, and S56D), we see a similar decrease in power, likely related to the lower efficiency of selection in small populations. When one population is undergoing exponential growth (Figures S3, S21, S39, S57) performance is generally quite good, likely the result of a larger effective selection coefficient in large populations. These two-population statistics generally outperform their single-population counterparts, especially at small diploid sample sizes and for sweeps that have reached fixation recently. Each of these statistics also have low false positive rates hovering around 1% (Tables S2-S5).

Next, we turn to comparing the performance of these unphased statistics to their phased counterparts when they are used to analyze either phased data or unphased data. In Figures 1E, 1F, 1G, 1H and Figures S4-S6, S13-S18, S22-S24, S31-S36, S40-S42, S49-S54, S58-S60, S67-S72) we plot the difference in power between the unphased statistics and the phased counterpart applied to data with phase known (red lines) or phase scrambled (blue lines). Where these lines are greater than or equal to 0 indicates that the unphased statistic performed as well as or better than the phased counterpart.

We find that iHS tends to underperform the traditional phased implementations, but nSL tends to perform as well as the phased versions (Figures 1E, 1F and Figures S4, S13, S14, S22, S31, S32, S40, S49, S50, S58, S67, S68). Although we note noticeable drops in unphased nSL power for softer sweeps in exponential growth scenarios (Figures S4F, S13F, S14F, S22F, S31F, S32F, S40F, S49F, S50F, S58F, S67F, S68F) and for sweeps near completion in small population sizes (Figures S4E, S13E, S14E, S22E, S31E, S32E, S40E, S49E, S50E, S58E, S67E, S68E).

When comparing the unphased versions of XP-EHH and XP-nSL, we find that they consistently perform as well or better than their phased counterparts (Figures 1G, 1H and Figures S5, S6, S17, S18, S23, S24, S35, S36, S41, S42, S53, S54, S59, S60, S71, S72), except in limited circumstances where phase is known, and the sweep is fairly young (sweeping allele at 0.7 frequency) or the divergence time is further in the past.

## 4 Discussion

We introduce multi-locus genotype versions of four popular haplotype-based selection statistics—iHS (Voight, et al., 2006), nSL (Ferrer-Admetlla, et al., 2014), XP-EHH (Sabeti, et al., 2007), and XP-nSL (Szpiech, et al., 2021)—that can be used when the phase of genotypes is unknown. Although phase would seem to be a critically important component of any haplotype-based method for detecting selection, here we show that, by collapsing haplotypes into derived allele counts (thus erasing phase information), we can achieve similar power to using this information. We observed that single-population statistics such as iHS and nSL require relatively large diploid sample sizes (*n* >= 100 for iHS, *n* >= 50 for nSL), but the two-population statistics XP-EHH and XP-nSL perform well even for diploid sample sizes down to *n* = 10 per population. This follows other work that has shown similar patterns with other haplotype-based statistics for detecting selection (DeGiorgio and Szpiech, 2022; Harris and DeGiorgio, 2020; Harris, et al., 2018; Klassmann and Gautier, 2022). Importantly, this approach now opens up the application of several popular haplotype-based selection statistics (based on extended haplotype homozygosity) to more species where phase information is challenging to know or infer.

For ease of use of these new unphased versions of iHS, nSL, XP-EHH, and XP-nSL, we implement these updates in the latest v2.0 update of the program selscan (Szpiech and Hernandez, 2014), with source code and pre-compiled binaries available at https://www.github.com/szpiech/selscan.

## 5 Methods

### 5.1 Simulations

We evaluate the performance of the phased and unphased versions of iHS, nSL, XP-EHH, and XP-nSL under a generic two-population divergence model using the coalescent simulation program discoal (Kern and Schrider, 2016). We explore five versions of this generic model and name them Demo 1 through Demo 5 (Table S1). Let *N*_0_ and *N*_1_ be the effective population sizes of Population 0 and Population 1 after the split from their ancestral population (of size *N*_*A*_). For Demo 1, we keep a constant population size post-split and let *N*_0_ = *N*_1_= 10,000. For Demo 2, we keep a constant population size post-split and let *N*_0_ = 2 *N*_1_= 10,000. For Demo 3, we keep a constant population size post-split and let 2 *N*_0_ = *N*_1_ = 10,000. For Demo 4, we initially set *N*_0_ = *N*_1_ = 10,000 and let *N*_0_ grow stepwise exponentially every 50 generations starting at 2,000 generations ago until *N*_0_ = 5 *N*_1_= 50,000. For Demo 5, we initially set *N*_0_ = *N*_1_ = 10,000 and let *N*_1_ grow stepwise exponentially every 50 generations starting at 2,000 generations ago until 5 *N*_0_ = *N*_1_ = 50,000.

For each demographic history we vary the population divergence time *t*_9_ ∈ {2000, 4000, 8000} generations ago. For non-neutral simulations, we simulate a sweep in Population 0 in the middle of the simulated region across a range of selection coefficients s ∈ {0.005, 0.01, 0.02}. We vary the frequency at which the adaptive allele starts sweeping as e ∈ {0, 0.01, 0.02, 0.05, 0.10}, where e = 0 indicates a hard sweep and e > 0 indicates a soft sweep, and we also vary the frequency of the selected allele at time of sampling f ∈ {0.7, 0.8, 0.9, 1.0} as well as g ∈ {50, 100} representing fixation of the sweeping allele g generations ago. For all simulations we set the genome length to be L = 500,000 basepairs, the ancestral effective population size to be *N*_*A*_ = 10,000, the per site per generation mutation rate at μ = 2.35 × 10^1 −8^, and the per site per generation recombination rate at r = 1.2 × 10^1−8^. For neutral simulations, we simulate 1,000 replicates for each parameter set, and for non-neutral simulations we simulate 100 replicates for each parameter set. We sample 2*n* ∈ {200, 100, 40, 20} haplotypes, randomly paired together to form *n* ∈ {100, 50, 20, 10} diploid individuals, from each population for analysis. These data sets represent the case where phase is known perfectly. We also create a set of “unphased” data sets from these phased data sets by swapping the alleles of each heterozygote to the opposing haplotype with probability 0.5.

As iHS and nSL are single population statistics, we only analyze Demo 1, Demo 3, and Demo 4 with these statistics, as Demo 2 and Demo 5 have a constant size history identical to Demo 1 for Population 0, where the sweeps are simulated. For XP-EHH and XP-nSL we analyze all five demographic histories.

For all simulations, we compute the relevant statistics (--ihs, --nsl, --xpehh, or --xpnsl) with selscan v2.0 using the --trunc-ok flag. We set --unphased when computing the unphased versions of these statistics, and we do not set it when computing the original phased versions. For iHS and XP-EHH, we also use the --pmap flag to use physical distance instead of a recombination map.

### 5.2 Power and False Positive Rate

Here we evaluate the power and false positive rate for the unphased version of iHS, nSL, XP-EHH, and XP-nSL. For comparison, we also compute the power for the original phased versions of these statistics in two different ways. We compute the phased statistics for a set of simulated datasets where perfect phase is known, and we compute them again for a set of simulated datasets where we destroy phase information (see section 5.1). As the unphased statistics collapse genotypes into derived allele counts, there is no functional difference between these two datasets for these statistics. We compute power in the same way for each statistic regardless of underlying dataset analyzed as described below.

To compute power for iHS and nSL, we follow the approach of Voight, et al. (2006). For these statistics, each non-neutral replicate is individually normalized jointly with all neutral replicates with matching demographic history in 1% allele frequency bins. Because extreme values of the statistic are likely to be clustered along the genome (Voight, et al., 2006), we then compute the proportion of extreme scores (|*iHS*| > 2 or |*nSL*| > 2) within 100kbp non-overlapping windows. We then bin these windows into 10 quantile bins based on the number of scores observed in each window and call the top 1% of these windows as putatively under selection. We calculate the proportion of non-neutral replicates that fall in this top 1% as the power. To compute the false positive rate, we compute the proportion of neutral simulations that fall within the top 1%.

To compute power for XP-EHH and XP-nSL, we follow the approach of (Szpiech, et al., 2021). For these statistics, each non-neutral replicate is individually normalized jointly with all matching neutral replicates. Because extreme values of the statistic are likely to be clustered along the genome (Szpiech, et al., 2021), we then compute the proportion of extreme scores (XP-EHH > 2 or XP-nSL > 2) within 100kbp non-overlapping windows. We then bin these windows into 10 quantile bins based on the number of scores observed in each window and call the top 1% of these windows as putatively under selection. We calculate the proportion of non-neutral replicates that fall in this top 1% as the power. To compute the false positive rate, we compute the proportion of neutral simulations that fall within the top 1%.

## 6 Acknowledgements

This work was supported by the National Institute of General Medical Sciences of the National Institutes of Health under Award Number R35GM146926 and by start-up funds from the Pennsylvania State University’s Department of Biology. Computations for this research were performed using the Pennsylvania State University’s Institute for Computational Data Sciences’ Roar supercomputer.

## Notes

### Competing Interest Statement

The authors have declared no competing interest.

### Summary of Updates

Expanded simulations for lower sample sizes.

