## Supplemental Information for "selscan 2.0: scanning for sweeps in unphased data"

**Table S1**. Demographic history parameters for simulations. $N_A$ represents the ancestral effective population size. $N_0$ represents the effective population size of the population experiencing the sweep. $N_0$ represents the effective population size of the non-sweep population. $t_d$ represents the split time between the two populations.

| | $N_A$ | $N_0$ at split | $N_0$ at present | $N_1$ at split | $N_1$ at present | $t_d$ |
|---|---|---|---|---|---|---|
| Demo 1 | 10,000 | 10,000 | 10,000 | 10,000 | 10,000 | 2,000/4,000/8,000 |
| Demo 2 | 10,000 | 10,000 | 10,000 | 5,000 | 5,000 | 2,000/4,000/8,000 |
| Demo 3 | 10,000 | 5,000 | 5,000 | 10,000 | 10,000 | 2,000/4,000/8,000 |
| Demo 4 | 10,000 | 10,000 | 50,000† | 10,000 | 10,000 | 2,000/4,000/8,000 |
| Demo 5 | 10,000 | 10,000 | 10,000 | 10,000 | 50,000† | 2,000/4,000/8,000 |

†The reached via exponential growth starting 2,000 generations ago.

**Table S2**. False positive rate computed from neutral simulations for varying $t_d$ and demographic history with n = 100 diploid samples (from each population for XP-EHH and XP-nSL).

| | | $t_d = 2000$ | $t_d = 4000$ | $t_d = 8000$ |
|---|---|---|---|---|
| iHS | Demo 1 | 0.013 | 0.1 | 0.009 |
| | Demo 3 | 0.007 | 0.013 | 0.007 |
| | Demo 4 | 0.015 | 0.018 | 0.008 |
| nSL | Demo 1 | 0.01 | 0.015 | 0.008 |
| | Demo 3 | 0.008 | 0.011 | 0.007 |
| | Demo 4 | 0.014 | 0.021 | 0.014 |
| XP-EHH | Demo 1 | 0.013 | 0.013 | 0.016 |
| | Demo 2 | 0.017 | 0.009 | 0.015 |
| | Demo 3 | 0.01 | 0.011 | 0.012 |
| | Demo 4 | 0.012 | 0.014 | 0.014 |
| | Demo 5 | 0.011 | 0.012 | 0.013 |
| XP-nSL | Demo 1 | 0.014 | 0.011 | 0.013 |
| | Demo 2 | 0.019 | 0.011 | 0.012 |
| | Demo 3 | 0.011 | 0.011 | 0.012 |
| | Demo 4 | 0.012 | 0.012 | 0.014 |
| | Demo 5 | 0.011 | 0.012 | 0.014 |

**Table S3**. False positive rate computed from neutral simulations for varying $t_d$ and demographic history with n = 50 diploid samples (from each population for XP-EHH and XP-nSL).

| | | $t_d = 2000$ | $t_d = 4000$ | $t_d = 8000$ |
|---|---|---|---|---|
| iHS | Demo 1 | 0.012 | 0.007 | 0.01 |
| | Demo 3 | 0.015 | 0.009 | 0.009 |
| | Demo 4 | 0.01 | 0.005 | 0.014 |
| nSL | Demo 1 | 0.014 | 0.012 | 0.01 |
| | Demo 3 | 0.01 | 0.012 | 0.011 |
| | Demo 4 | 0.007 | 0.013 | 0.013 |
| XP-EHH | Demo 1 | 0.015 | 0.01 | 0.015 |
| | Demo 2 | 0.018 | 0.009 | 0.013 |
| | Demo 3 | 0.009 | 0.014 | 0.013 |
| | Demo 4 | 0.011 | 0.011 | 0.015 |
| | Demo 5 | 0.012 | 0.016 | 0.02 |
| XP-nSL | Demo 1 | 0.014 | 0.01 | 0.012 |
| | Demo 2 | 0.018 | 0.011 | 0.013 |
| | Demo 3 | 0.009 | 0.014 | 0.014 |
| | Demo 4 | 0.011 | 0.013 | 0.014 |
| | Demo 5 | 0.012 | 0.015 | 0.022 |

**Table S4**. False positive rate computed from neutral simulations for varying $t_d$ and demographic history with n = 20 diploid samples (from each population for XP-EHH and XP-nSL).

| | | $t_d = 2000$ | $t_d = 4000$ | $t_d = 8000$ |
|---|---|---|---|---|
| iHS | Demo 1 | 0.001 | 0.003 | 0.002 |
| | Demo 3 | 0.004 | 0.002 | 0.003 |
| | Demo 4 | 0.002 | 0.0 | 0.003 |
| nSL | Demo 1 | 0.001 | 0.002 | 0.0 |
| | Demo 3 | 0.002 | 0.003 | 0.002 |
| | Demo 4 | 0.0 | 0.0 | 0.002 |
| XP-EHH | Demo 1 | 0.019 | 0.012 | 0.015 |
| | Demo 2 | 0.017 | 0.006 | 0.012 |
| | Demo 3 | 0.011 | 0.013 | 0.013 |
| | Demo 4 | 0.008 | 0.014 | 0.013 |
| | Demo 5 | 0.008 | 0.013 | 0.017 |
| XP-nSL | Demo 1 | 0.018 | 0.014 | 0.014 |
| | Demo 2 | 0.016 | 0.011 | 0.011 |
| | Demo 3 | 0.01 | 0.013 | 0.014 |
| | Demo 4 | 0.009 | 0.013 | 0.013 |
| | Demo 5 | 0.009 | 0.012 | 0.02 |

**Table S5**. False positive rate computed from neutral simulations for varying $t_d$ and demographic history with n = 10 diploid samples (from each population for XP-EHH and XP-nSL).

| | | $t_d = 2000$ | $t_d = 4000$ | $t_d = 8000$ |
|---|---|---|---|---|
| iHS | Demo 1 | 0.0 | 0.0 | 0.0 |
| | Demo 3 | 0.0 | 0.0 | 0.0 |
| | Demo 4 | 0.0 | 0.0 | 0.0 |
| nSL | Demo 1 | 0.0 | 0.0 | 0.0 |
| | Demo 3 | 0.0 | 0.0 | 0.0 |
| | Demo 4 | 0.0 | 0.0 | 0.0 |
| XP-EHH | Demo 1 | 0.014 | 0.013 | 0.014 |
| | Demo 2 | 0.01 | 0.008 | 0.016 |
| | Demo 3 | 0.01 | 0.012 | 0.015 |
| | Demo 4 | 0.01 | 0.014 | 0.012 |
| | Demo 5 | 0.008 | 0.019 | 0.012 |
| XP-nSL | Demo 1 | 0.012 | 0.013 | 0.016 |
| | Demo 2 | 0.011 | 0.006 | 0.013 |
| | Demo 3 | 0.01 | 0.013 | 0.015 |
| | Demo 4 | 0.01 | 0.014 | 0.012 |
| | Demo 5 | 0.009 | 0.02 | 0.01 |

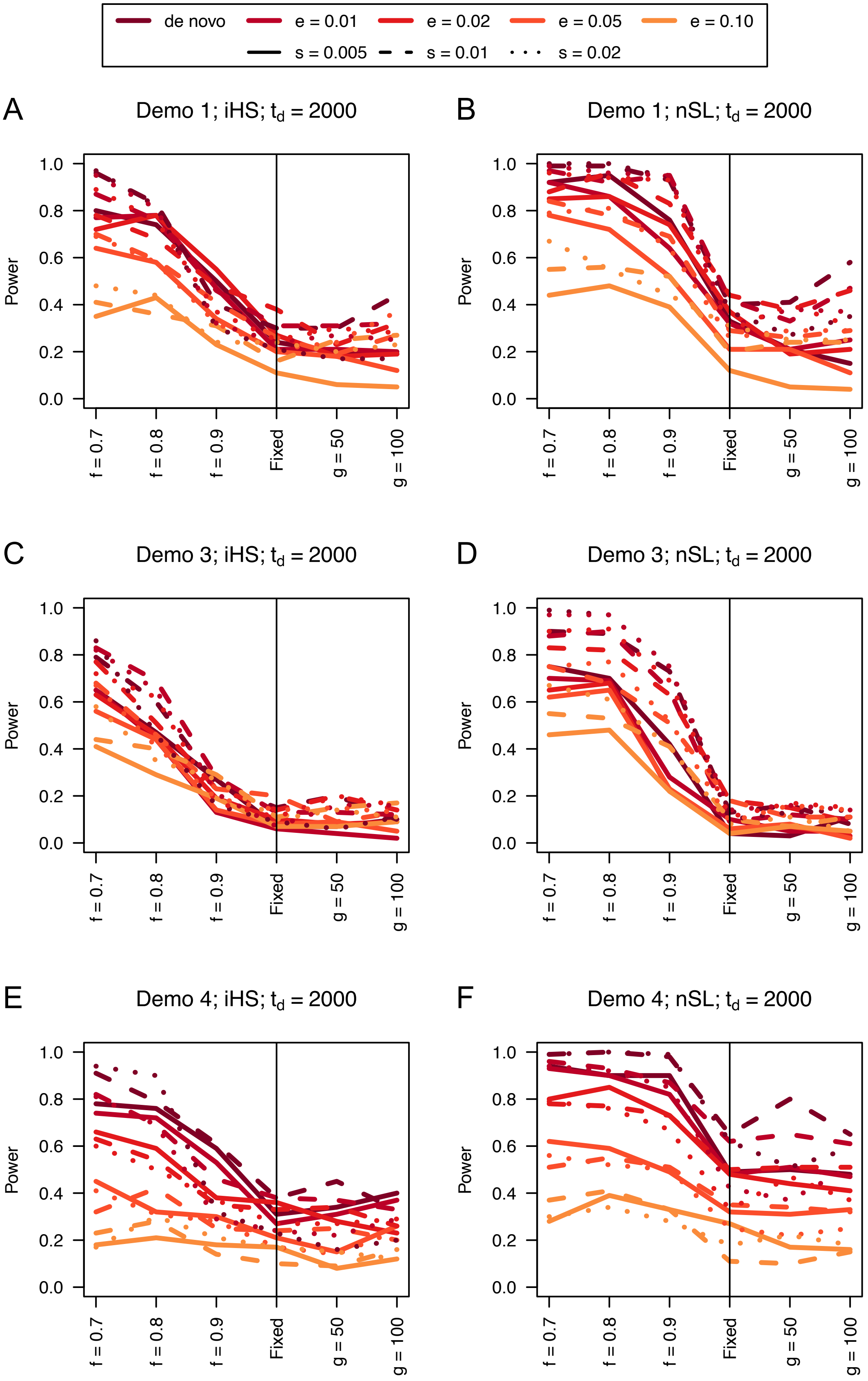

de novo
e = 0.01
e = 0.02
e = 0.05
e = 0.10
s = 0.005
s = 0.01
s = 0.02
A
Demo 1; iHS; $t_d$ = 2000
B
Demo 1; nSL; $t_d$ = 2000
C
Demo 3; iHS; $t_d$ = 2000
D
Demo 3; nSL; $t_d$ = 2000
E
Demo 4; iHS; $t_d$ = 2000
F
Demo 4; nSL; $t_d$ = 2000
Power
f = 0.7
f = 0.8
f = 0.9
Fixed
g = 50
g = 100

**Figure S1**. Power curves for unphased implementations of iHS (A, C, and E) and nSL (B, D, and F) under demographic histories Demo 1 (A and B), Demo 3 (C and D), and Demo 4 (E and F) with n = 100 diploid samples. $s$ is the selection coefficient, $f$ is the frequency of the adaptive allele at time of sampling, $g$ is the number of generations at time of sampling since fixation, $e$ is the frequency at which selection began, and $t_d = 2000$ is the time in generations since the two populations diverged.

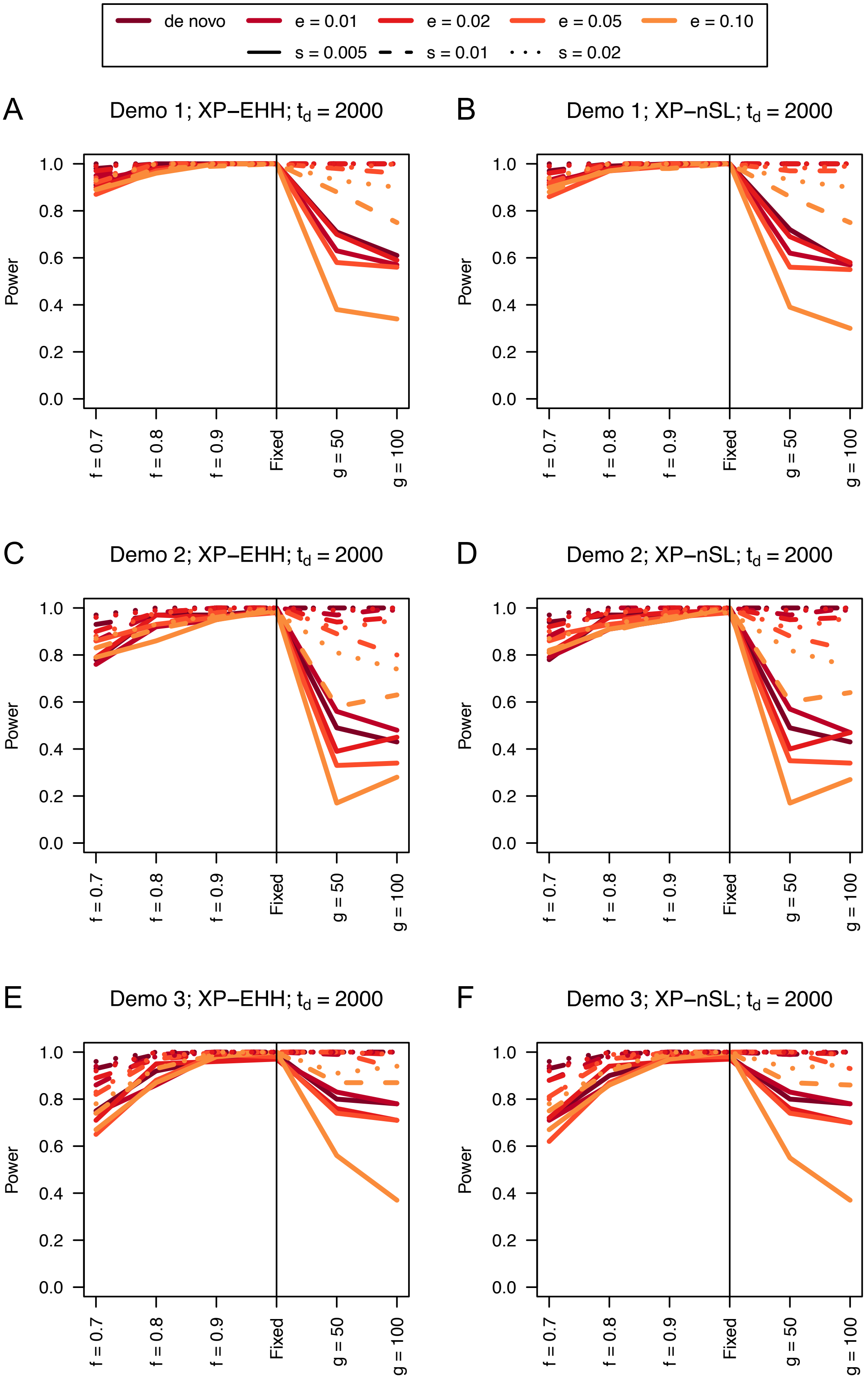
de novo
e = 0.01
e = 0.02
e = 0.05
e = 0.10
s = 0.005
s = 0.01
s = 0.02
A
Demo 1; XP−EHH; $t_d$ = 2000
B
Demo 1; XP−nSL; $t_d$ = 2000
C
Demo 2; XP−EHH; $t_d$ = 2000
D
Demo 2; XP−nSL; $t_d$ = 2000
E
Demo 3; XP−EHH; $t_d$ = 2000
F
Demo 3; XP−nSL; $t_d$ = 2000
Power
0.0
0.2
0.4
0.6
0.8
1.0
f = 0.7
f = 0.8
f = 0.9
Fixed
g = 50
g = 100

**Figure S2**. Power curves for unphased implementations of XP-EHH (A, C, and E) and XP-nSL (B, D, and F) under demographic histories Demo 1 (A and B), Demo 2 (C and D), and Demo 3 (E and F) with n = 100 diploid samples from each population. $s$ is the selection coefficient, $f$ is the frequency of the adaptive allele at time of sampling, $g$ is the number of generations at time of sampling since fixation, $e$ is the frequency at which selection began, and $t_d = 2000$ is the time in generations since the two populations diverged.

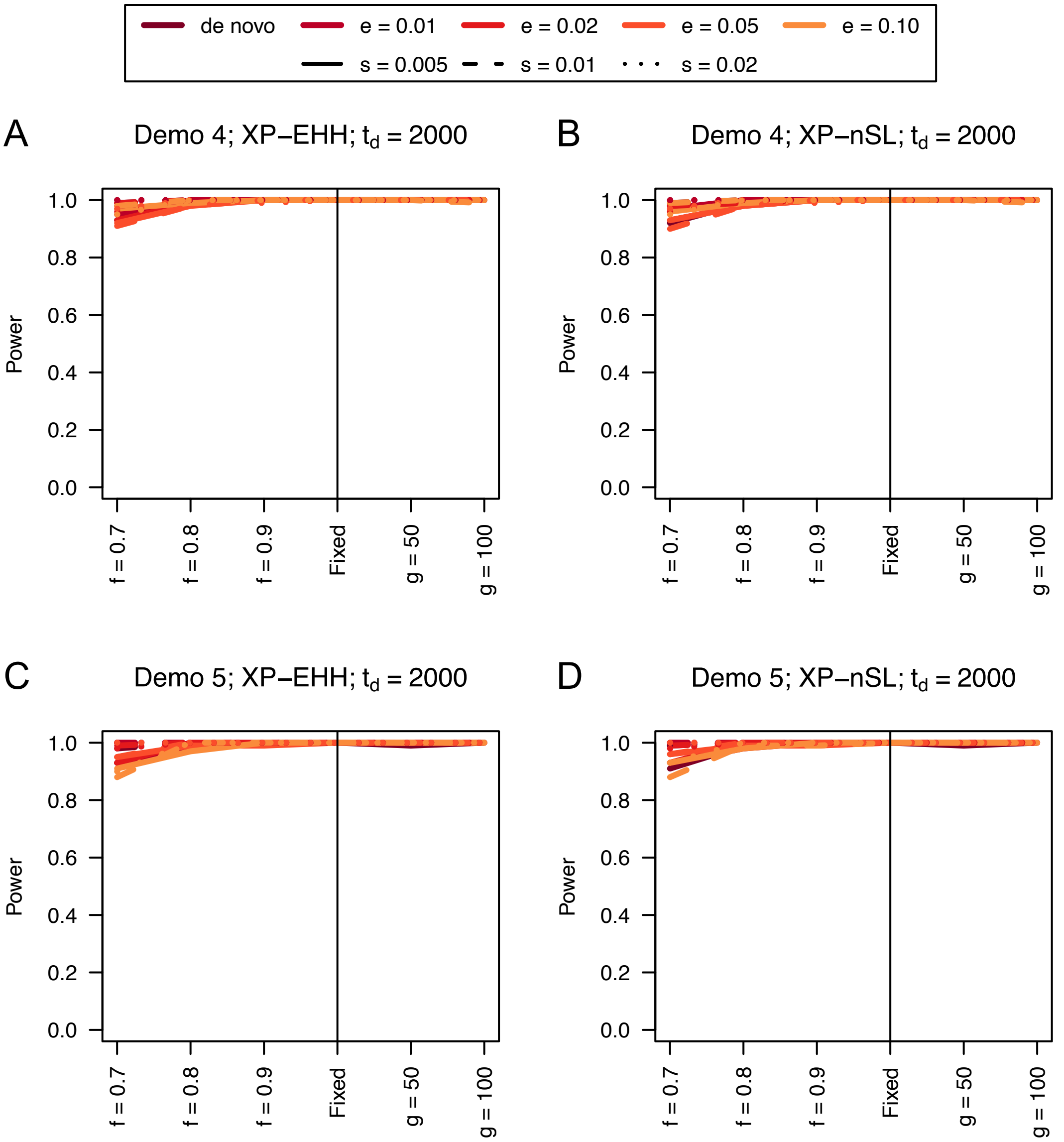


**Figure S3**. Power curves for unphased implementations of XP-EHH (A and C) and XP-nSL (B and D) under demographic histories Demo 4 (A and B), and Demo 5 (C and D) with n = 100 diploid samples from each population. $s$ is the selection coefficient, $f$ is the frequency of the adaptive allele at time of sampling, $g$ is the number of generations at time of sampling since fixation, $e$ is the frequency at which selection began, and $t_d = 2000$ is the time in generations since the two populations diverged.

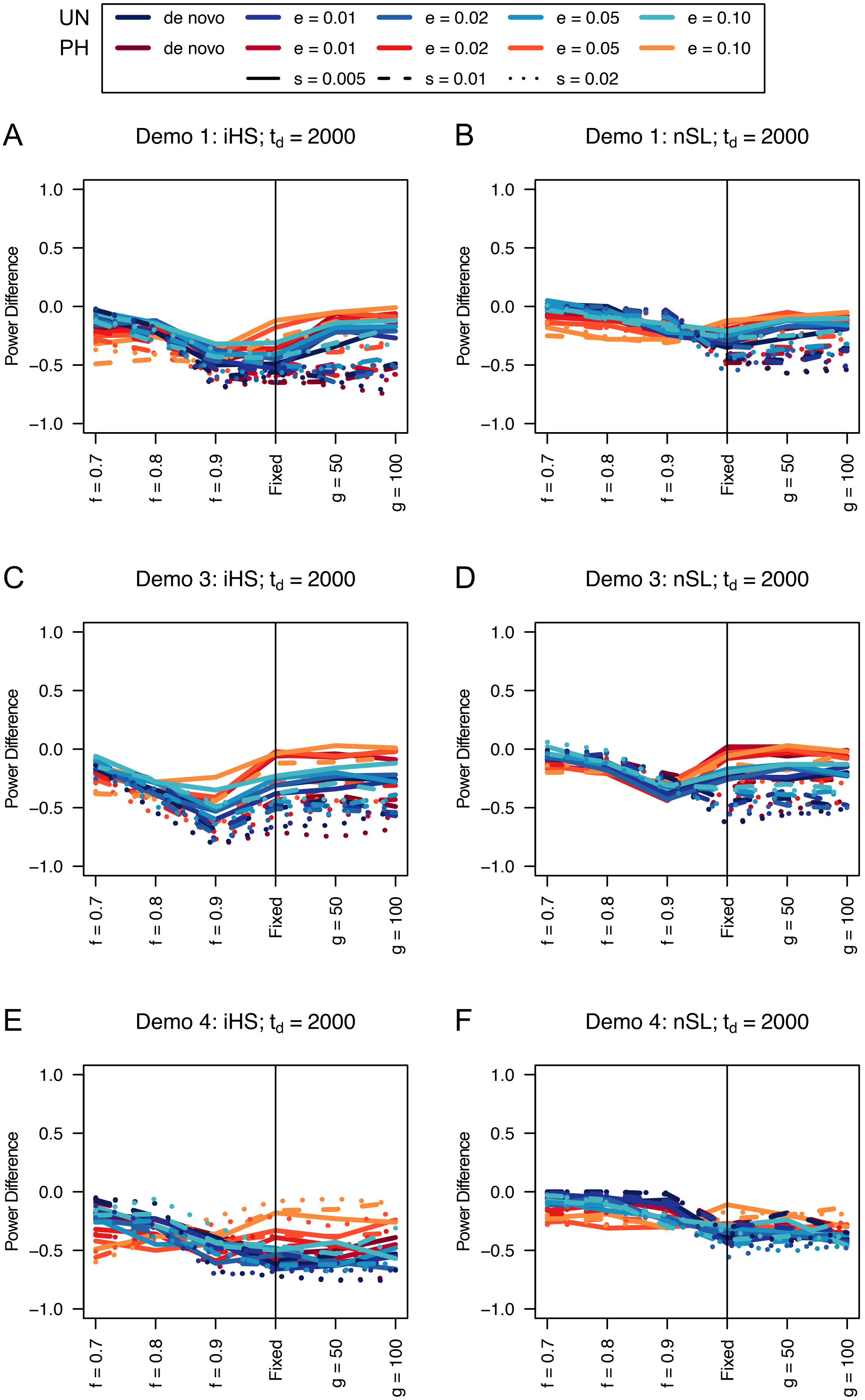
UN
PH
de novo
e = 0.01
e = 0.02
e = 0.05
e = 0.10
s = 0.005
s = 0.01
s = 0.02
A
Demo 1: iHS; $t_d$ = 2000
B
Demo 1: nSL; $t_d$ = 2000
C
Demo 3: iHS; $t_d$ = 2000
D
Demo 3: nSL; $t_d$ = 2000
E
Demo 4: iHS; $t_d$ = 2000
F
Demo 4: nSL; $t_d$ = 2000
Power Difference
1.0
0.5
0.0
−0.5
−1.0
f = 0.7
f = 0.8
f = 0.9
Fixed
g = 50
g = 100

**Figure S4**. Power difference between unphased implementations of iHS (A, C, and E) and nSL (B, D, and F) and phased implementations. Blue curves represent the power difference between the unphased and phased statistics when applied to unphased data (UN). Red curves represent the power difference between the unphased and phased statistics when applied to perfectly phased data (PH). Values greater than 0 indicate the unphased statistic had higher power. Applied to demographic histories Demo 1 (A and B), Demo 3 (C and D), and Demo 4 (E and F) with n = 100 diploid samples. $s$ is the selection coefficient, $f$ is the frequency of the adaptive allele at time of sampling, $g$ is the number of generations at time of sampling since fixation, $e$ is the frequency at which selection began, and $t_d = 2000$ is the time in generations since the two populations diverged.

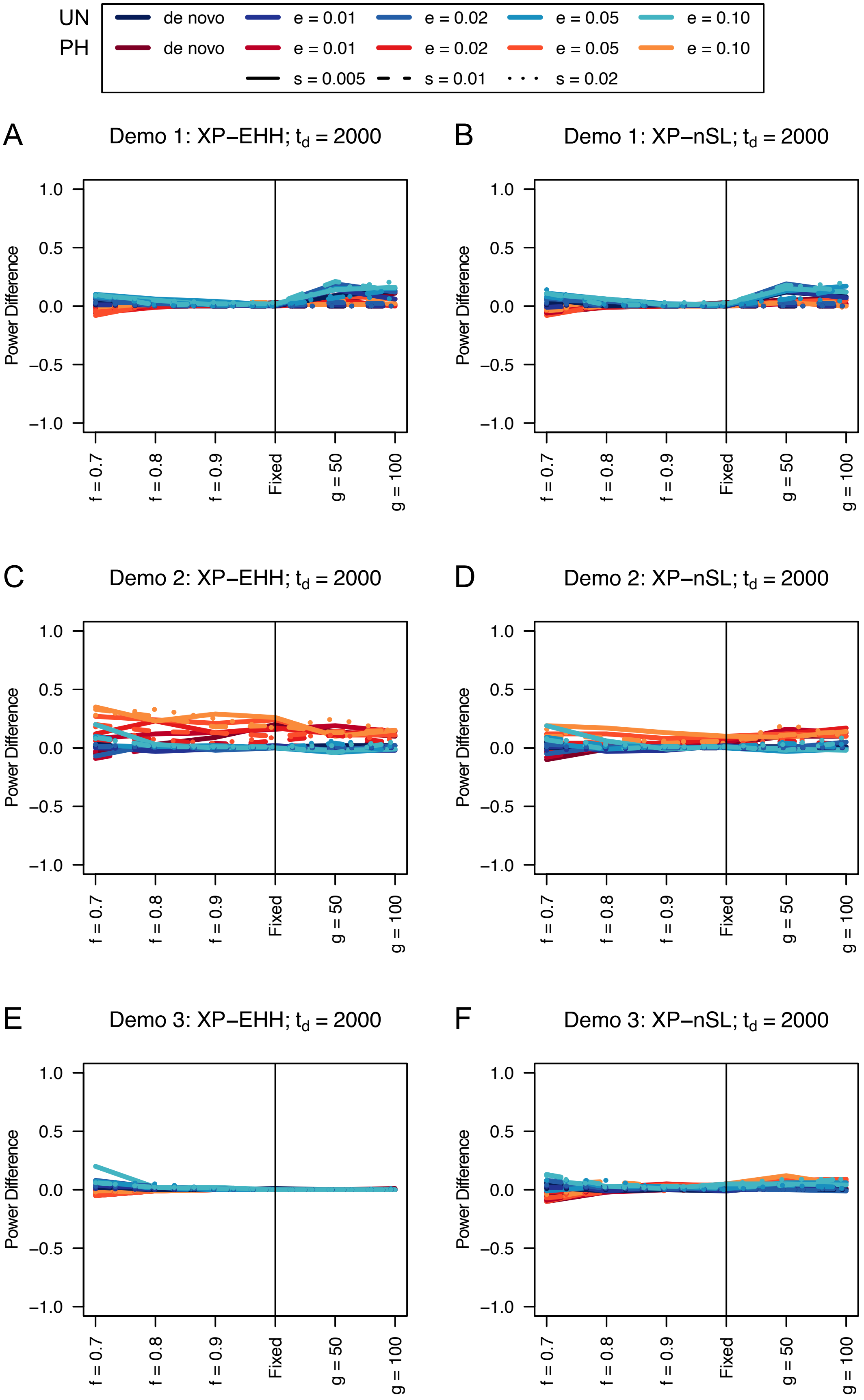
UN
PH
de novo
e = 0.01
e = 0.02
e = 0.05
e = 0.10
s = 0.005
s = 0.01
s = 0.02
A
Demo 1: XP−EHH; $t_d$ = 2000
B
Demo 1: XP−nSL; $t_d$ = 2000
C
Demo 2: XP−EHH; $t_d$ = 2000
D
Demo 2: XP−nSL; $t_d$ = 2000
E
Demo 3: XP−EHH; $t_d$ = 2000
F
Demo 3: XP−nSL; $t_d$ = 2000
Power Difference
1.0
0.5
0.0
−0.5
−1.0
f = 0.7
f = 0.8
f = 0.9
Fixed
g = 50
g = 100

**Figure S5**. Power difference between unphased implementations of XP-EHH (A, C, and E) and XP-nSL (B, D, and F) and phased implementations. Blue curves represent the power difference between the unphased and phased statistics when applied to unphased data (UN). Red curves represent the power difference between the unphased and phased statistics when applied to perfectly phased data (PH). Values greater than 0 indicate the unphased statistic had higher power. Applied to demographic histories Demo 1 (A and B), Demo 2 (C and D), and Demo 3 (E and F) with n = 100 diploid samples from each population. $s$ is the selection coefficient, $f$ is the frequency of the adaptive allele at time of sampling, $g$ is the number of generations at time of sampling since fixation, $e$ is the frequency at which selection began, and $t_d = 2000$ is the time in generations since the two populations diverged.

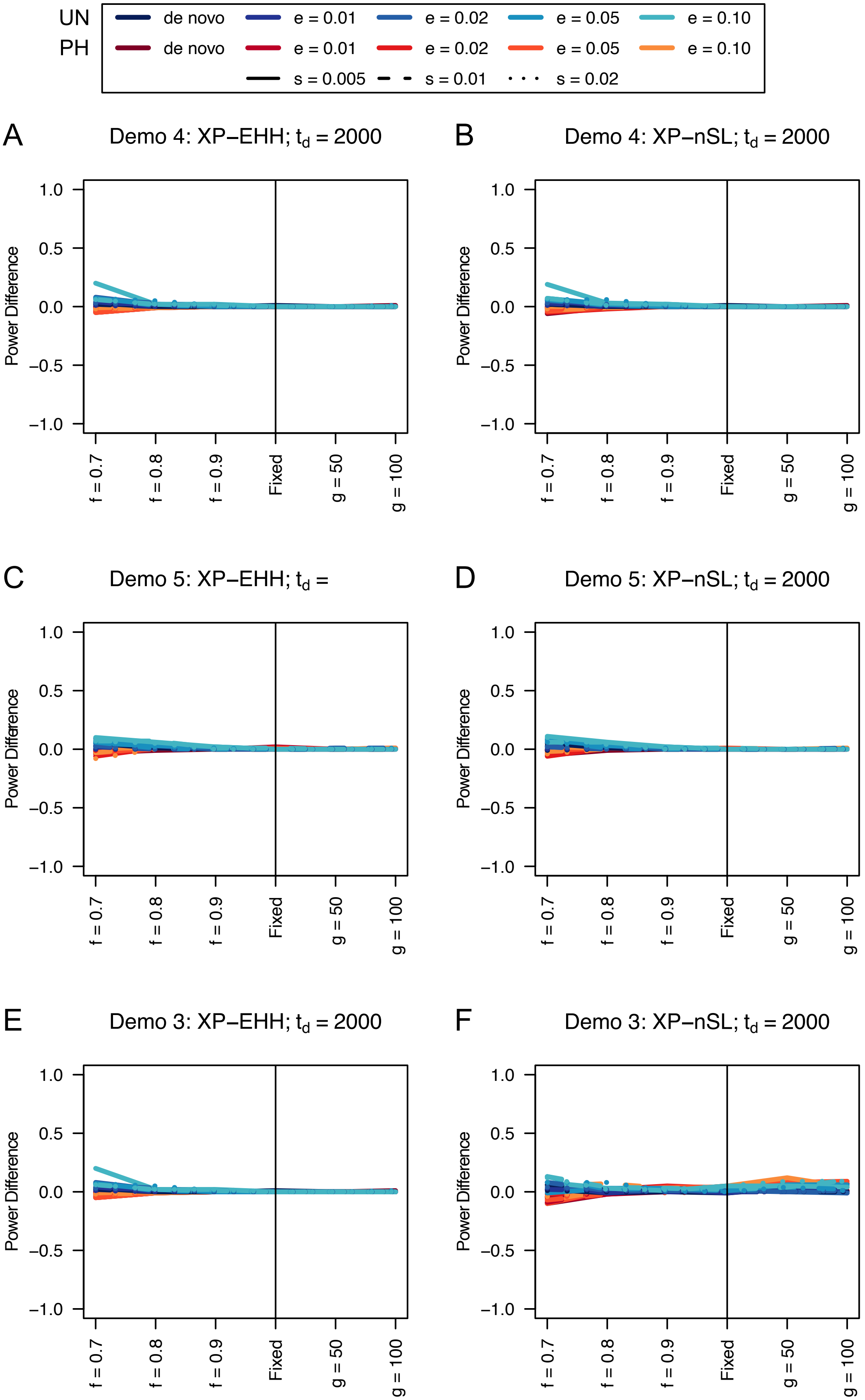

UN de novo e = 0.01 e = 0.02 e = 0.05 e = 0.10
PH de novo e = 0.01 e = 0.02 e = 0.05 e = 0.10
s = 0.005 s = 0.01 s = 0.02
A
Demo 4: XP−EHH; t_d = 2000
B
Demo 4: XP−nSL; t_d = 2000
C
Demo 5: XP−EHH; t_d =
D
Demo 5: XP−nSL; t_d = 2000
E
Demo 3: XP−EHH; t_d = 2000
F
Demo 3: XP−nSL; t_d = 2000
Power Difference
1.0
0.5
0.0
−0.5
−1.0
f = 0.7
f = 0.8
f = 0.9
Fixed
g = 50
g = 100

**Figure S6**. Power difference between unphased implementations of XP-EHH (A and C) and XP-nSL (B and D) and phased implementations. Blue curves represent the power difference between the unphased and phased statistics when applied to unphased data (UN). Red curves represent the power difference between the unphased and phased statistics when applied to perfectly phased data (PH). Values greater than 0 indicate the unphased statistic had higher power. Applied to demographic histories Demo 4 (A and B), and Demo 5 (C and D) with n = 100 diploid samples from each population. $s$ is the selection coefficient, $f$ is the frequency of the adaptive allele at time of sampling, $g$ is the number of generations at time of sampling since fixation, $e$ is the frequency at which selection began, and $t_d = 2000$ is the time in generations since the two populations diverged.

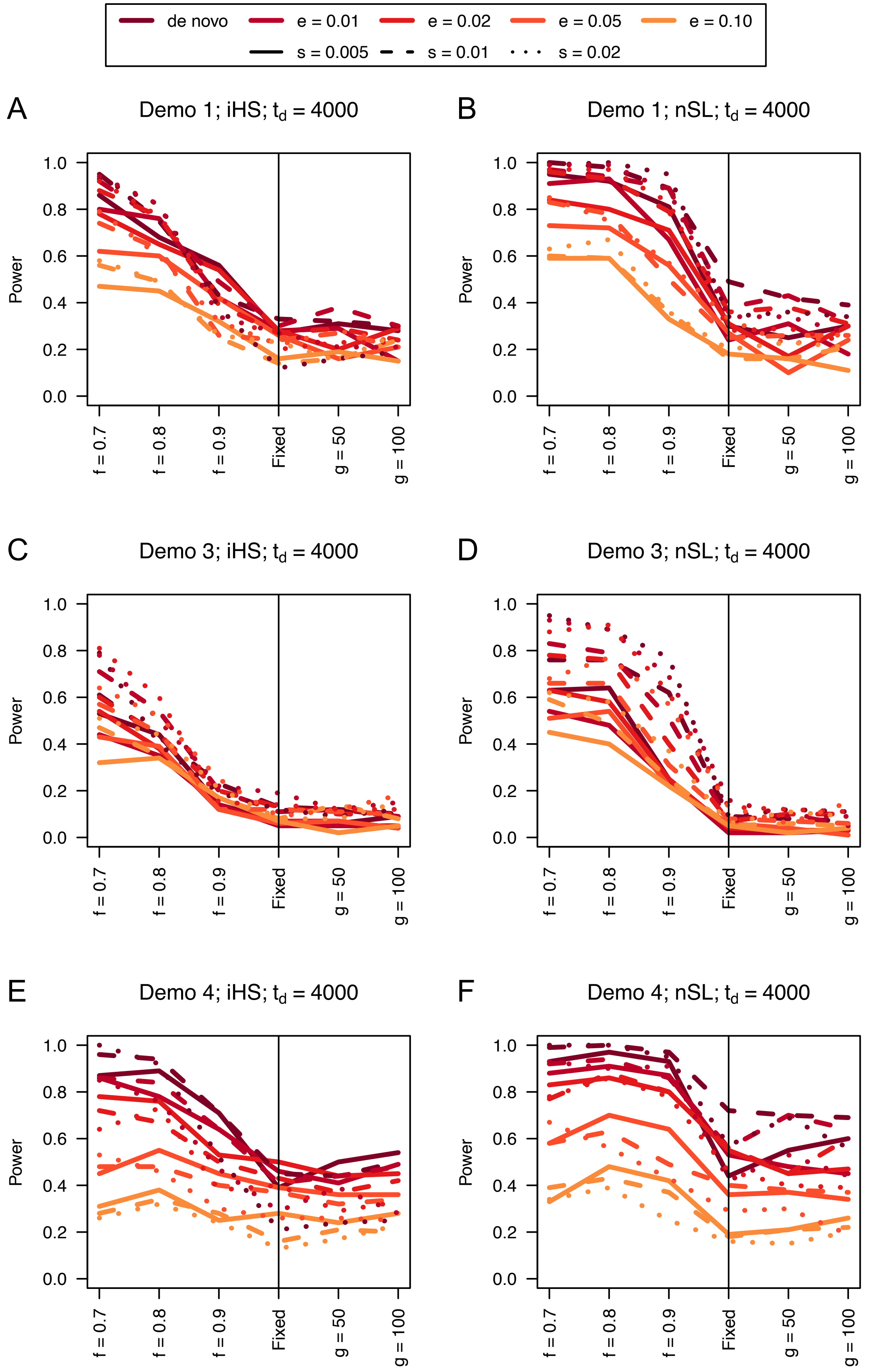
de novo
e = 0.01
e = 0.02
e = 0.05
e = 0.10
s = 0.005
s = 0.01
s = 0.02
A
Demo 1; iHS; t_d = 4000
B
Demo 1; nSL; t_d = 4000
C
Demo 3; iHS; t_d = 4000
D
Demo 3; nSL; t_d = 4000
E
Demo 4; iHS; t_d = 4000
F
Demo 4; nSL; t_d = 4000
Power
0.0
0.2
0.4
0.6
0.8
1.0
f = 0.7
f = 0.8
f = 0.9
Fixed
g = 50
g = 100

**Figure S7**. Power curves for unphased implementations of iHS (A, C, and E) and nSL (B, D, and F) under demographic histories Demo 1 (A and B), Demo 3 (C and D), and Demo 4 (E and F) with n = 100 diploid samples. $s$ is the selection coefficient, $f$ is the frequency of the adaptive allele at time of sampling, $g$ is the number of generations at time of sampling since fixation, $e$ is the frequency at which selection began, and $t_d = 4000$ is the time in generations since the two populations diverged.

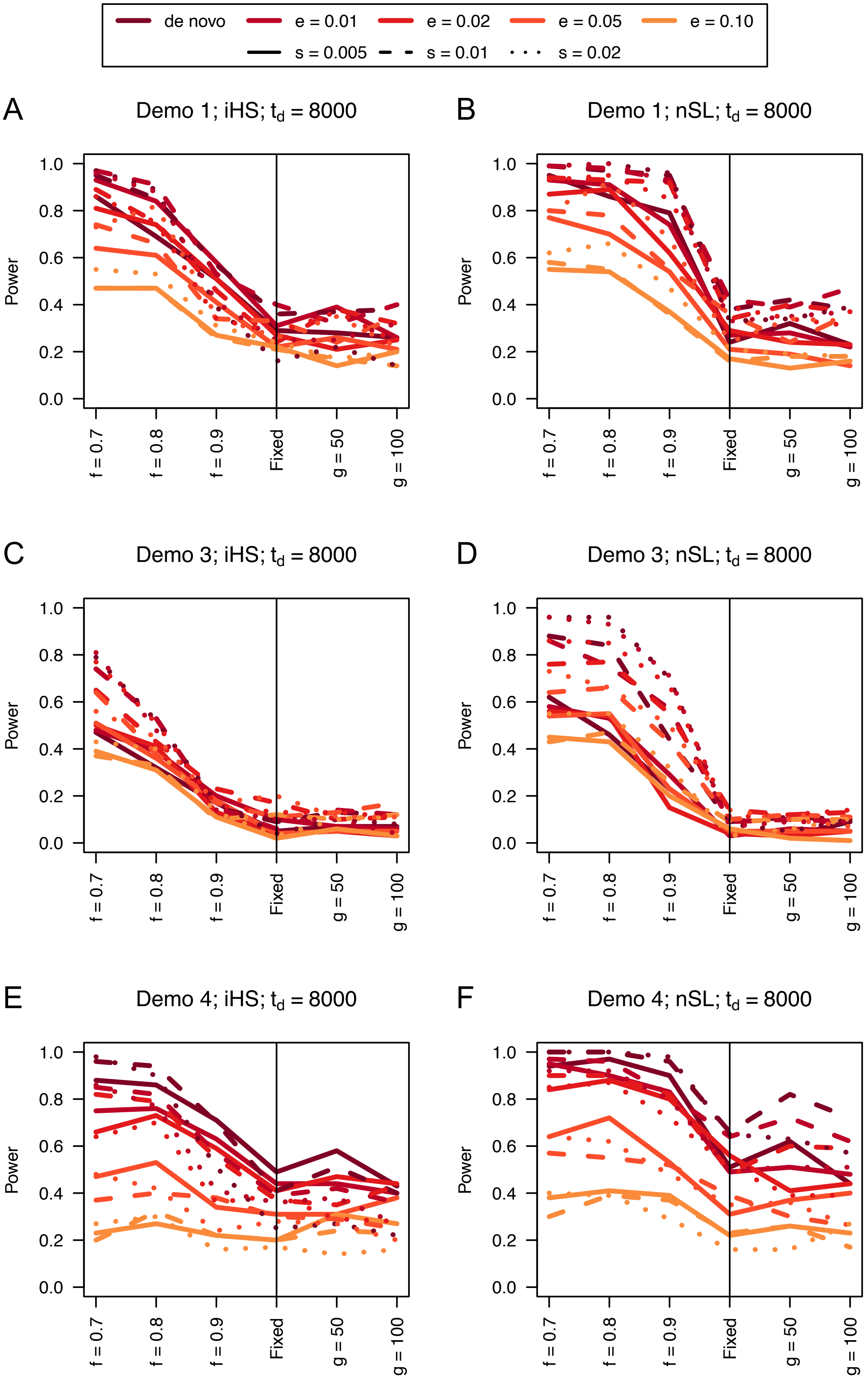

de novo
e = 0.01
e = 0.02
e = 0.05
e = 0.10
s = 0.005
s = 0.01
s = 0.02
A
Demo 1; iHS; $t_d$ = 8000
B
Demo 1; nSL; $t_d$ = 8000
C
Demo 3; iHS; $t_d$ = 8000
D
Demo 3; nSL; $t_d$ = 8000
E
Demo 4; iHS; $t_d$ = 8000
F
Demo 4; nSL; $t_d$ = 8000
Power
0.0
0.2
0.4
0.6
0.8
1.0
f = 0.7
f = 0.8
f = 0.9
Fixed
g = 50
g = 100

**Figure S8**. Power curves for unphased implementations of iHS (A, C, and E) and nSL (B, D, and F) under demographic histories Demo 1 (A and B), Demo 3 (C and D), and Demo 4 (E and F) with n = 100 diploid samples. $s$ is the selection coefficient, $f$ is the frequency of the adaptive allele at time of sampling, $g$ is the number of generations at time of sampling since fixation, $e$ is the frequency at which selection began, and $t_d = 8000$ is the time in generations since the two populations diverged.

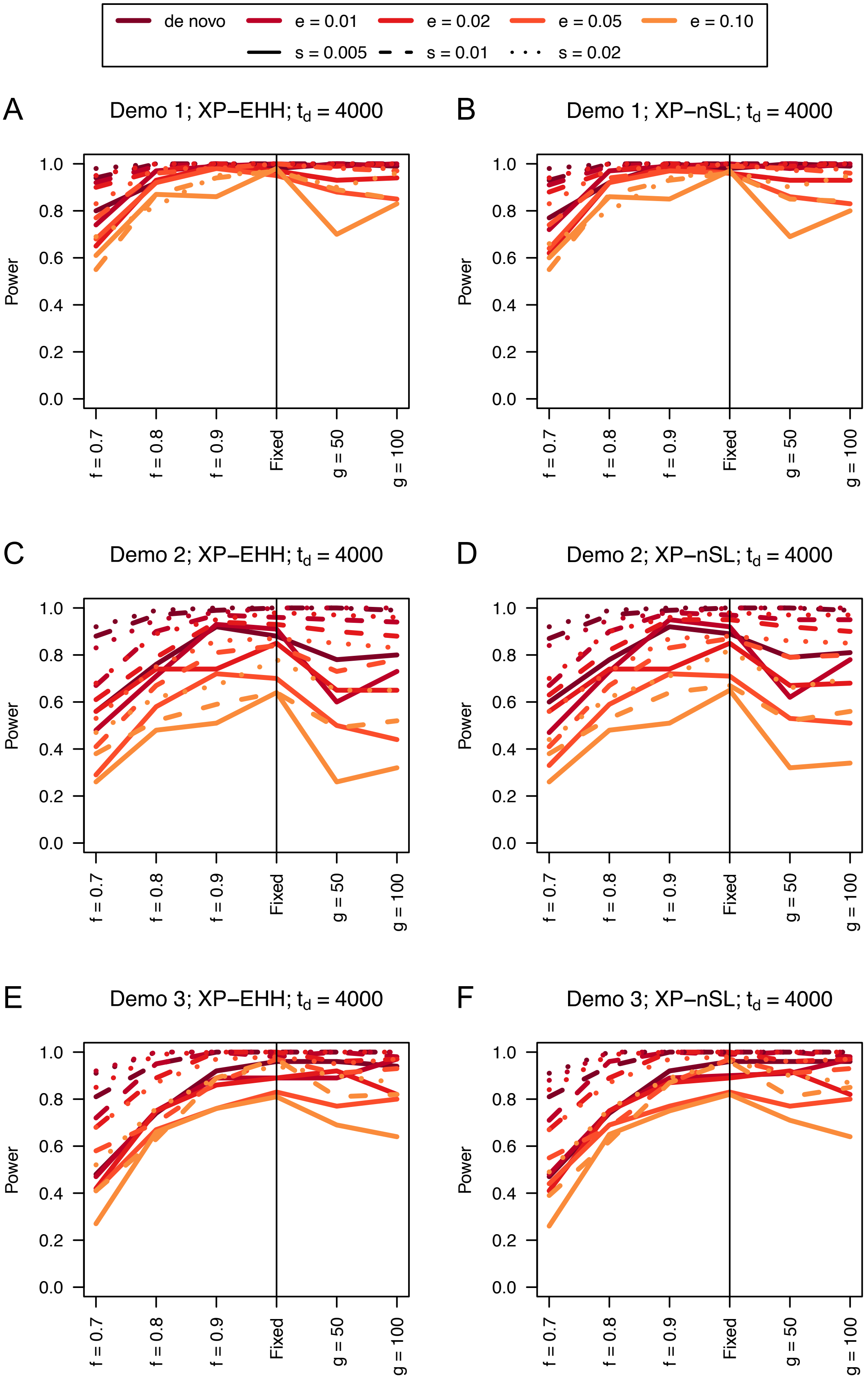

de novo
e = 0.01
e = 0.02
e = 0.05
e = 0.10
s = 0.005
s = 0.01
s = 0.02
A
Demo 1; XP−EHH; $t_d$ = 4000
B
Demo 1; XP−nSL; $t_d$ = 4000
C
Demo 2; XP−EHH; $t_d$ = 4000
D
Demo 2; XP−nSL; $t_d$ = 4000
E
Demo 3; XP−EHH; $t_d$ = 4000
F
Demo 3; XP−nSL; $t_d$ = 4000
Power
0.0
0.2
0.4
0.6
0.8
1.0
f = 0.7
f = 0.8
f = 0.9
Fixed
g = 50
g = 100

**Figure S9**. Power difference between unphased implementations of XP-EHH (A, C, and E) and XP-nSL (B, D, and F) and phased implementations. Blue curves represent the power difference between the unphased and phased statistics when applied to unphased data (UN). Red curves represent the power difference between the unphased and phased statistics when applied to perfectly phased data (PH). Values greater than 0 indicate the unphased statistic had higher power. Applied to demographic histories Demo 1 (A and B), Demo 2 (C and D), and Demo 3 (E and F) with n = 100 diploid samples from each population. $s$ is the selection coefficient, $f$ is the frequency of the adaptive allele at time of sampling, $g$ is the number of generations at time of sampling since fixation, $e$ is the frequency at which selection began, and $t_d = 4000$ is the time in generations since the two populations diverged.

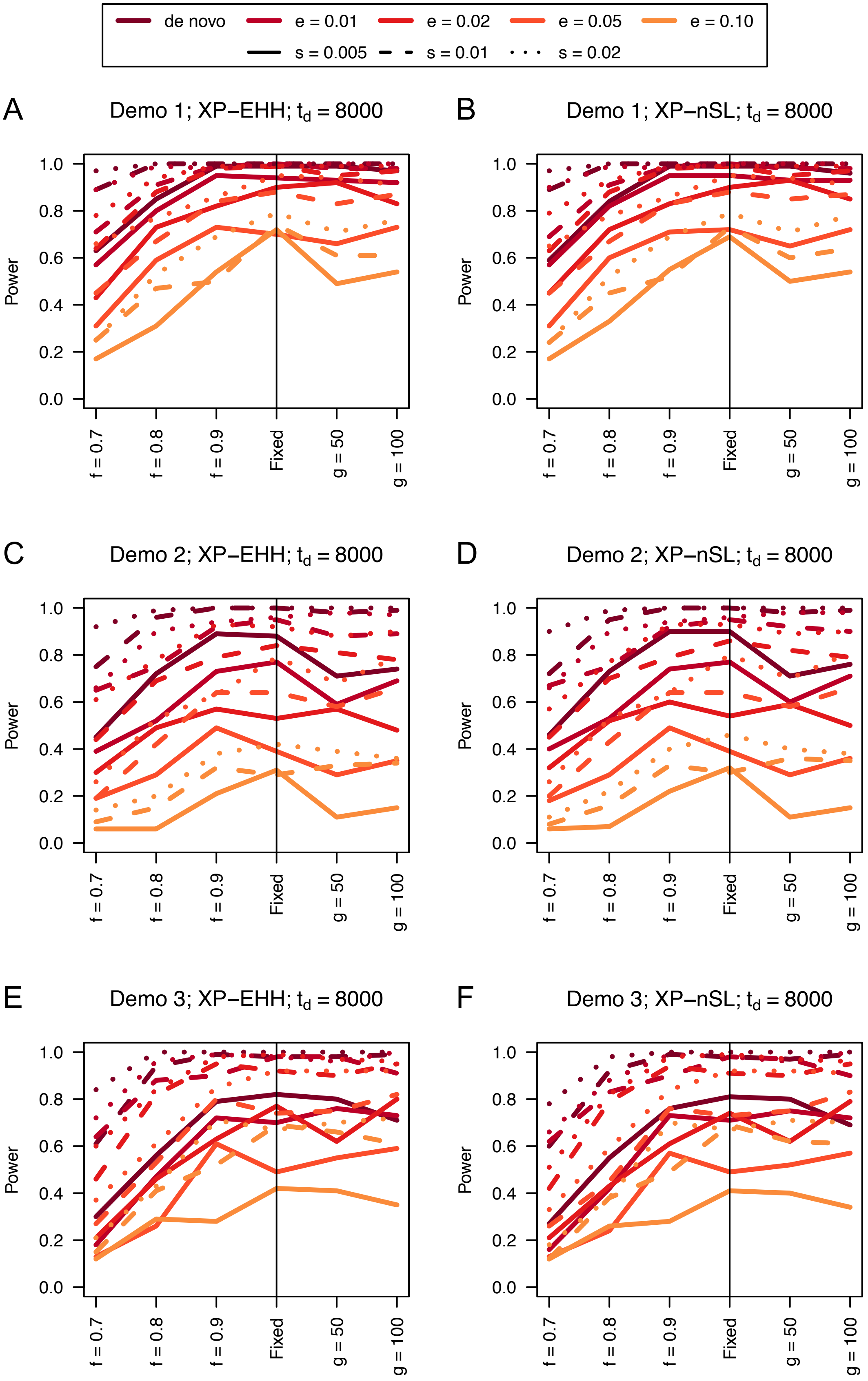
de novo
e = 0.01
e = 0.02
e = 0.05
e = 0.10
s = 0.005
s = 0.01
s = 0.02
A
Demo 1; XP−EHH; $t_d$ = 8000
B
Demo 1; XP−nSL; $t_d$ = 8000
C
Demo 2; XP−EHH; $t_d$ = 8000
D
Demo 2; XP−nSL; $t_d$ = 8000
E
Demo 3; XP−EHH; $t_d$ = 8000
F
Demo 3; XP−nSL; $t_d$ = 8000
Power
0.0
0.2
0.4
0.6
0.8
1.0
f = 0.7
f = 0.8
f = 0.9
Fixed
g = 50
g = 100

**Figure S10**. Power difference between unphased implementations of XP-EHH (A, C, and E) and XP-nSL (B, D, and F) and phased implementations. Blue curves represent the power difference between the unphased and phased statistics when applied to unphased data (UN). Red curves represent the power difference between the unphased and phased statistics when applied to perfectly phased data (PH). Values greater than 0 indicate the unphased statistic had higher power. Applied to demographic histories Demo 1 (A and B), Demo 2 (C and D), and Demo 3 (E and F) with n = 100 diploid samples from each population. $s$ is the selection coefficient, $f$ is the frequency of the adaptive allele at time of sampling, $g$ is the number of generations at time of sampling since fixation, $e$ is the frequency at which selection began, and $t_d = 8000$ is the time in generations since the two populations diverged.

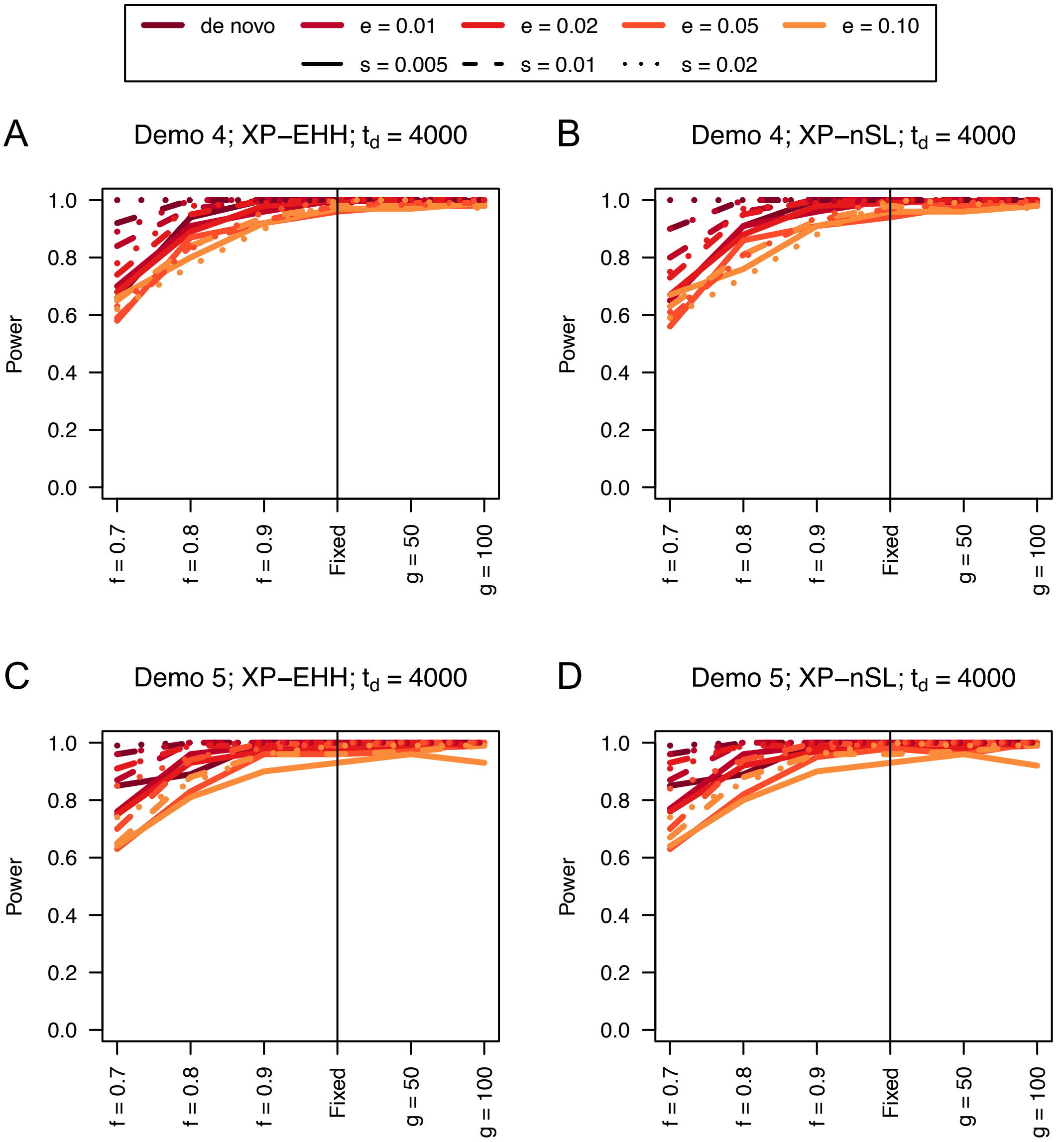


**Figure S11**. Power curves for unphased implementations of XP-EHH (A and C) and XP-nSL (B and D) under demographic histories Demo 4 (A and B), and Demo 5 (C and D) with n = 100 diploid samples from each population. $s$ is the selection coefficient, $f$ is the frequency of the adaptive allele at time of sampling, $g$ is the number of generations at time of sampling since fixation, $e$ is the frequency at which selection began, and $t_d$ = 4000 is the time in generations since the two populations diverged.

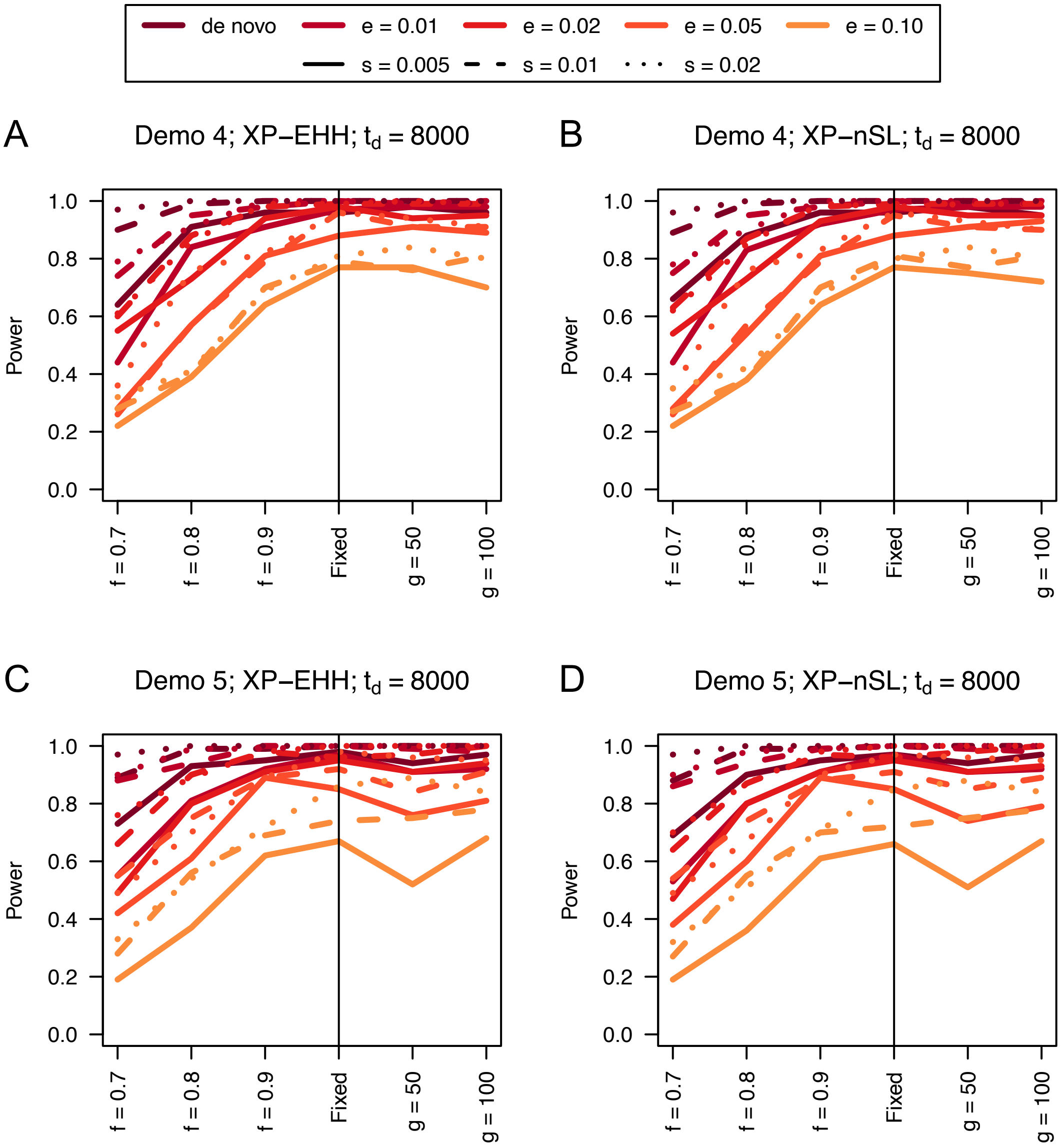


**Figure S12**. Power curves for unphased implementations of XP-EHH (A and C) and XP-nSL (B and D) under demographic histories Demo 4 (A and B), and Demo 5 (C and D) with n = 100 diploid samples from each population. $s$ is the selection coefficient, $f$ is the frequency of the adaptive allele at time of sampling, $g$ is the number of generations at time of sampling since fixation, $e$ is the frequency at which selection began, and $t_d = 8000$ is the time in generations since the two populations diverged.

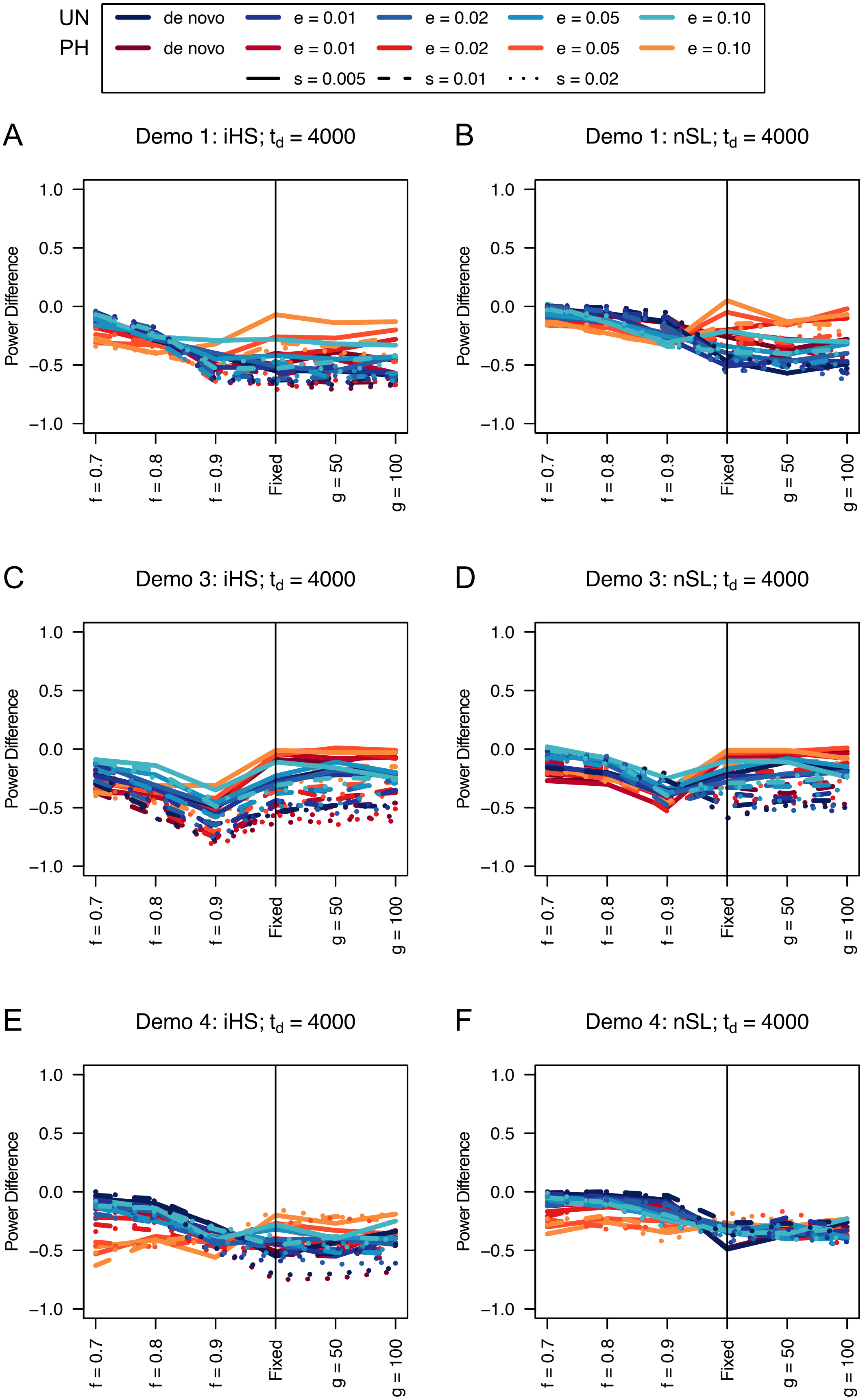
UN
PH
de novo
e = 0.01
e = 0.02
e = 0.05
e = 0.10
de novo
e = 0.01
e = 0.02
e = 0.05
e = 0.10
s = 0.005
s = 0.01
s = 0.02
A
Demo 1: iHS; $t_d$ = 4000
B
Demo 1: nSL; $t_d$ = 4000
C
Demo 3: iHS; $t_d$ = 4000
D
Demo 3: nSL; $t_d$ = 4000
E
Demo 4: iHS; $t_d$ = 4000
F
Demo 4: nSL; $t_d$ = 4000
Power Difference
1.0
0.5
0.0
−0.5
−1.0
f = 0.7
f = 0.8
f = 0.9
Fixed
g = 50
g = 100

**Figure S13**. Power difference between unphased implementations of iHS (A, C, and E) and nSL (B, D, and F) and phased implementations. Blue curves represent the power difference between the unphased and phased statistics when applied to unphased data (UN). Red curves represent the power difference between the unphased and phased statistics when applied to perfectly phased data (PH). Values greater than 0 indicate the unphased statistic had higher power. Applied to demographic histories Demo 1 (A and B), Demo 3 (C and D), and Demo 4 (E and F) with n = 100 diploid samples. $s$ is the selection coefficient, $f$ is the frequency of the adaptive allele at time of sampling, $g$ is the number of generations at time of sampling since fixation, $e$ is the frequency at which selection began, and $t_d = 4000$ is the time in generations since the two populations diverged.

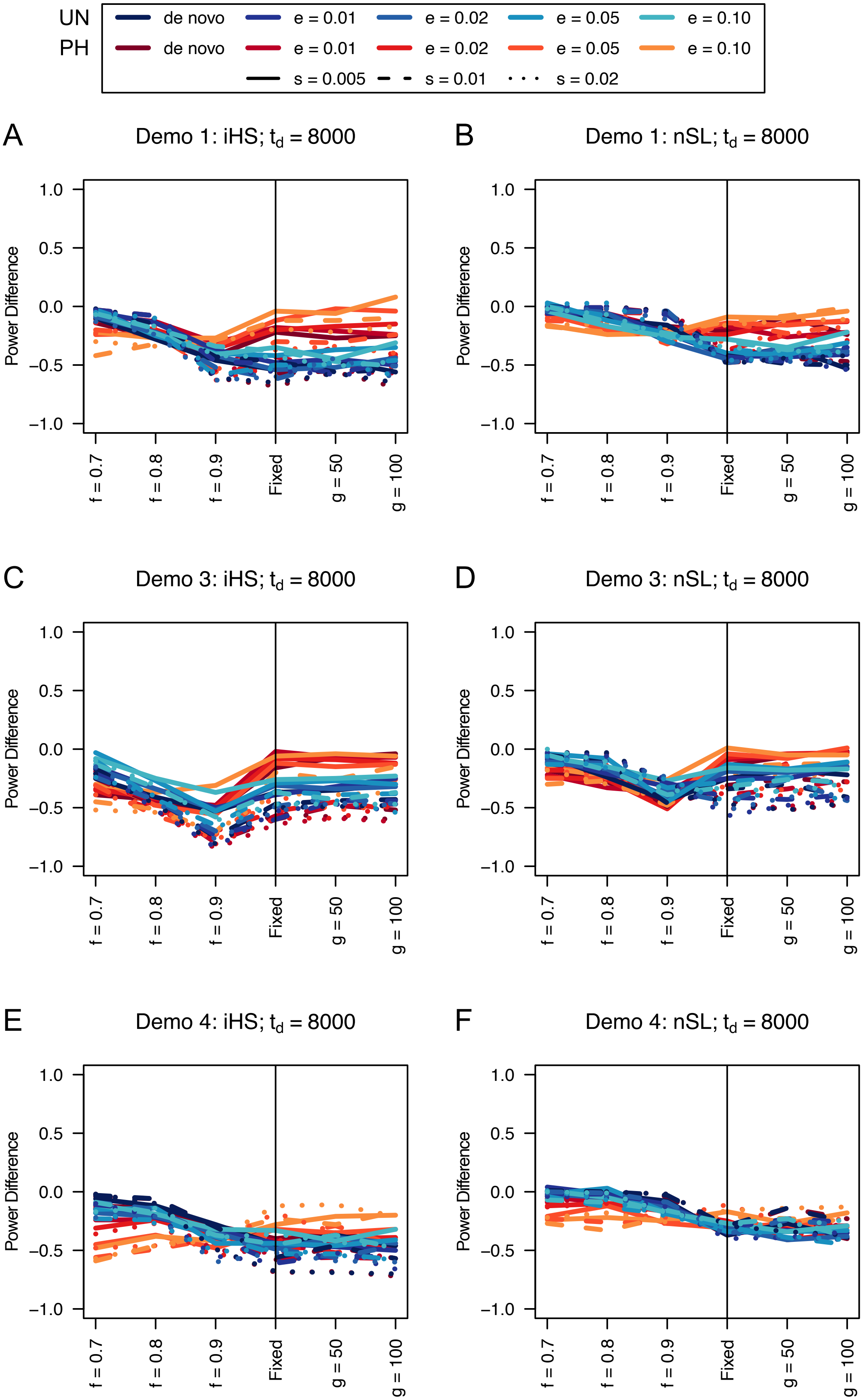

UN
PH
de novo
e = 0.01
e = 0.02
e = 0.05
e = 0.10
s = 0.005
s = 0.01
s = 0.02
A
Demo 1: iHS; $t_d$ = 8000
B
Demo 1: nSL; $t_d$ = 8000
C
Demo 3: iHS; $t_d$ = 8000
D
Demo 3: nSL; $t_d$ = 8000
E
Demo 4: iHS; $t_d$ = 8000
F
Demo 4: nSL; $t_d$ = 8000
Power Difference
1.0
0.5
0.0
−0.5
−1.0
f = 0.7
f = 0.8
f = 0.9
Fixed
g = 50
g = 100

**Figure S14**. Power difference between unphased implementations of iHS (A, C, and E) and nSL (B, D, and F) and phased implementations. Blue curves represent the power difference between the unphased and phased statistics when applied to unphased data (UN). Red curves represent the power difference between the unphased and phased statistics when applied to perfectly phased data (PH). Values greater than 0 indicate the unphased statistic had higher power. Applied to demographic histories Demo 1 (A and B), Demo 3 (C and D), and Demo 4 (E and F) with n = 100 diploid samples. $s$ is the selection coefficient, $f$ is the frequency of the adaptive allele at time of sampling, $g$ is the number of generations at time of sampling since fixation, $e$ is the frequency at which selection began, and $t_d = 8000$ is the time in generations since the two populations diverged.

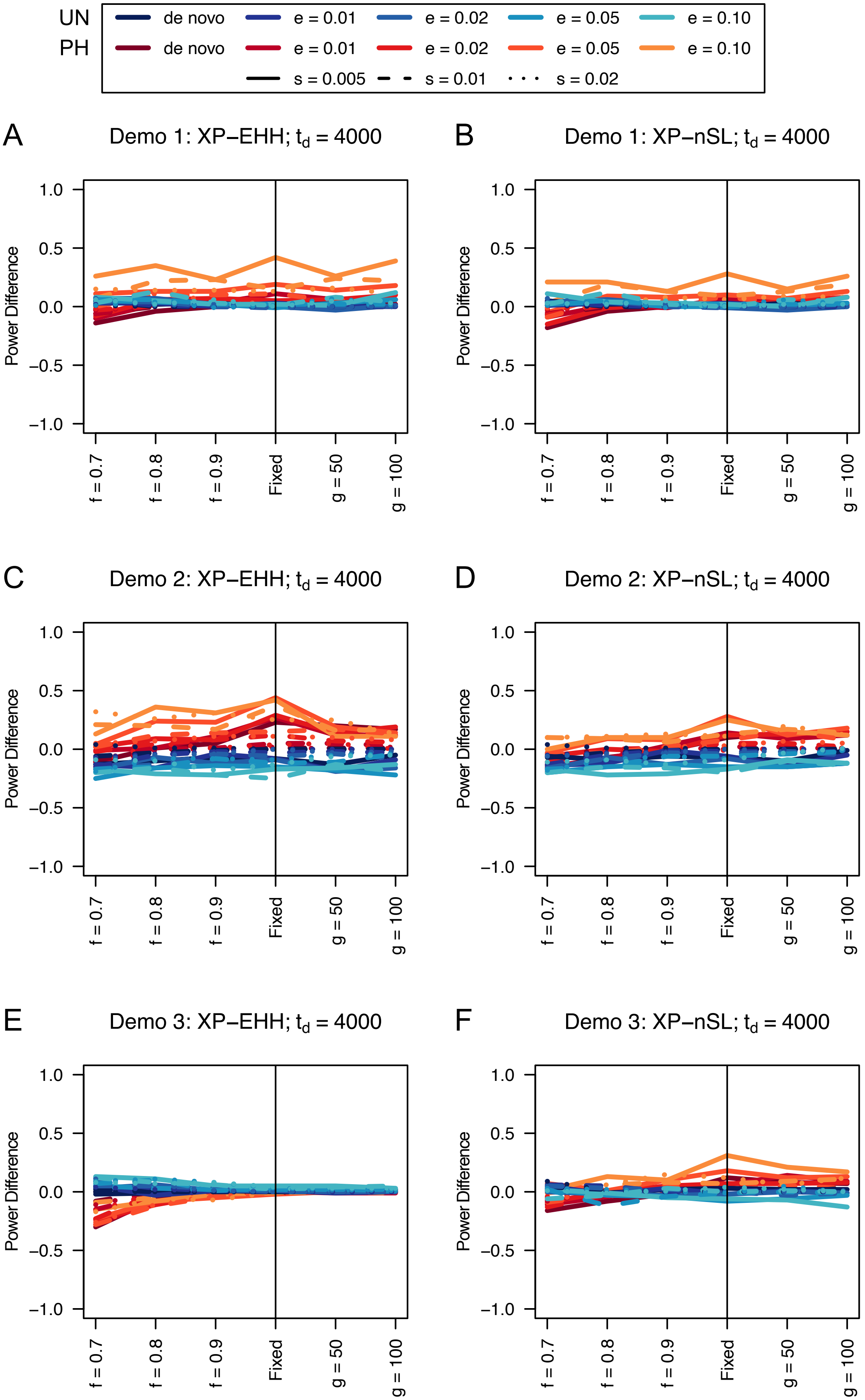

UN
PH
de novo
e = 0.01
e = 0.02
e = 0.05
e = 0.10
de novo
e = 0.01
e = 0.02
e = 0.05
e = 0.10
s = 0.005
s = 0.01
s = 0.02
A
Demo 1: XP−EHH; $t_d$ = 4000
B
Demo 1: XP−nSL; $t_d$ = 4000
C
Demo 2: XP−EHH; $t_d$ = 4000
D
Demo 2: XP−nSL; $t_d$ = 4000
E
Demo 3: XP−EHH; $t_d$ = 4000
F
Demo 3: XP−nSL; $t_d$ = 4000
Power Difference
1.0
0.5
0.0
−0.5
−1.0
f = 0.7
f = 0.8
f = 0.9
Fixed
g = 50
g = 100

**Figure S15**. Power difference between unphased implementations of XP-EHH (A, C, and E) and XP-nSL (B, D, and F) and phased implementations. Blue curves represent the power difference between the unphased and phased statistics when applied to unphased data (UN). Red curves represent the power difference between the unphased and phased statistics when applied to perfectly phased data (PH). Values greater than 0 indicate the unphased statistic had higher power. Applied to demographic histories Demo 1 (A and B), Demo 2 (C and D), and Demo 3 (E and F) with n = 100 diploid samples from each population. $s$ is the selection coefficient, $f$ is the frequency of the adaptive allele at time of sampling, $g$ is the number of generations at time of sampling since fixation, $e$ is the frequency at which selection began, and $t_d = 4000$ is the time in generations since the two populations diverged.

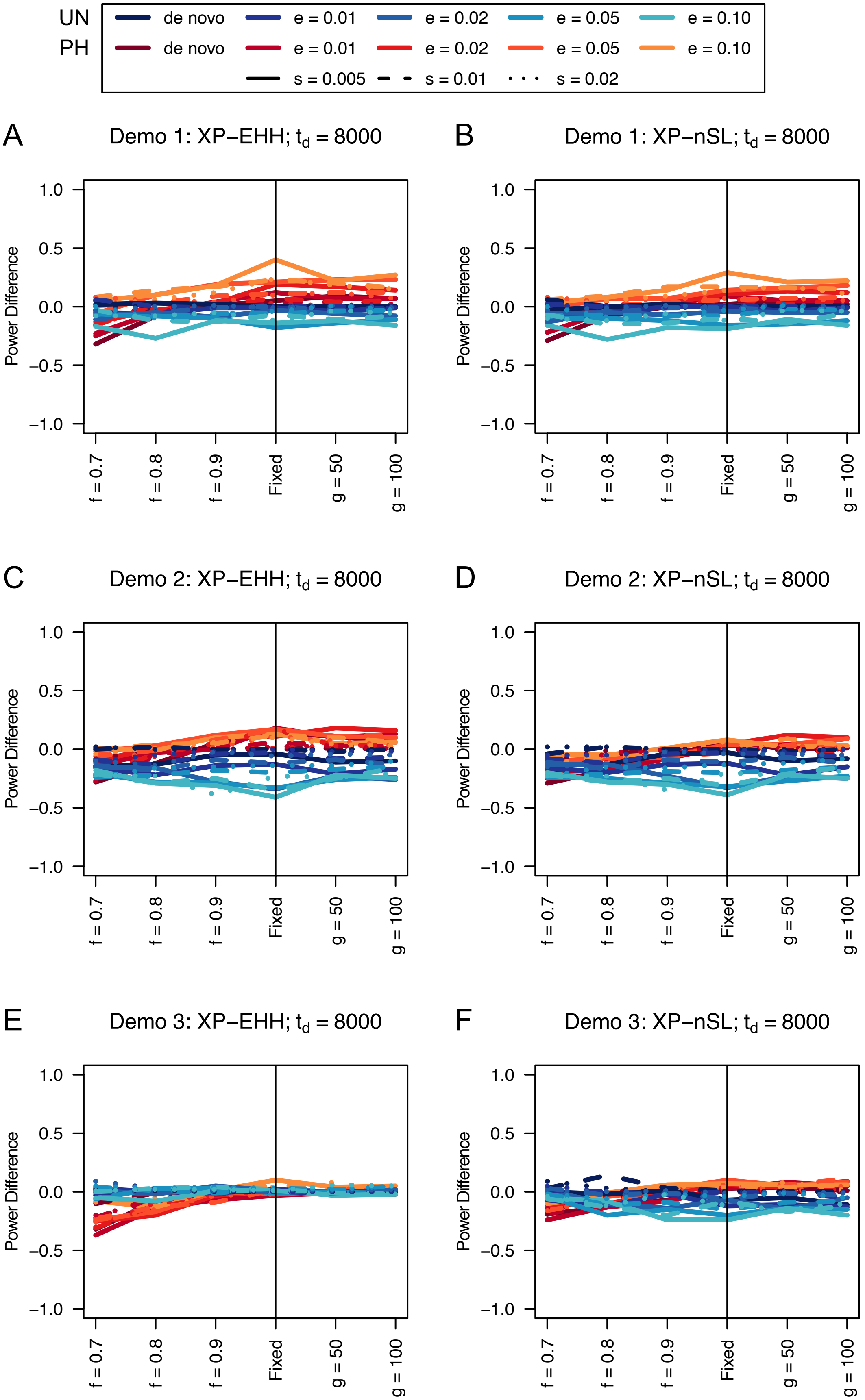

UN
PH
de novo
e = 0.01
e = 0.02
e = 0.05
e = 0.10
s = 0.005
s = 0.01
s = 0.02
A
Demo 1: XP−EHH; $t_d$ = 8000
B
Demo 1: XP−nSL; $t_d$ = 8000
C
Demo 2: XP−EHH; $t_d$ = 8000
D
Demo 2: XP−nSL; $t_d$ = 8000
E
Demo 3: XP−EHH; $t_d$ = 8000
F
Demo 3: XP−nSL; $t_d$ = 8000
Power Difference
1.0
0.5
0.0
−0.5
−1.0
f = 0.7
f = 0.8
f = 0.9
Fixed
g = 50
g = 100

**Figure S16**. Power difference between unphased implementations of XP-EHH (A, C, and E) and XP-nSL (B, D, and F) and phased implementations. Blue curves represent the power difference between the unphased and phased statistics when applied to unphased data (UN). Red curves represent the power difference between the unphased and phased statistics when applied to perfectly phased data (PH). Values greater than 0 indicate the unphased statistic had higher power. Applied to demographic histories Demo 1 (A and B), Demo 2 (C and D), and Demo 3 (E and F) with n = 100 diploid samples from each population. $s$ is the selection coefficient, $f$ is the frequency of the adaptive allele at time of sampling, $g$ is the number of generations at time of sampling since fixation, $e$ is the frequency at which selection began, and $t_d = 8000$ is the time in generations since the two populations diverged.

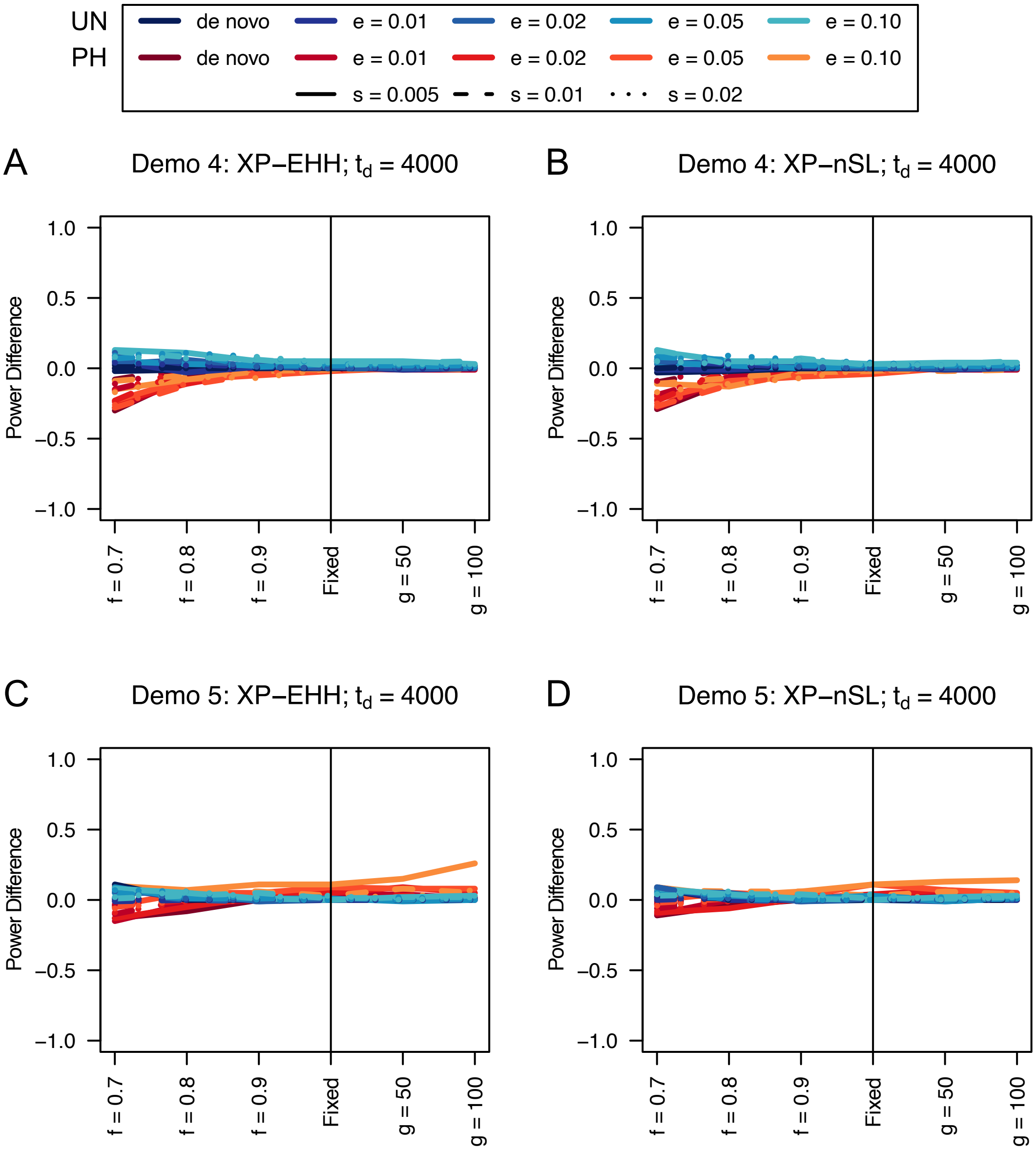


**Figure S17**. Power difference between unphased implementations of XP-EHH (A and C) and XP-nSL (B and D) and phased implementations. Blue curves represent the power difference between the unphased and phased statistics when applied to unphased data (UN). Red curves represent the power difference between the unphased and phased statistics when applied to perfectly phased data (PH). Values greater than 0 indicate the unphased statistic had higher power. Applied to demographic histories Demo 4 (A and B), and Demo 5 (C and D) with n = 100 diploid samples from each population. $s$ is the selection coefficient, $f$ is the frequency of the adaptive allele at time of sampling, $g$ is the number of generations at time of sampling since fixation, $e$ is the frequency at which selection began, and $t_d = 4000$ is the time in generations since the two populations diverged.

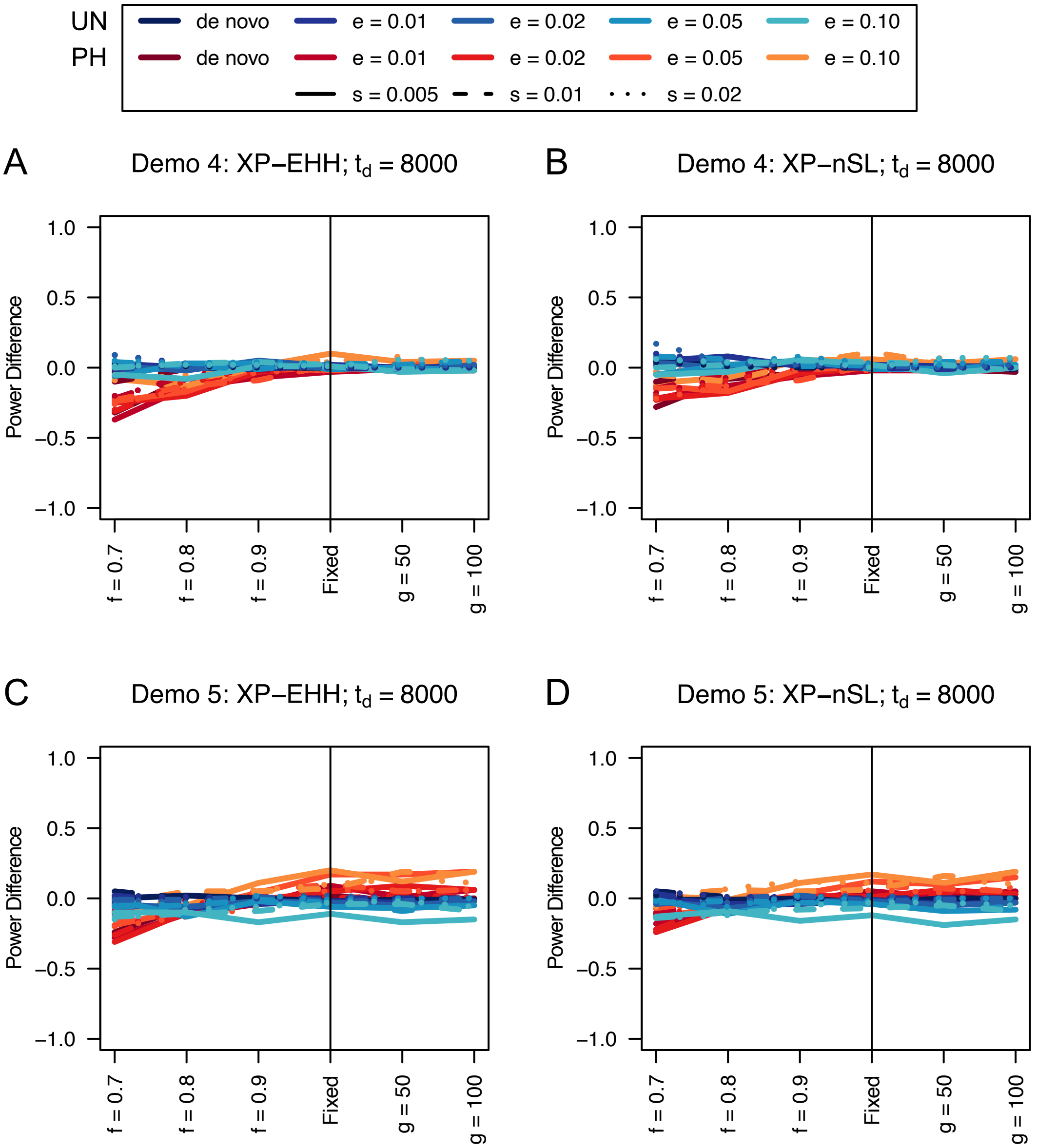


**Table S18**. Power difference between unphased implementations of XP-EHH (A and C) and XP-nSL (B and D) and phased implementations. Blue curves represent the power difference between the unphased and phased statistics when applied to unphased data (UN). Red curves represent the power difference between the unphased and phased statistics when applied to perfectly phased data (PH). Values greater than 0 indicate the unphased statistic had higher power. Applied to demographic histories Demo 4 (A and B), and Demo 5 (C and D) with n = 100 diploid samples from each population. $s$ is the selection coefficient, $f$ is the frequency of the adaptive allele at time of sampling, $g$ is the number of generations at time of sampling since fixation, $e$ is the frequency at which selection began, and $t_d$ = 8000 is the time in generations since the two populations diverged.

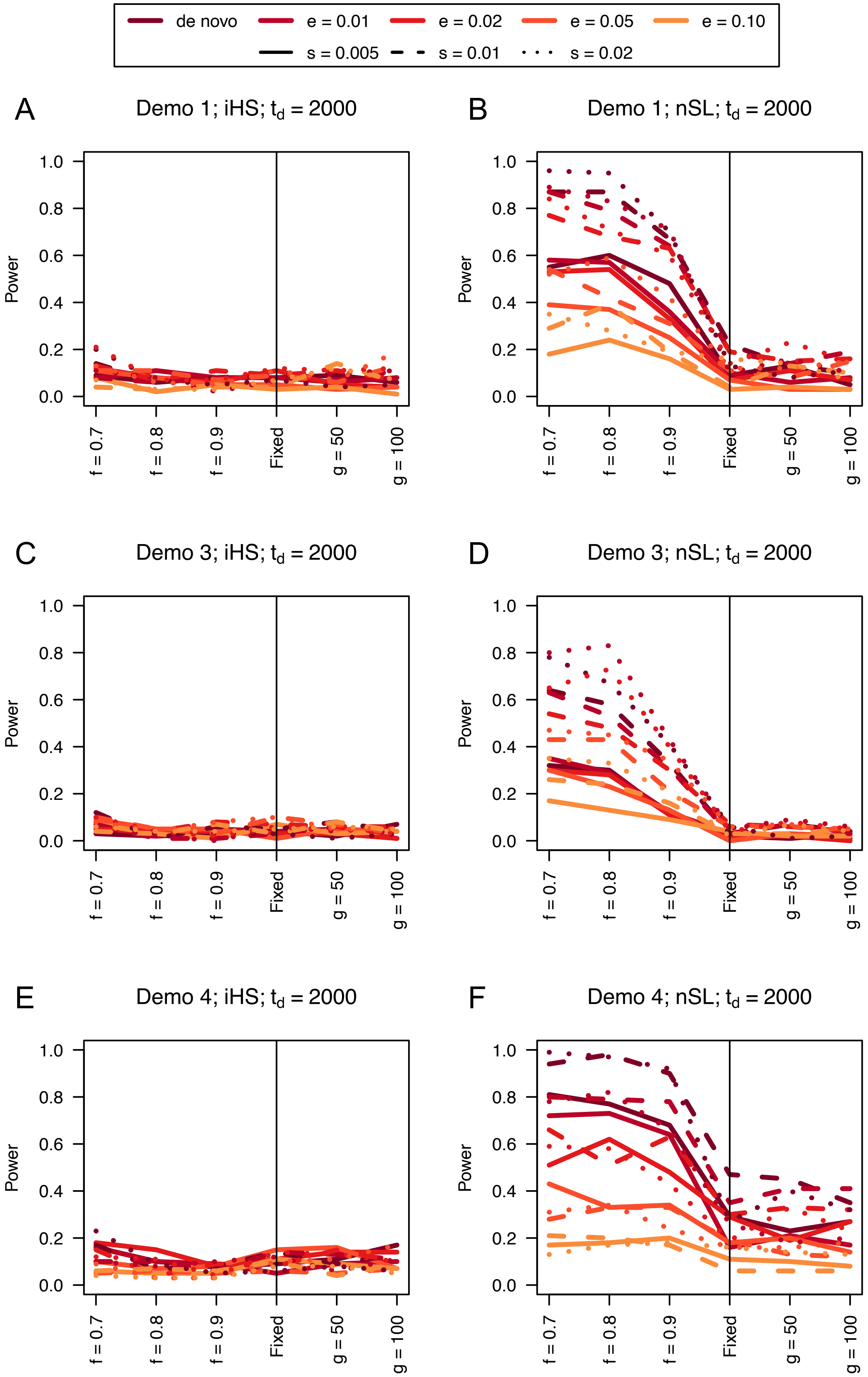

de novo
e = 0.01
e = 0.02
e = 0.05
e = 0.10
s = 0.005
s = 0.01
s = 0.02
A
Demo 1; iHS; t_d = 2000
B
Demo 1; nSL; t_d = 2000
C
Demo 3; iHS; t_d = 2000
D
Demo 3; nSL; t_d = 2000
E
Demo 4; iHS; t_d = 2000
F
Demo 4; nSL; t_d = 2000
Power
f = 0.7
f = 0.8
f = 0.9
Fixed
g = 50
g = 100

**Figure S19**. Power curves for unphased implementations of iHS (A, C, and E) and nSL (B, D, and F) under demographic histories Demo 1 (A and B), Demo 3 (C and D), and Demo 4 (E and F) with n = 50 diploid samples. $s$ is the selection coefficient, $f$ is the frequency of the adaptive allele at time of sampling, $g$ is the number of generations at time of sampling since fixation, $e$ is the frequency at which selection began, and $t_d = 2000$ is the time in generations since the two populations diverged.

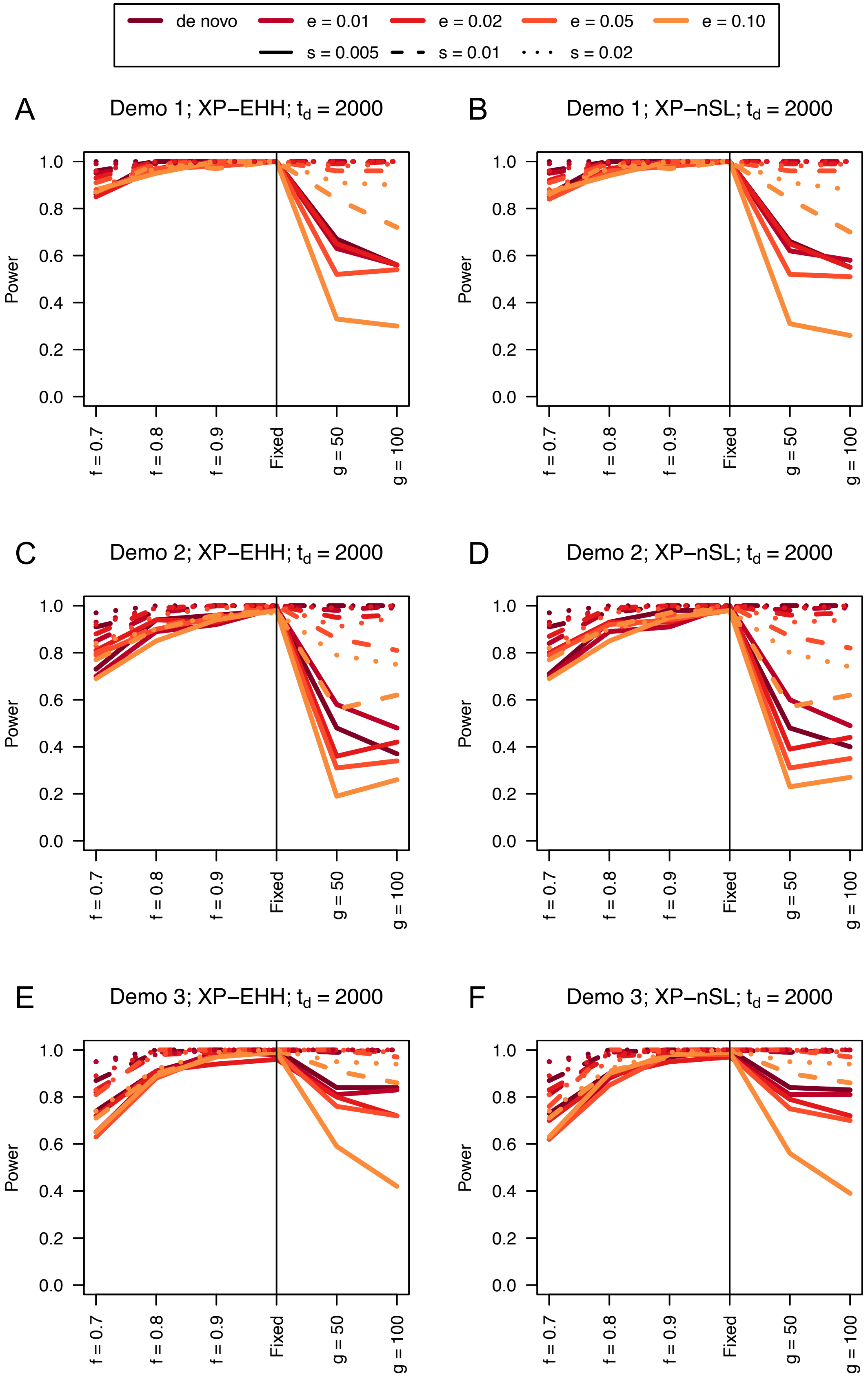

de novo
e = 0.01
e = 0.02
e = 0.05
e = 0.10
s = 0.005
s = 0.01
s = 0.02
A
Demo 1; XP−EHH; t_d = 2000
B
Demo 1; XP−nSL; t_d = 2000
C
Demo 2; XP−EHH; t_d = 2000
D
Demo 2; XP−nSL; t_d = 2000
E
Demo 3; XP−EHH; t_d = 2000
F
Demo 3; XP−nSL; t_d = 2000
Power
f = 0.7
f = 0.8
f = 0.9
Fixed
g = 50
g = 100

**Figure S20**. Power curves for unphased implementations of XP-EHH (A, C, and E) and XP-nSL (B, D, and F) under demographic histories Demo 1 (A and B), Demo 2 (C and D), and Demo 3 (E and F) with n = 50 diploid samples from each population. $s$ is the selection coefficient, $f$ is the frequency of the adaptive allele at time of sampling, $g$ is the number of generations at time of sampling since fixation, $e$ is the frequency at which selection began, and $t_d = 2000$ is the time in generations since the two populations diverged.

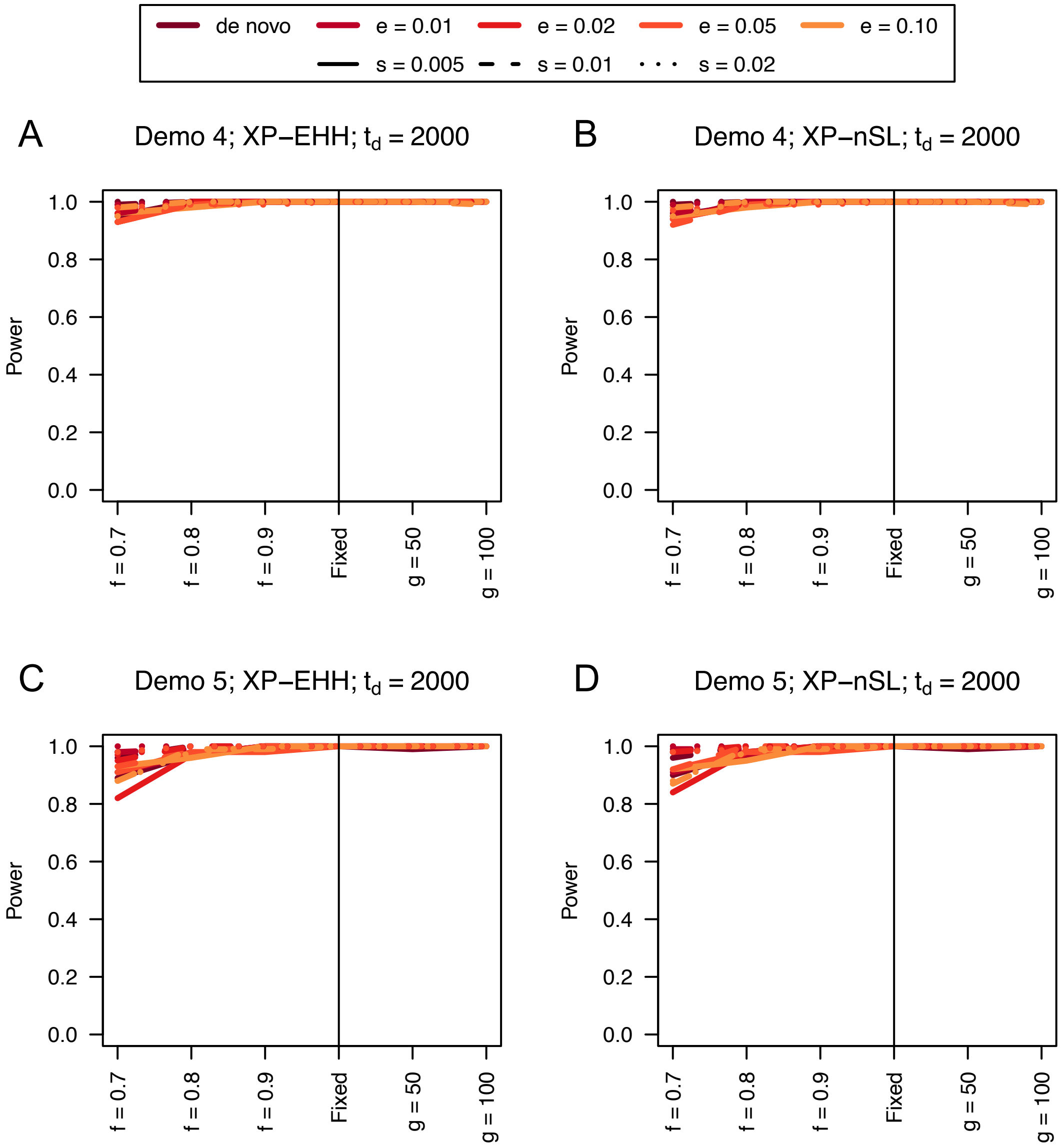


**Figure S21**. Power curves for unphased implementations of XP-EHH (A and C) and XP-nSL (B and D) under demographic histories Demo 4 (A and B), and Demo 5 (C and D) with n = 50 diploid samples from each population. $s$ is the selection coefficient, $f$ is the frequency of the adaptive allele at time of sampling, $g$ is the number of generations at time of sampling since fixation, $e$ is the frequency at which selection began, and $t_d = 2000$ is the time in generations since the two populations diverged.

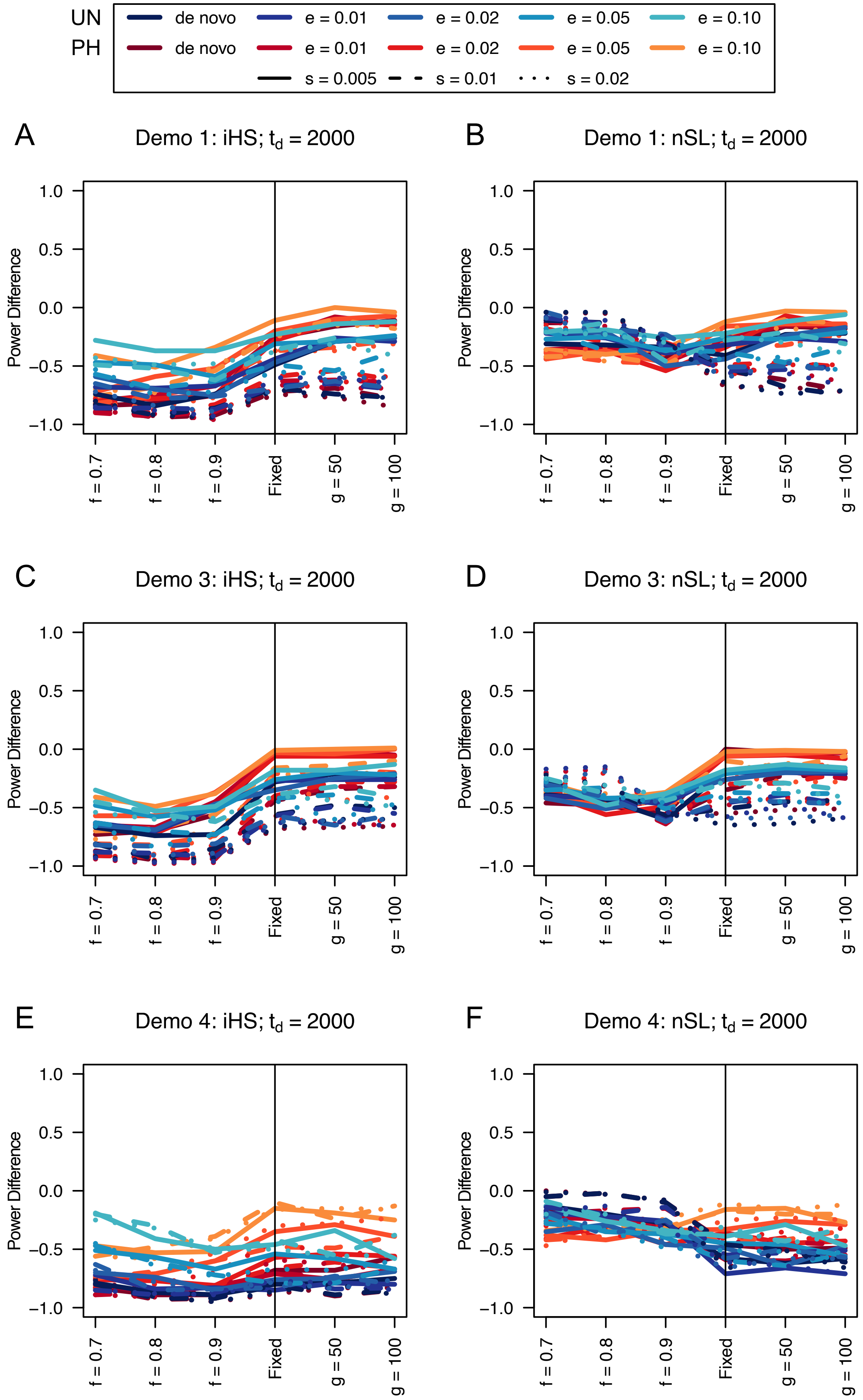
UN
PH
de novo
e = 0.01
e = 0.02
e = 0.05
e = 0.10
s = 0.005
s = 0.01
s = 0.02
A
Demo 1: iHS; $t_d$ = 2000
B
Demo 1: nSL; $t_d$ = 2000
C
Demo 3: iHS; $t_d$ = 2000
D
Demo 3: nSL; $t_d$ = 2000
E
Demo 4: iHS; $t_d$ = 2000
F
Demo 4: nSL; $t_d$ = 2000
Power Difference
1.0
0.5
0.0
−0.5
−1.0
f = 0.7
f = 0.8
f = 0.9
Fixed
g = 50
g = 100

**Figure S22**. Power difference between unphased implementations of iHS (A, C, and E) and nSL (B, D, and F) and phased implementations. Blue curves represent the power difference between the unphased and phased statistics when applied to unphased data (UN). Red curves represent the power difference between the unphased and phased statistics when applied to perfectly phased data (PH). Values greater than 0 indicate the unphased statistic had higher power. Applied to demographic histories Demo 1 (A and B), Demo 3 (C and D), and Demo 4 (E and F) with n = 50 diploid samples. $s$ is the selection coefficient, $f$ is the frequency of the adaptive allele at time of sampling, $g$ is the number of generations at time of sampling since fixation, $e$ is the frequency at which selection began, and $t_d = 2000$ is the time in generations since the two populations diverged.

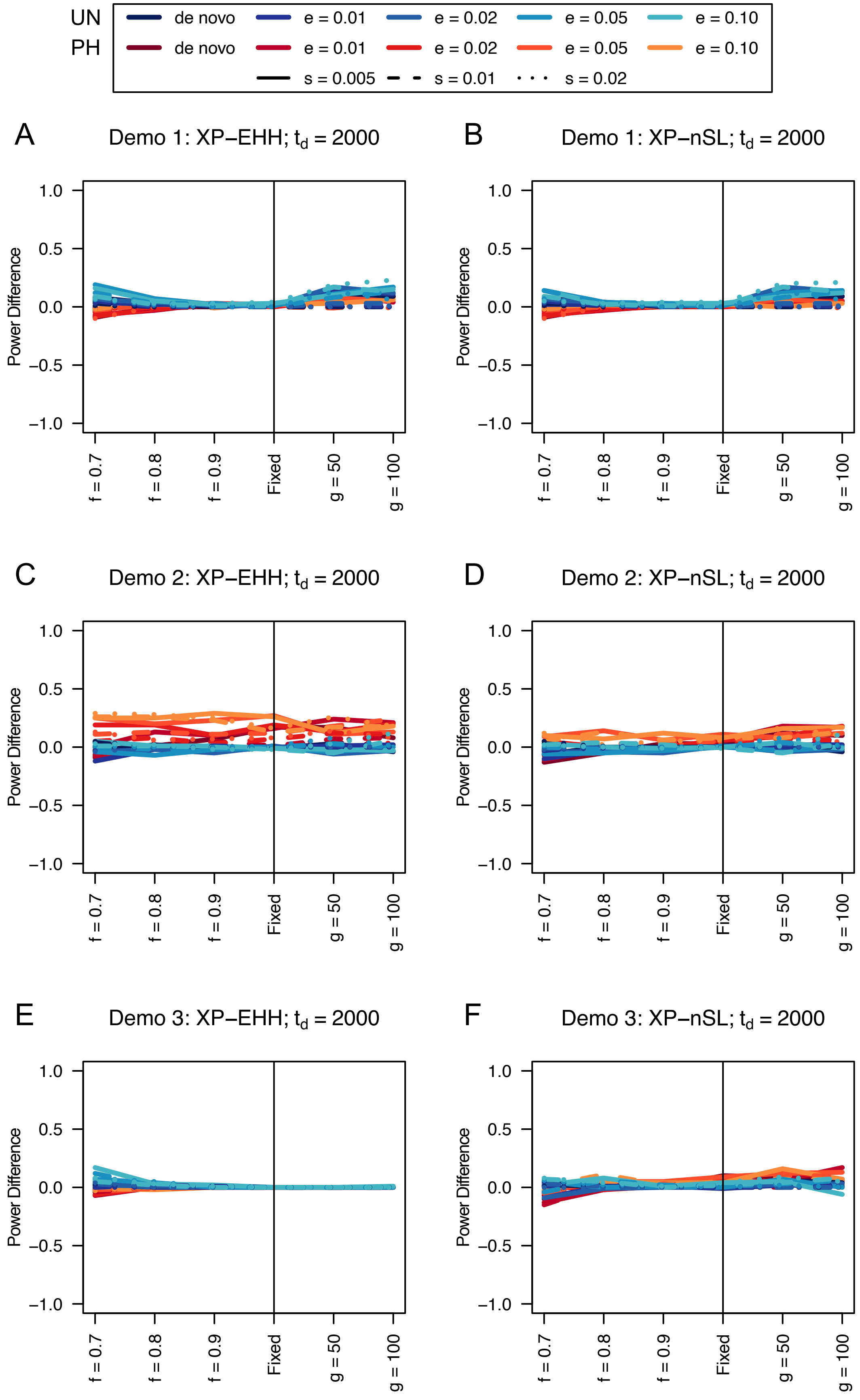
UN de novo e = 0.01 e = 0.02 e = 0.05 e = 0.10
PH de novo e = 0.01 e = 0.02 e = 0.05 e = 0.10
s = 0.005 s = 0.01 s = 0.02
A
Demo 1: XP−EHH; $t_d$ = 2000
B
Demo 1: XP−nSL; $t_d$ = 2000
C
Demo 2: XP−EHH; $t_d$ = 2000
D
Demo 2: XP−nSL; $t_d$ = 2000
E
Demo 3: XP−EHH; $t_d$ = 2000
F
Demo 3: XP−nSL; $t_d$ = 2000
Power Difference
1.0
0.5
0.0
−0.5
−1.0
f = 0.7
f = 0.8
f = 0.9
Fixed
g = 50
g = 100

**Figure S23**. Power difference between unphased implementations of XP-EHH (A, C, and E) and XP-nSL (B, D, and F) and phased implementations. Blue curves represent the power difference between the unphased and phased statistics when applied to unphased data (UN). Red curves represent the power difference between the unphased and phased statistics when applied to perfectly phased data (PH). Values greater than 0 indicate the unphased statistic had higher power. Applied to demographic histories Demo 1 (A and B), Demo 2 (C and D), and Demo 3 (E and F) with n = 50 diploid samples from each population. $s$ is the selection coefficient, $f$ is the frequency of the adaptive allele at time of sampling, $g$ is the number of generations at time of sampling since fixation, $e$ is the frequency at which selection began, and $t_d = 2000$ is the time in generations since the two populations diverged.

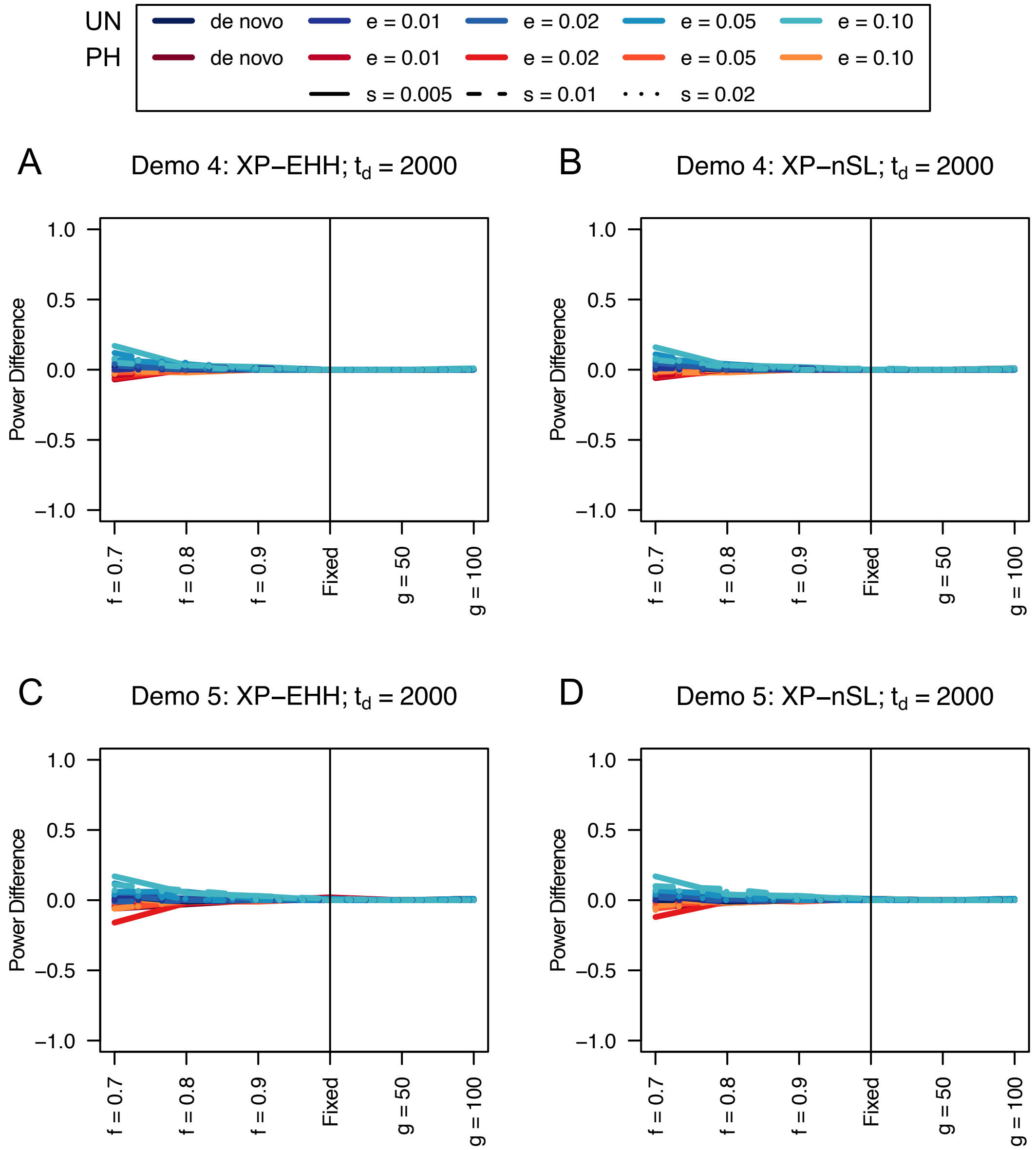


**Figure S24**. Power difference between unphased implementations of XP-EHH (A and C) and XP-nSL (B and D) and phased implementations. Blue curves represent the power difference between the unphased and phased statistics when applied to unphased data (UN). Red curves represent the power difference between the unphased and phased statistics when applied to perfectly phased data (PH). Values greater than 0 indicate the unphased statistic had higher power. Applied to demographic histories Demo 4 (A and B), and Demo 5 (C and D) with n = 50 diploid samples from each population. $s$ is the selection coefficient, $f$ is the frequency of the adaptive allele at time of sampling, $g$ is the number of generations at time of sampling since fixation, $e$ is the frequency at which selection began, and $t_d = 2000$ is the time in generations since the two populations diverged.

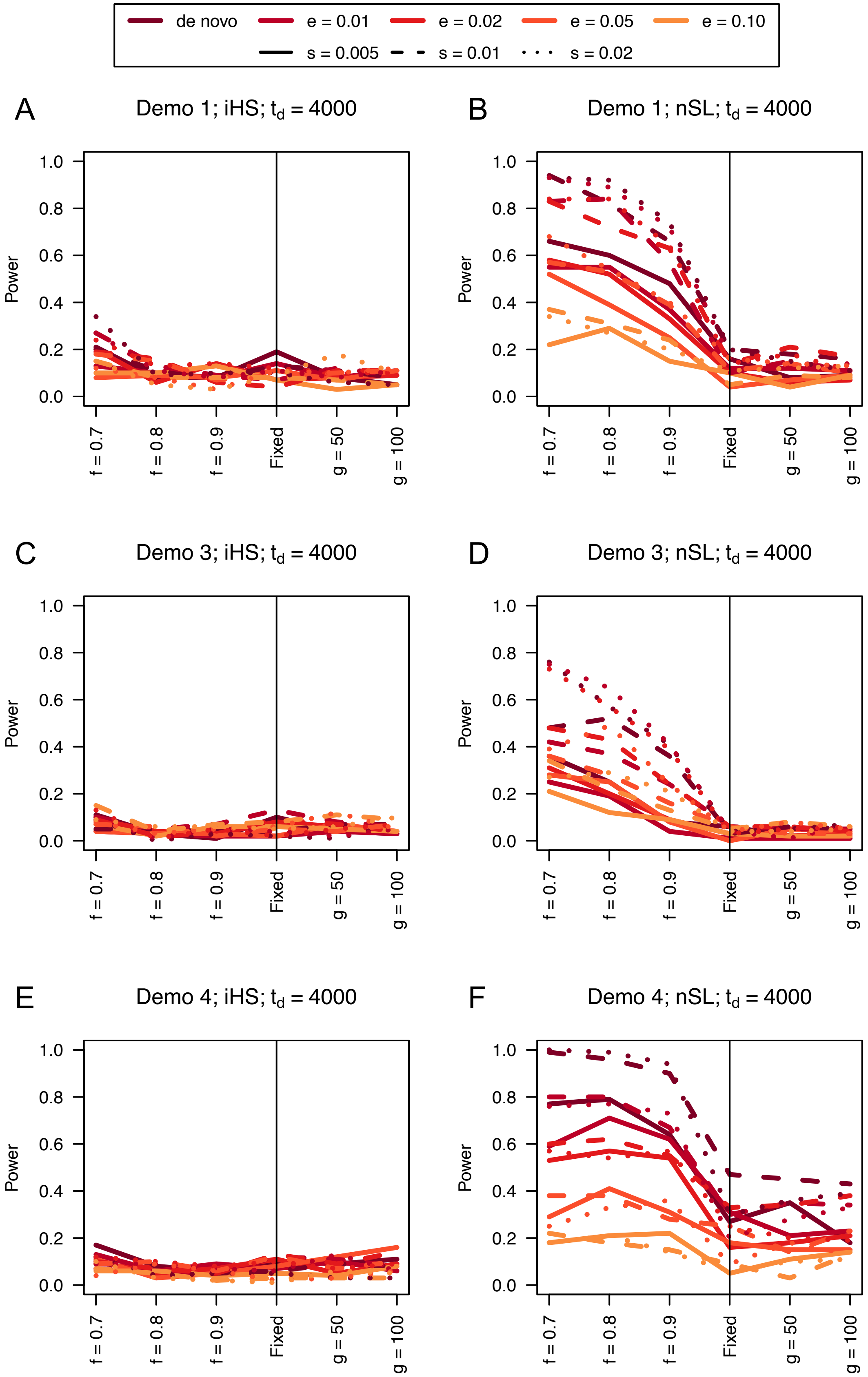
de novo
e = 0.01
e = 0.02
e = 0.05
e = 0.10
s = 0.005
s = 0.01
s = 0.02
A
Demo 1; iHS; $t_d$ = 4000
B
Demo 1; nSL; $t_d$ = 4000
C
Demo 3; iHS; $t_d$ = 4000
D
Demo 3; nSL; $t_d$ = 4000
E
Demo 4; iHS; $t_d$ = 4000
F
Demo 4; nSL; $t_d$ = 4000
Power
1.0
0.8
0.6
0.4
0.2
0.0
f = 0.7
f = 0.8
f = 0.9
Fixed
g = 50
g = 100

**Figure S25**. Power curves for unphased implementations of iHS (A, C, and E) and nSL (B, D, and F) under demographic histories Demo 1 (A and B), Demo 3 (C and D), and Demo 4 (E and F) with n = 50 diploid samples. $s$ is the selection coefficient, $f$ is the frequency of the adaptive allele at time of sampling, $g$ is the number of generations at time of sampling since fixation, $e$ is the frequency at which selection began, and $t_d = 4000$ is the time in generations since the two populations diverged.

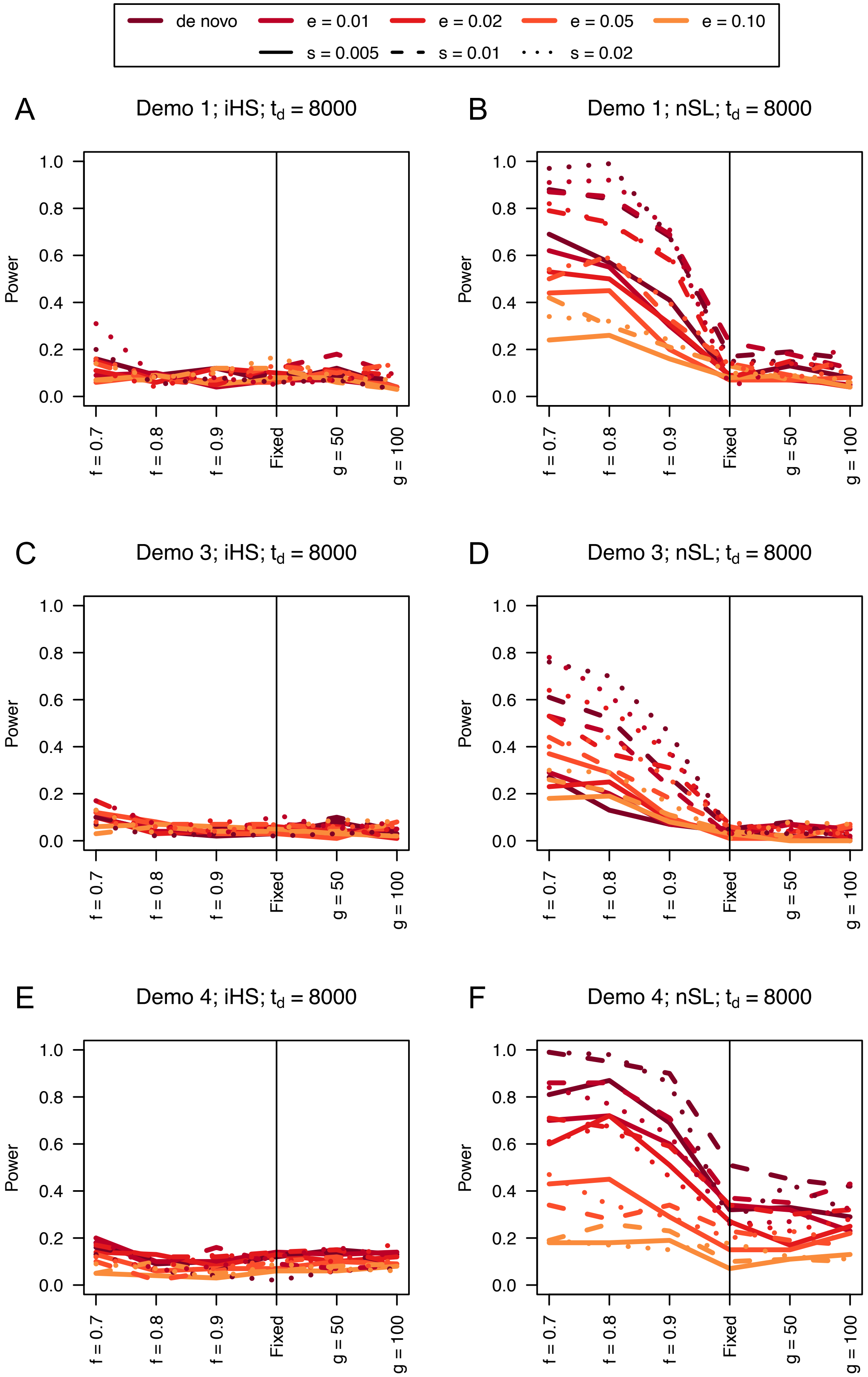
de novo
e = 0.01
e = 0.02
e = 0.05
e = 0.10
s = 0.005
s = 0.01
s = 0.02
A
Demo 1; iHS; $t_d = 8000$
B
Demo 1; nSL; $t_d = 8000$
C
Demo 3; iHS; $t_d = 8000$
D
Demo 3; nSL; $t_d = 8000$
E
Demo 4; iHS; $t_d = 8000$
F
Demo 4; nSL; $t_d = 8000$
Power
1.0
0.8
0.6
0.4
0.2
0.0
f = 0.7
f = 0.8
f = 0.9
Fixed
g = 50
g = 100

**Figure S26**. Power curves for unphased implementations of iHS (A, C, and E) and nSL (B, D, and F) under demographic histories Demo 1 (A and B), Demo 3 (C and D), and Demo 4 (E and F) with n = 50 diploid samples. $s$ is the selection coefficient, $f$ is the frequency of the adaptive allele at time of sampling, $g$ is the number of generations at time of sampling since fixation, $e$ is the frequency at which selection began, and $t_d = 8000$ is the time in generations since the two populations diverged.

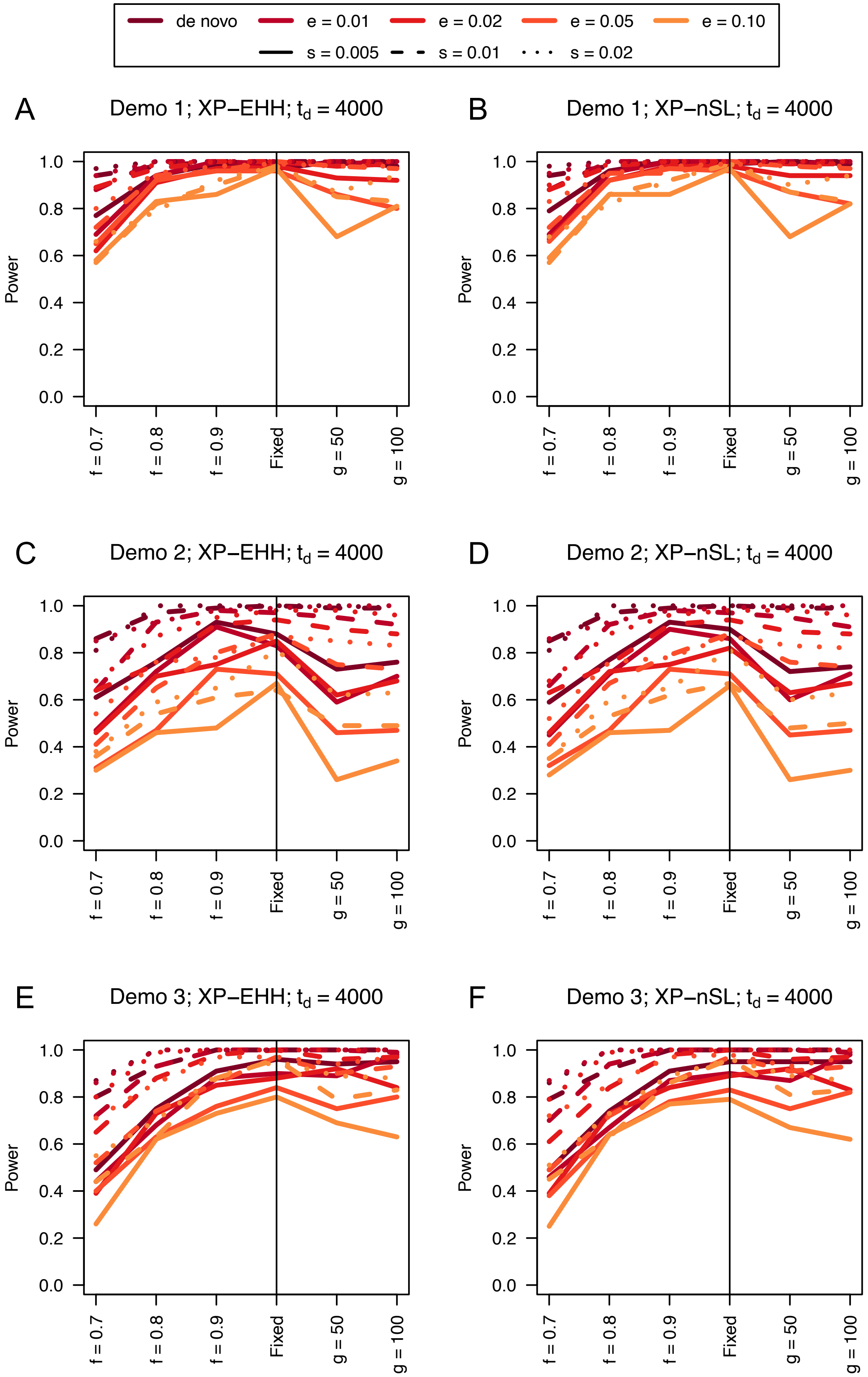

de novo
e = 0.01
e = 0.02
e = 0.05
e = 0.10
s = 0.005
s = 0.01
s = 0.02
A
Demo 1; XP−EHH; $t_d$ = 4000
B
Demo 1; XP−nSL; $t_d$ = 4000
C
Demo 2; XP−EHH; $t_d$ = 4000
D
Demo 2; XP−nSL; $t_d$ = 4000
E
Demo 3; XP−EHH; $t_d$ = 4000
F
Demo 3; XP−nSL; $t_d$ = 4000
Power
0.0
0.2
0.4
0.6
0.8
1.0
f = 0.7
f = 0.8
f = 0.9
Fixed
g = 50
g = 100

**Figure S27**. Power difference between unphased implementations of XP-EHH (A, C, and E) and XP-nSL (B, D, and F) and phased implementations. Blue curves represent the power difference between the unphased and phased statistics when applied to unphased data (UN). Red curves represent the power difference between the unphased and phased statistics when applied to perfectly phased data (PH). Values greater than 0 indicate the unphased statistic had higher power. Applied to demographic histories Demo 1 (A and B), Demo 2 (C and D), and Demo 3 (E and F) with n = 50 diploid samples from each population. $s$ is the selection coefficient, $f$ is the frequency of the adaptive allele at time of sampling, $g$ is the number of generations at time of sampling since fixation, $e$ is the frequency at which selection began, and $t_d = 4000$ is the time in generations since the two populations diverged.

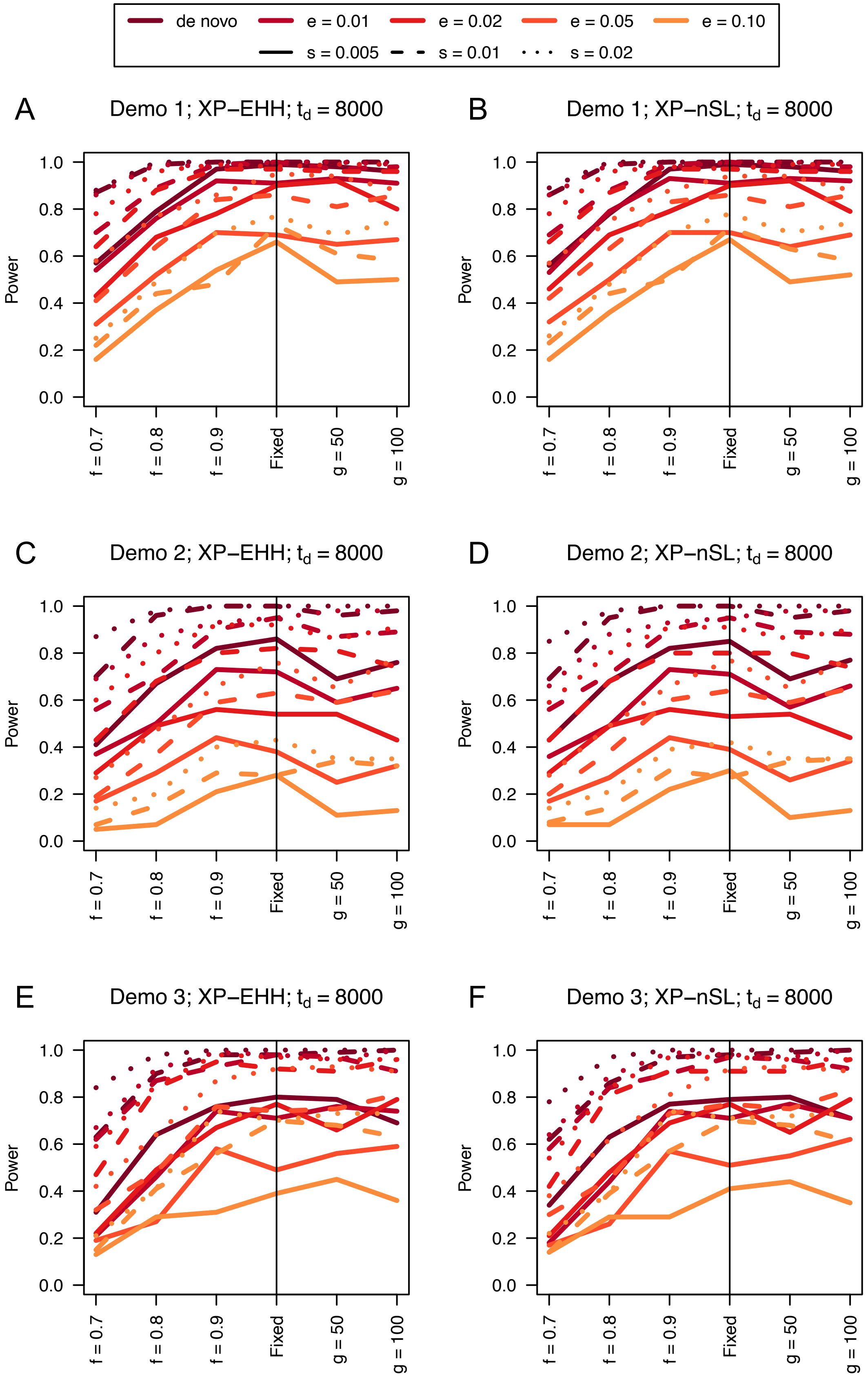
de novo
e = 0.01
e = 0.02
e = 0.05
e = 0.10
s = 0.005
s = 0.01
s = 0.02
A
Demo 1; XP−EHH; $t_d$ = 8000
B
Demo 1; XP−nSL; $t_d$ = 8000
C
Demo 2; XP−EHH; $t_d$ = 8000
D
Demo 2; XP−nSL; $t_d$ = 8000
E
Demo 3; XP−EHH; $t_d$ = 8000
F
Demo 3; XP−nSL; $t_d$ = 8000
Power
0.0
0.2
0.4
0.6
0.8
1.0
f = 0.7
f = 0.8
f = 0.9
Fixed
g = 50
g = 100

**Figure S28**. Power difference between unphased implementations of XP-EHH (A, C, and E) and XP-nSL (B, D, and F) and phased implementations. Blue curves represent the power difference between the unphased and phased statistics when applied to unphased data (UN). Red curves represent the power difference between the unphased and phased statistics when applied to perfectly phased data (PH). Values greater than 0 indicate the unphased statistic had higher power. Applied to demographic histories Demo 1 (A and B), Demo 2 (C and D), and Demo 3 (E and F) with n = 50 diploid samples from each population. $s$ is the selection coefficient, $f$ is the frequency of the adaptive allele at time of sampling, $g$ is the number of generations at time of sampling since fixation, $e$ is the frequency at which selection began, and $t_d = 8000$ is the time in generations since the two populations diverged.

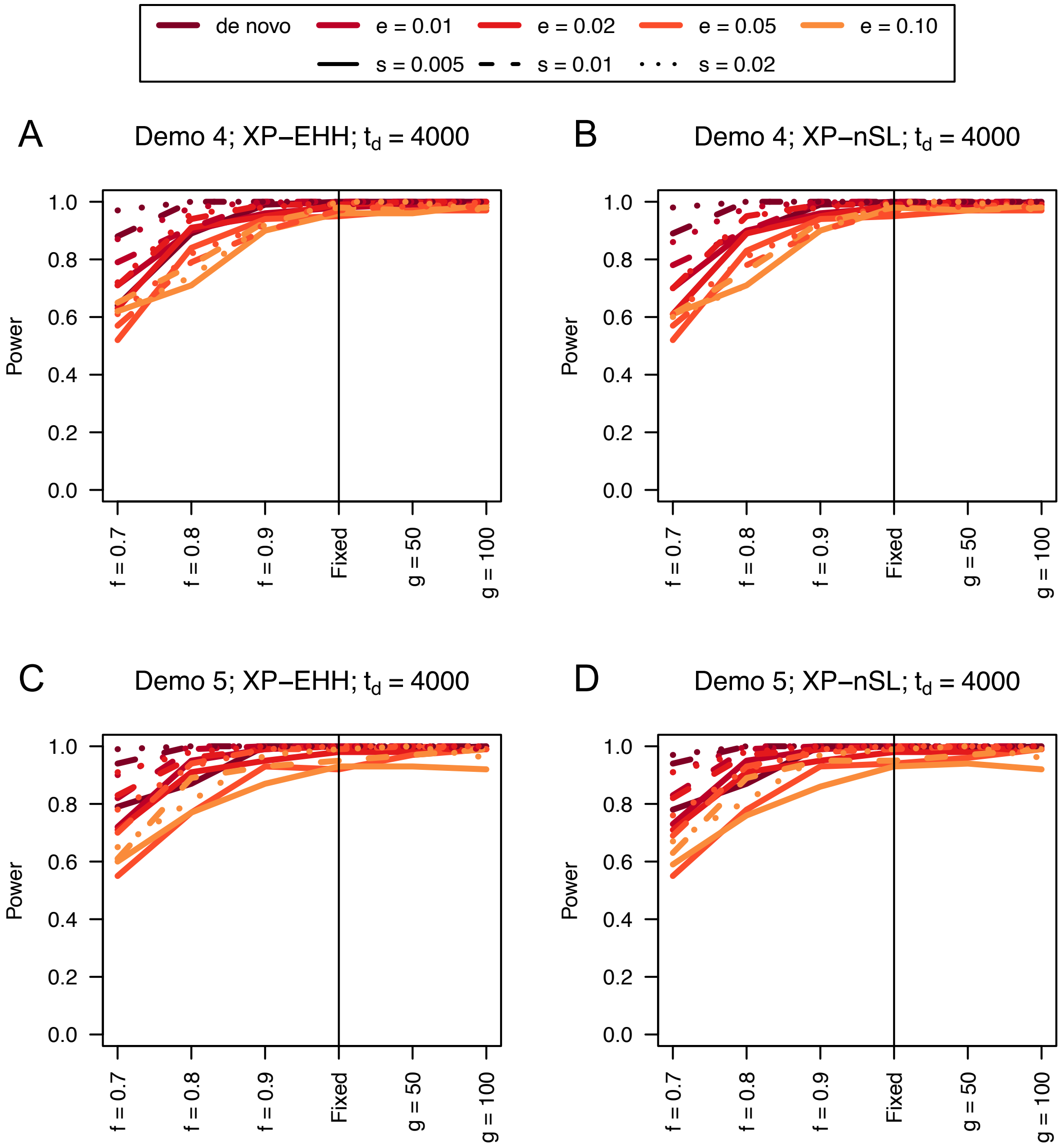


**Figure S29**. Power curves for unphased implementations of XP-EHH (A and C) and XP-nSL (B and D) under demographic histories Demo 4 (A and B), and Demo 5 (C and D) with n = 50 diploid samples from each population. $s$ is the selection coefficient, $f$ is the frequency of the adaptive allele at time of sampling, $g$ is the number of generations at time of sampling since fixation, $e$ is the frequency at which selection began, and $t_d = 4000$ is the time in generations since the two populations diverged.

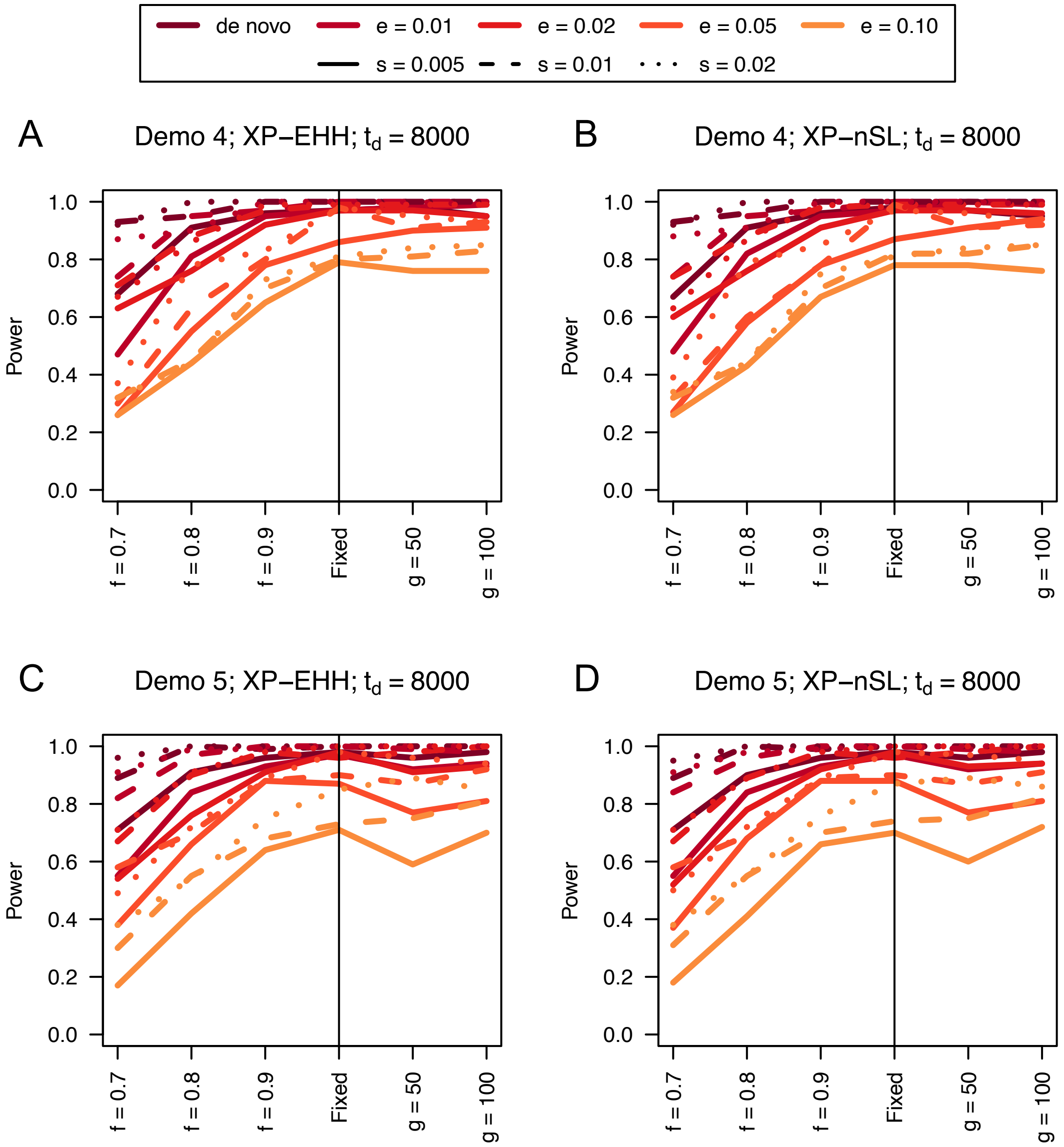


**Figure S30**. Power curves for unphased implementations of XP-EHH (A and C) and XP-nSL (B and D) under demographic histories Demo 4 (A and B), and Demo 5 (C and D) with n = 50 diploid samples from each population. $s$ is the selection coefficient, $f$ is the frequency of the adaptive allele at time of sampling, $g$ is the number of generations at time of sampling since fixation, $e$ is the frequency at which selection began, and $t_d = 8000$ is the time in generations since the two populations diverged.

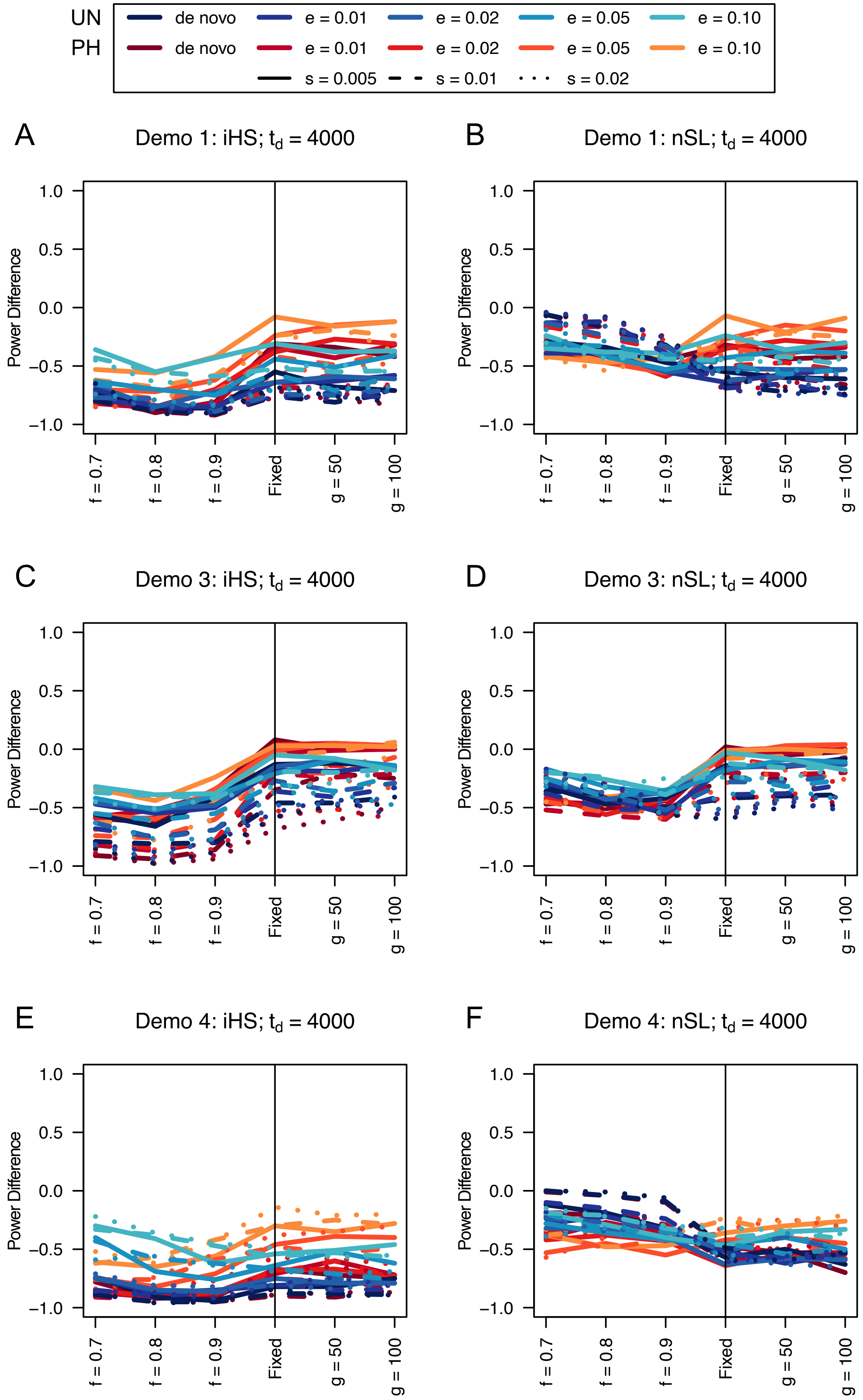

UN de novo e = 0.01 e = 0.02 e = 0.05 e = 0.10
PH de novo e = 0.01 e = 0.02 e = 0.05 e = 0.10
s = 0.005 s = 0.01 s = 0.02
A Demo 1: iHS; $t_d$ = 4000
B Demo 1: nSL; $t_d$ = 4000
C Demo 3: iHS; $t_d$ = 4000
D Demo 3: nSL; $t_d$ = 4000
E Demo 4: iHS; $t_d$ = 4000
F Demo 4: nSL; $t_d$ = 4000
Power Difference
1.0
0.5
0.0
−0.5
−1.0
f = 0.7
f = 0.8
f = 0.9
Fixed
g = 50
g = 100


347

**Figure S31**. Power difference between unphased implementations of iHS (A, C, and E) and nSL (B, D, and F) and phased implementations. Blue curves represent the power difference between the unphased and phased statistics when applied to unphased data (UN). Red curves represent the power difference between the unphased and phased statistics when applied to perfectly phased data (PH). Values greater than 0 indicate the unphased statistic had higher power. Applied to demographic histories Demo 1 (A and B), Demo 3 (C and D), and Demo 4 (E and F) with n = 50 diploid samples. $s$ is the selection coefficient, $f$ is the frequency of the adaptive allele at time of sampling, $g$ is the number of generations at time of sampling since fixation, $e$ is the frequency at which selection began, and $t_d = 4000$ is the time in generations since the two populations diverged.

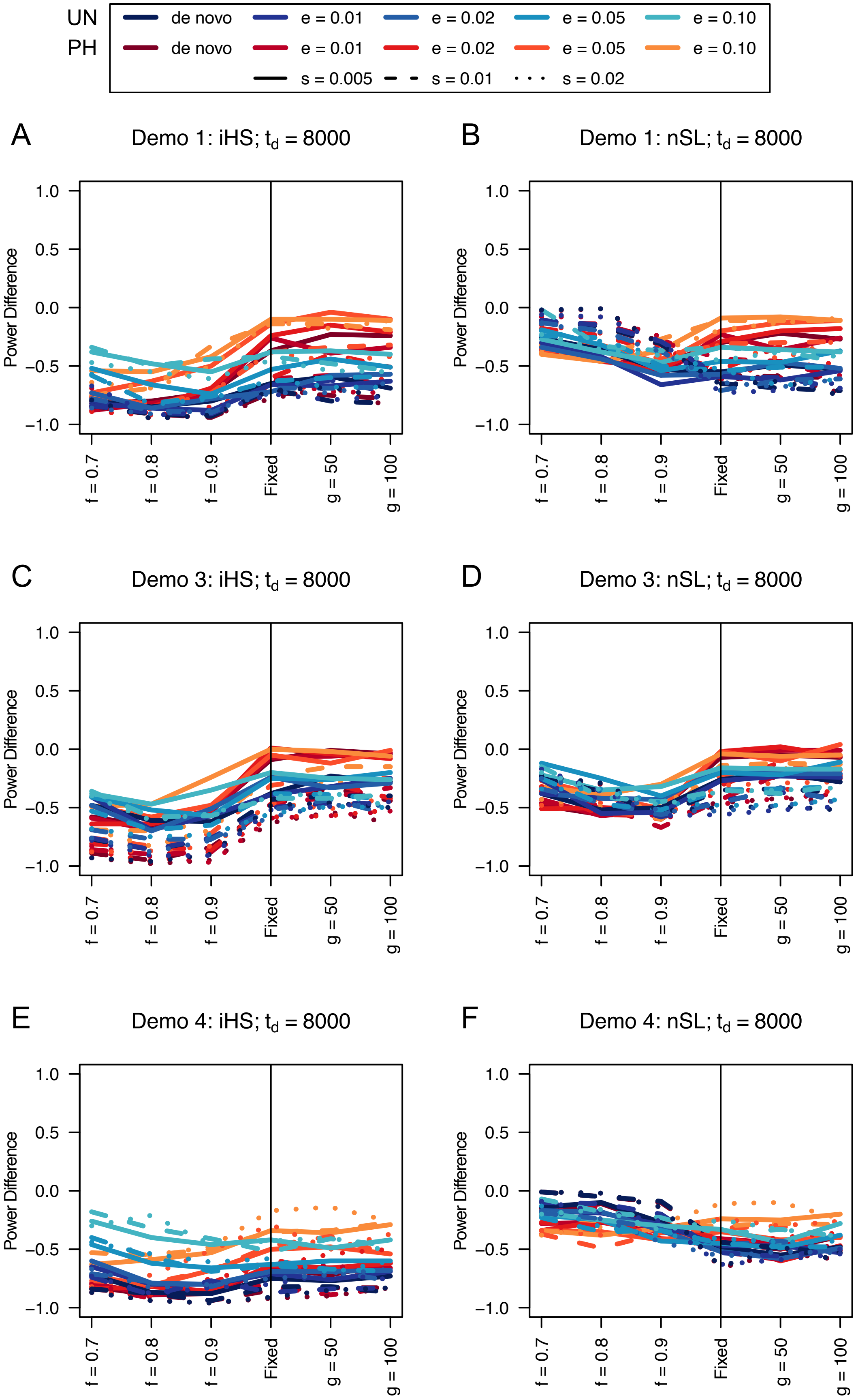
UN
PH
de novo
e = 0.01
e = 0.02
e = 0.05
e = 0.10
s = 0.005
s = 0.01
s = 0.02
A
Demo 1: iHS; $t_d$ = 8000
B
Demo 1: nSL; $t_d$ = 8000
C
Demo 3: iHS; $t_d$ = 8000
D
Demo 3: nSL; $t_d$ = 8000
E
Demo 4: iHS; $t_d$ = 8000
F
Demo 4: nSL; $t_d$ = 8000
Power Difference
1.0
0.5
0.0
−0.5
−1.0
f = 0.7
f = 0.8
f = 0.9
Fixed
g = 50
g = 100

**Figure S32**. Power difference between unphased implementations of iHS (A, C, and E) and nSL (B, D, and F) and phased implementations. Blue curves represent the power difference between the unphased and phased statistics when applied to unphased data (UN). Red curves represent the power difference between the unphased and phased statistics when applied to perfectly phased data (PH). Values greater than 0 indicate the unphased statistic had higher power. Applied to demographic histories Demo 1 (A and B), Demo 3 (C and D), and Demo 4 (E and F) with n = 50 diploid samples. $s$ is the selection coefficient, $f$ is the frequency of the adaptive allele at time of sampling, $g$ is the number of generations at time of sampling since fixation, $e$ is the frequency at which selection began, and $t_d = 8000$ is the time in generations since the two populations diverged.

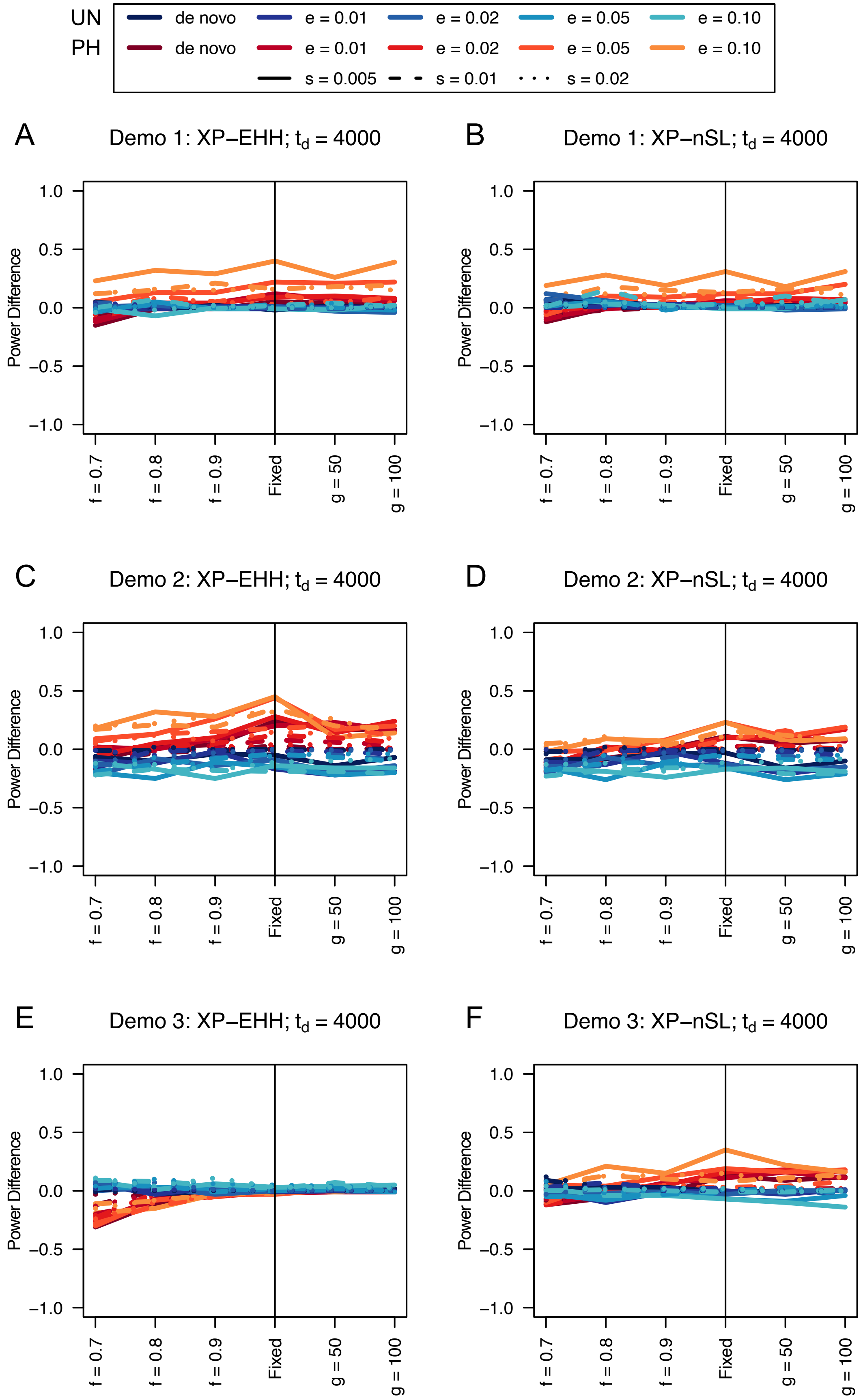
UN
PH
de novo
e = 0.01
e = 0.02
e = 0.05
e = 0.10
s = 0.005
s = 0.01
s = 0.02
A
Demo 1: XP−EHH; $t_d$ = 4000
B
Demo 1: XP−nSL; $t_d$ = 4000
C
Demo 2: XP−EHH; $t_d$ = 4000
D
Demo 2: XP−nSL; $t_d$ = 4000
E
Demo 3: XP−EHH; $t_d$ = 4000
F
Demo 3: XP−nSL; $t_d$ = 4000
Power Difference
1.0
0.5
0.0
−0.5
−1.0
f = 0.7
f = 0.8
f = 0.9
Fixed
g = 50
g = 100

**Figure S33**. Power difference between unphased implementations of XP-EHH (A, C, and E) and XP-nSL (B, D, and F) and phased implementations. Blue curves represent the power difference between the unphased and phased statistics when applied to unphased data (UN). Red curves represent the power difference between the unphased and phased statistics when applied to perfectly phased data (PH). Values greater than 0 indicate the unphased statistic had higher power. Applied to demographic histories Demo 1 (A and B), Demo 2 (C and D), and Demo 3 (E and F) with n = 50 diploid samples from each population. $s$ is the selection coefficient, $f$ is the frequency of the adaptive allele at time of sampling, $g$ is the number of generations at time of sampling since fixation, $e$ is the frequency at which selection began, and $t_d = 4000$ is the time in generations since the two populations diverged.

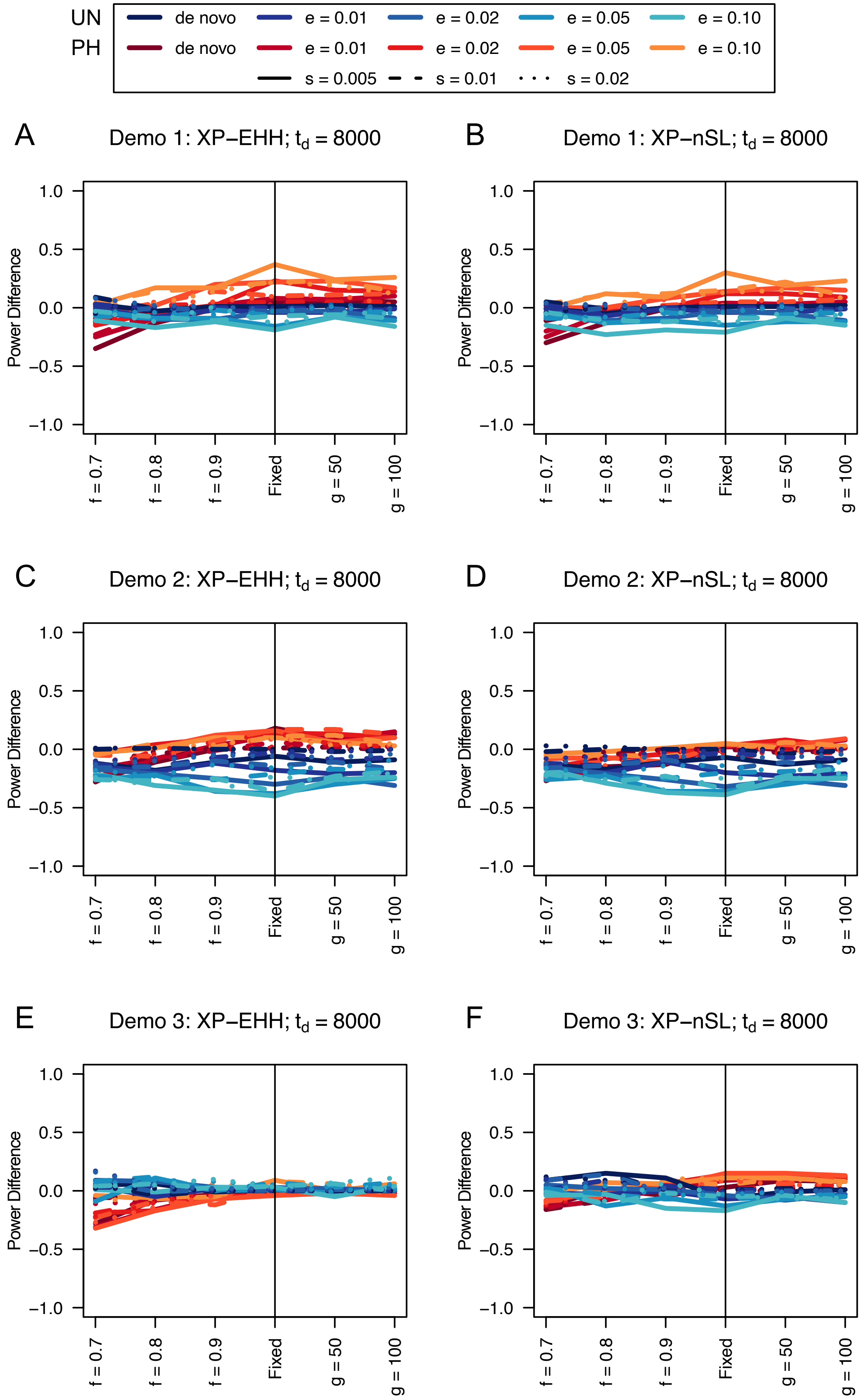

UN
PH
de novo
e = 0.01
e = 0.02
e = 0.05
e = 0.10
s = 0.005
s = 0.01
s = 0.02
A
Demo 1: XP−EHH; $t_d$ = 8000
B
Demo 1: XP−nSL; $t_d$ = 8000
C
Demo 2: XP−EHH; $t_d$ = 8000
D
Demo 2: XP−nSL; $t_d$ = 8000
E
Demo 3: XP−EHH; $t_d$ = 8000
F
Demo 3: XP−nSL; $t_d$ = 8000
Power Difference
1.0
0.5
0.0
−0.5
−1.0
f = 0.7
f = 0.8
f = 0.9
Fixed
g = 50
g = 100

**Figure S34**. Power difference between unphased implementations of XP-EHH (A, C, and E) and XP-nSL (B, D, and F) and phased implementations. Blue curves represent the power difference between the unphased and phased statistics when applied to unphased data (UN). Red curves represent the power difference between the unphased and phased statistics when applied to perfectly phased data (PH). Values greater than 0 indicate the unphased statistic had higher power. Applied to demographic histories Demo 1 (A and B), Demo 2 (C and D), and Demo 3 (E and F) with n = 50 diploid samples from each population. $s$ is the selection coefficient, $f$ is the frequency of the adaptive allele at time of sampling, $g$ is the number of generations at time of sampling since fixation, $e$ is the frequency at which selection began, and $t_d = 8000$ is the time in generations since the two populations diverged.

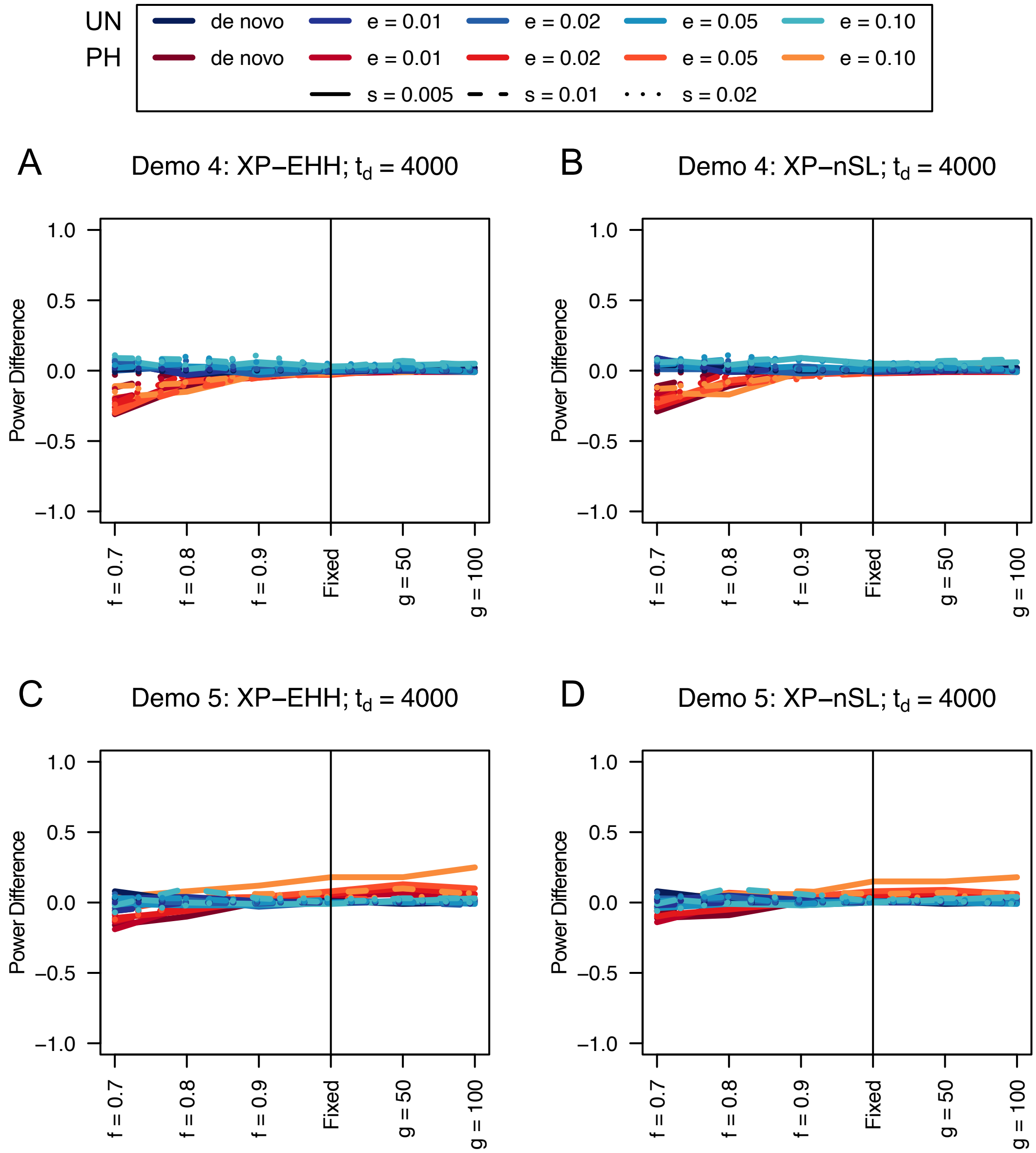


**Figure S35**. Power difference between unphased implementations of XP-EHH (A and C) and XP-nSL (B and D) and phased implementations. Blue curves represent the power difference between the unphased and phased statistics when applied to unphased data (UN). Red curves represent the power difference between the unphased and phased statistics when applied to perfectly phased data (PH). Values greater than 0 indicate the unphased statistic had higher power. Applied to demographic histories Demo 4 (A and B), and Demo 5 (C and D) with n = 50 diploid samples from each population. $s$ is the selection coefficient, $f$ is the frequency of the adaptive allele at time of sampling, $g$ is the number of generations at time of sampling since fixation, $e$ is the frequency at which selection began, and $t_d = 4000$ is the time in generations since the two populations diverged.

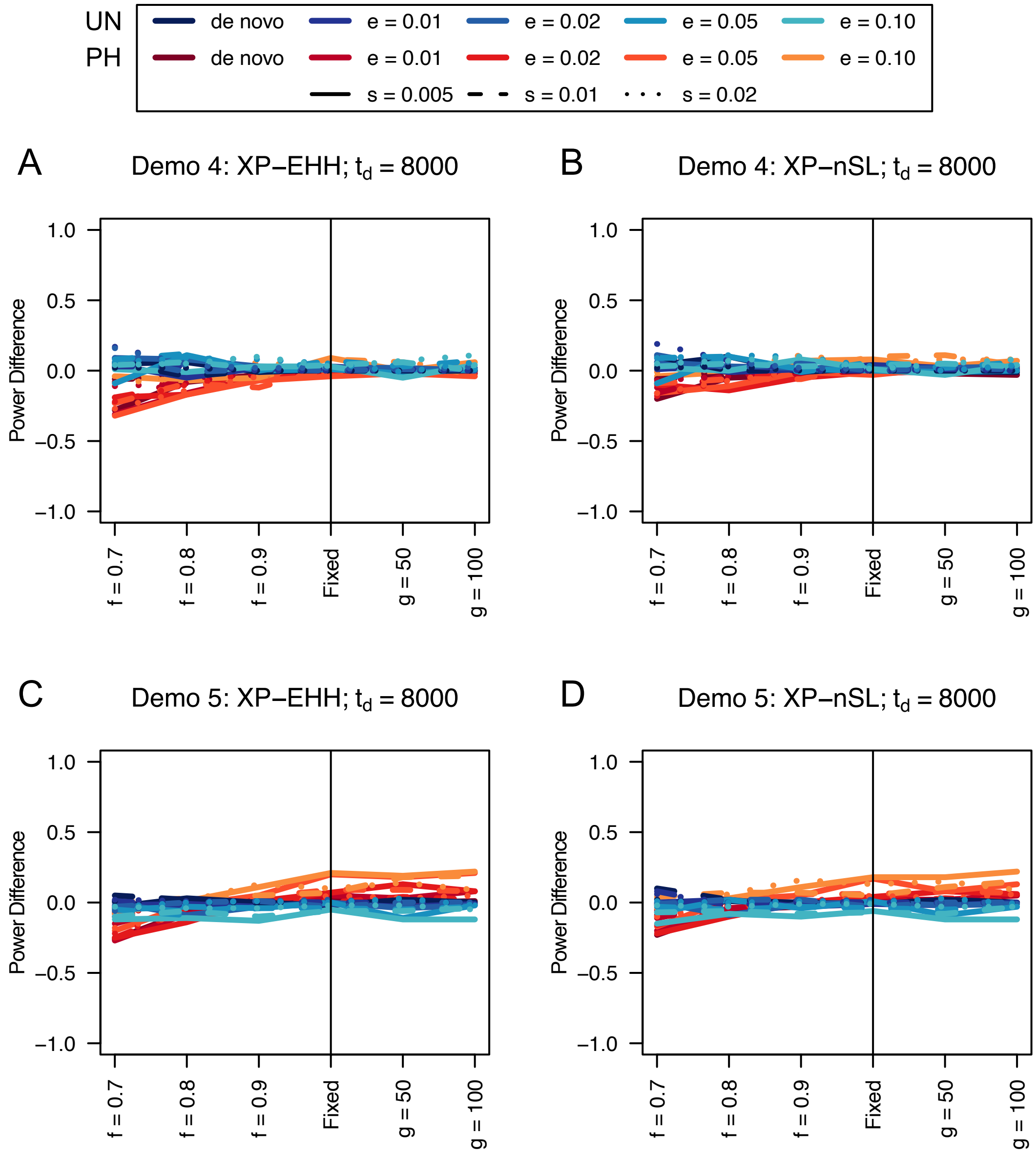


**Table S36**. Power difference between unphased implementations of XP-EHH (A and C) and XP-nSL (B and D) and phased implementations. Blue curves represent the power difference between the unphased and phased statistics when applied to unphased data (UN). Red curves represent the power difference between the unphased and phased statistics when applied to perfectly phased data (PH). Values greater than 0 indicate the unphased statistic had higher power. Applied to demographic histories Demo 4 (A and B), and Demo 5 (C and D) with n = 50 diploid samples from each population. $s$ is the selection coefficient, $f$ is the frequency of the adaptive allele at time of sampling, $g$ is the number of generations at time of sampling since fixation, $e$ is the frequency at which selection began, and $t_d = 8000$ is the time in generations since the two populations diverged.

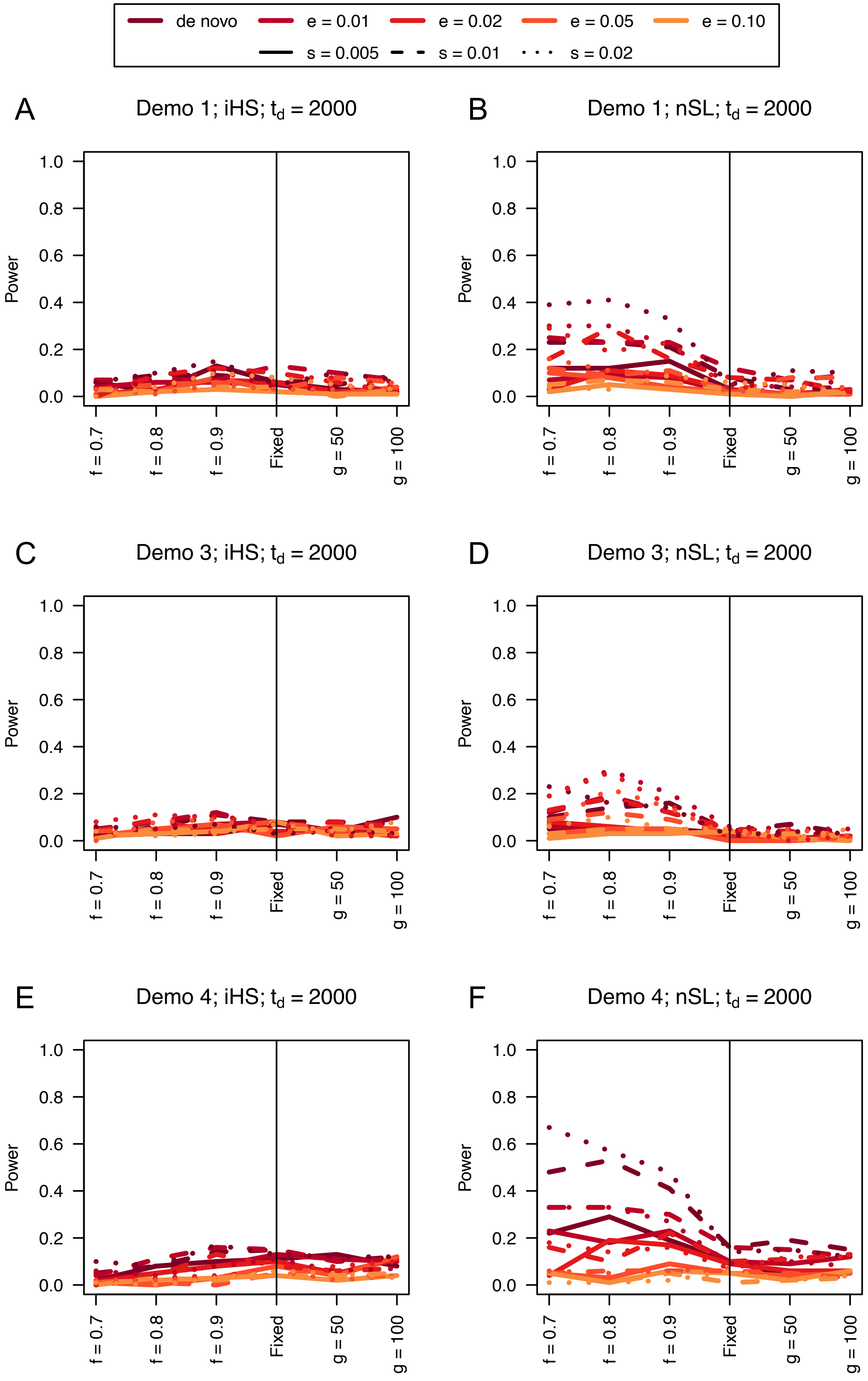

de novo
e = 0.01
e = 0.02
e = 0.05
e = 0.10
s = 0.005
s = 0.01
s = 0.02
A
Demo 1; iHS; $t_d$ = 2000
B
Demo 1; nSL; $t_d$ = 2000
C
Demo 3; iHS; $t_d$ = 2000
D
Demo 3; nSL; $t_d$ = 2000
E
Demo 4; iHS; $t_d$ = 2000
F
Demo 4; nSL; $t_d$ = 2000
Power
0.0
0.2
0.4
0.6
0.8
1.0
f = 0.7
f = 0.8
f = 0.9
Fixed
g = 50
g = 100


415

**Figure S37**. Power curves for unphased implementations of iHS (A, C, and E) and nSL (B, D, and F) under demographic histories Demo 1 (A and B), Demo 3 (C and D), and Demo 4 (E and F) with n = 20 diploid samples. $s$ is the selection coefficient, $f$ is the frequency of the adaptive allele at time of sampling, $g$ is the number of generations at time of sampling since fixation, $e$ is the frequency at which selection began, and $t_d = 2000$ is the time in generations since the two populations diverged.

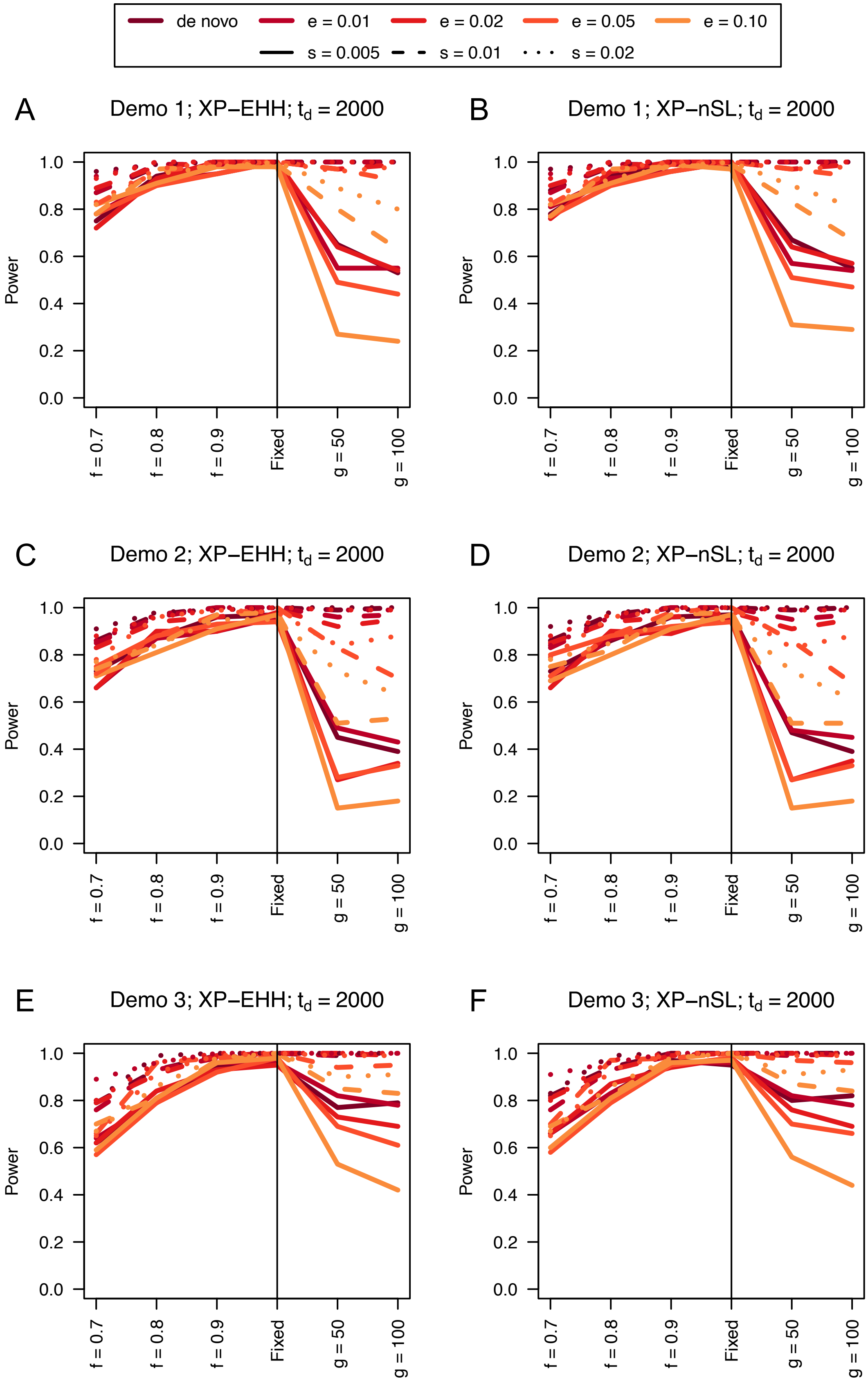

de novo
e = 0.01
e = 0.02
e = 0.05
e = 0.10
s = 0.005
s = 0.01
s = 0.02
A
Demo 1; XP−EHH; $t_d$ = 2000
B
Demo 1; XP−nSL; $t_d$ = 2000
C
Demo 2; XP−EHH; $t_d$ = 2000
D
Demo 2; XP−nSL; $t_d$ = 2000
E
Demo 3; XP−EHH; $t_d$ = 2000
F
Demo 3; XP−nSL; $t_d$ = 2000
Power
0.0
0.2
0.4
0.6
0.8
1.0
f = 0.7
f = 0.8
f = 0.9
Fixed
g = 50
g = 100

**Figure S38**. Power curves for unphased implementations of XP-EHH (A, C, and E) and XP-nSL (B, D, and F) under demographic histories Demo 1 (A and B), Demo 2 (C and D), and Demo 3 (E and F) with n = 20 diploid samples from each population. $s$ is the selection coefficient, $f$ is the frequency of the adaptive allele at time of sampling, $g$ is the number of generations at time of sampling since fixation, $e$ is the frequency at which selection began, and $t_d = 2000$ is the time in generations since the two populations diverged.

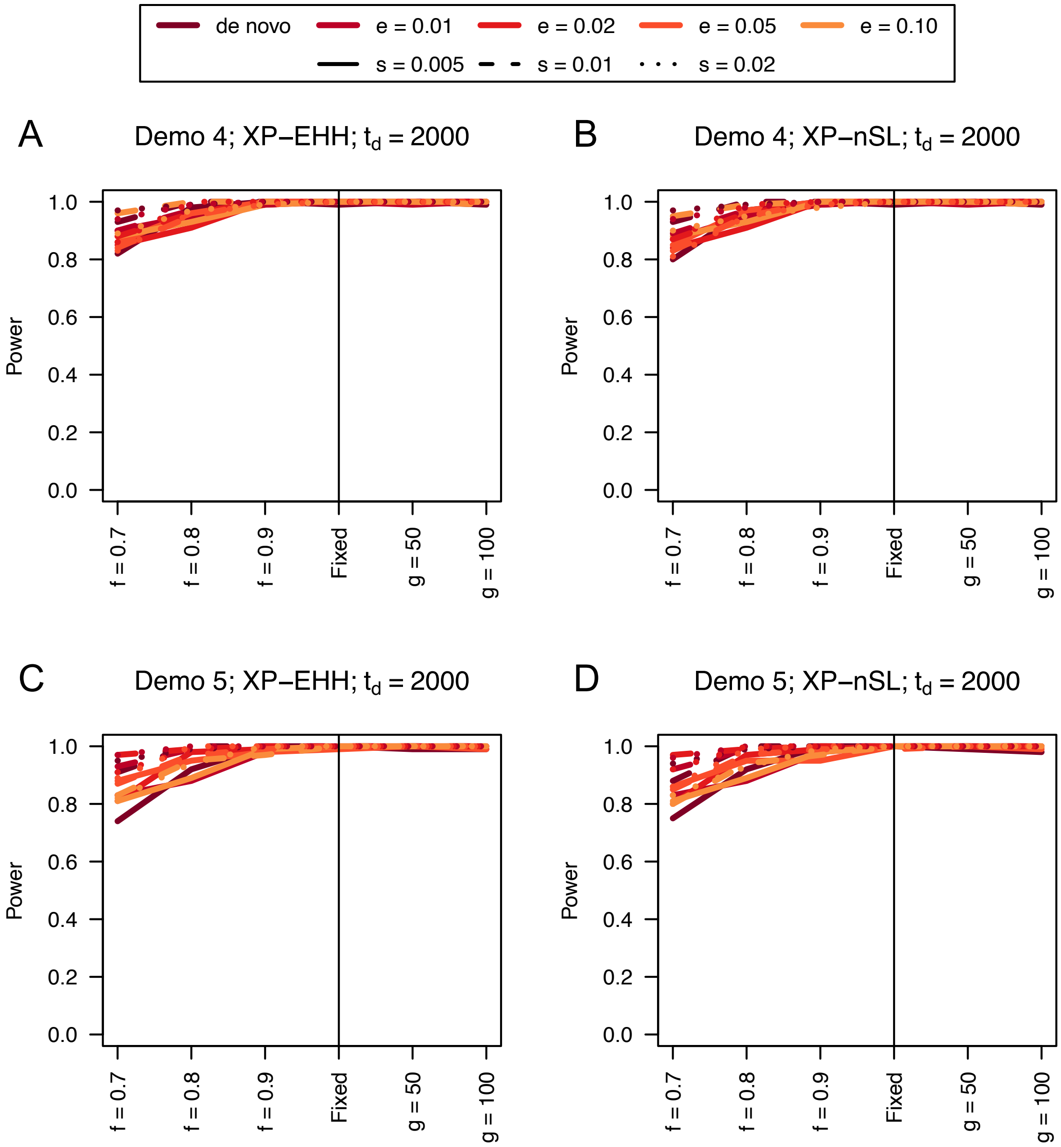


**Figure S39**. Power curves for unphased implementations of XP-EHH (A and C) and XP-nSL (B and D) under demographic histories Demo 4 (A and B), and Demo 5 (C and D) with n = 20 diploid samples from each population. $s$ is the selection coefficient, $f$ is the frequency of the adaptive allele at time of sampling, $g$ is the number of generations at time of sampling since fixation, $e$ is the frequency at which selection began, and $t_d = 2000$ is the time in generations since the two populations diverged.

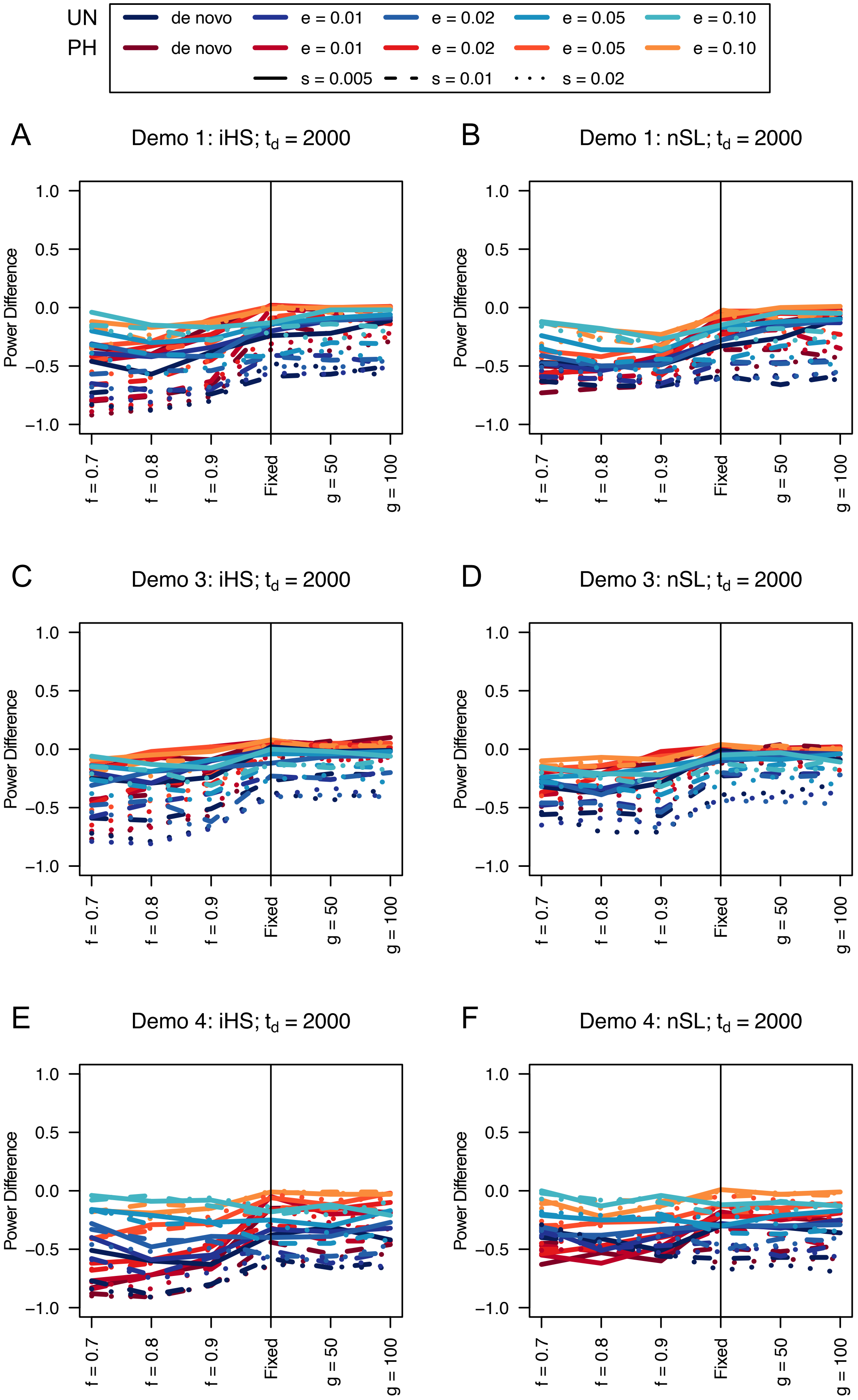

UN
PH
de novo
e = 0.01
e = 0.02
e = 0.05
e = 0.10
s = 0.005
s = 0.01
s = 0.02
A
Demo 1: iHS; $t_d$ = 2000
B
Demo 1: nSL; $t_d$ = 2000
C
Demo 3: iHS; $t_d$ = 2000
D
Demo 3: nSL; $t_d$ = 2000
E
Demo 4: iHS; $t_d$ = 2000
F
Demo 4: nSL; $t_d$ = 2000
Power Difference
1.0
0.5
0.0
−0.5
−1.0
f = 0.7
f = 0.8
f = 0.9
Fixed
g = 50
g = 100

**Figure S40**. Power difference between unphased implementations of iHS (A, C, and E) and nSL (B, D, and F) and phased implementations. Blue curves represent the power difference between the unphased and phased statistics when applied to unphased data (UN). Red curves represent the power difference between the unphased and phased statistics when applied to perfectly phased data (PH). Values greater than 0 indicate the unphased statistic had higher power. Applied to demographic histories Demo 1 (A and B), Demo 3 (C and D), and Demo 4 (E and F) with n = 20 diploid samples. $s$ is the selection coefficient, $f$ is the frequency of the adaptive allele at time of sampling, $g$ is the number of generations at time of sampling since fixation, $e$ is the frequency at which selection began, and $t_d = 2000$ is the time in generations since the two populations diverged.

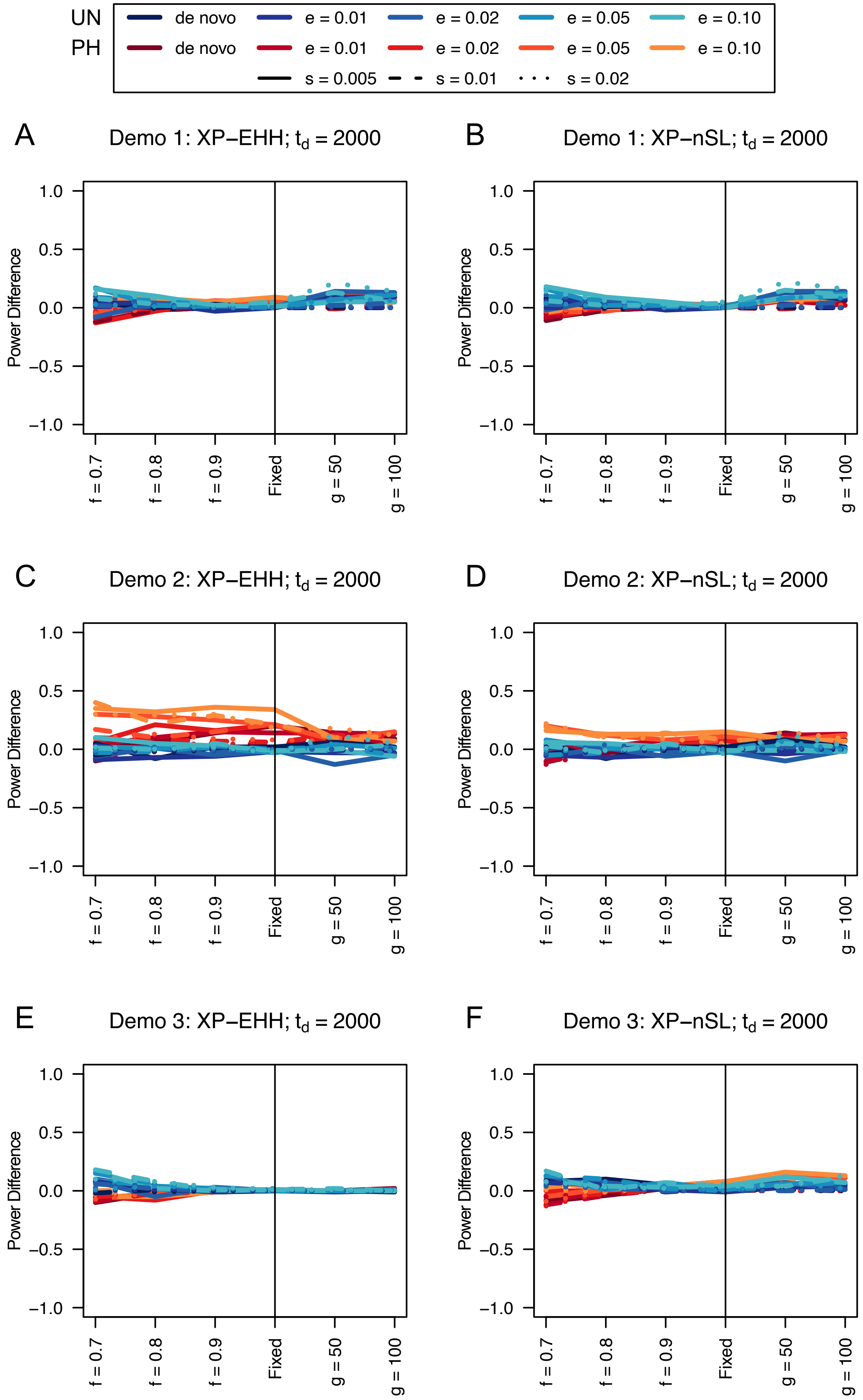
UN
de novo
e = 0.01
e = 0.02
e = 0.05
e = 0.10
PH
de novo
e = 0.01
e = 0.02
e = 0.05
e = 0.10
s = 0.005
s = 0.01
s = 0.02
A
Demo 1: XP−EHH; t_d = 2000
B
Demo 1: XP−nSL; t_d = 2000
C
Demo 2: XP−EHH; t_d = 2000
D
Demo 2: XP−nSL; t_d = 2000
E
Demo 3: XP−EHH; t_d = 2000
F
Demo 3: XP−nSL; t_d = 2000
Power Difference
1.0
0.5
0.0
−0.5
−1.0
f = 0.7
f = 0.8
f = 0.9
Fixed
g = 50
g = 100

**Figure S41**. Power difference between unphased implementations of XP-EHH (A, C, and E) and XP-nSL (B, D, and F) and phased implementations. Blue curves represent the power difference between the unphased and phased statistics when applied to unphased data (UN). Red curves represent the power difference between the unphased and phased statistics when applied to perfectly phased data (PH). Values greater than 0 indicate the unphased statistic had higher power. Applied to demographic histories Demo 1 (A and B), Demo 2 (C and D), and Demo 3 (E and F) with n = 20 diploid samples from each population. $s$ is the selection coefficient, $f$ is the frequency of the adaptive allele at time of sampling, $g$ is the number of generations at time of sampling since fixation, $e$ is the frequency at which selection began, and $t_d = 2000$ is the time in generations since the two populations diverged.

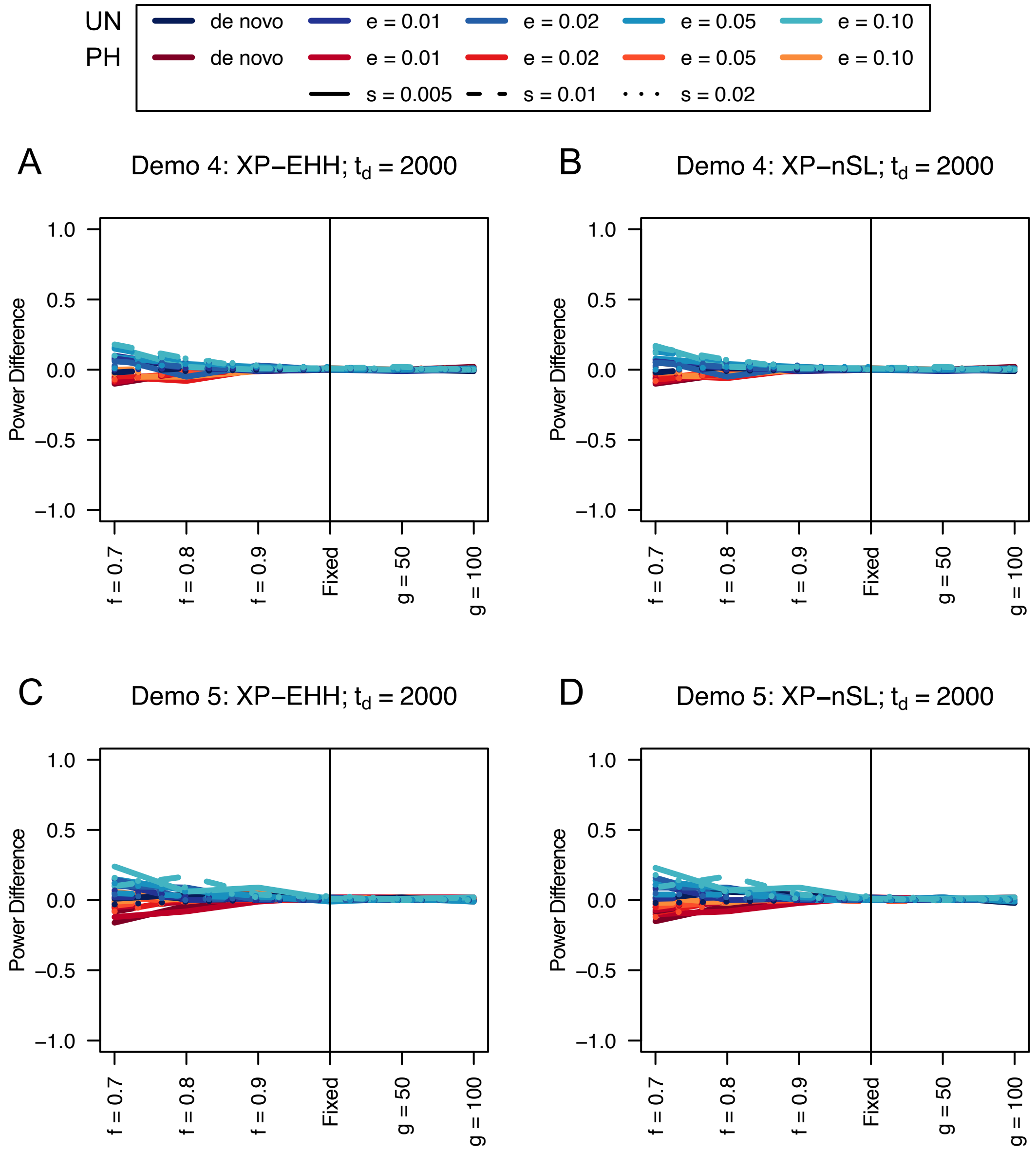


**Figure S42**. Power difference between unphased implementations of XP-EHH (A and C) and XP-nSL (B and D) and phased implementations. Blue curves represent the power difference between the unphased and phased statistics when applied to unphased data (UN). Red curves represent the power difference between the unphased and phased statistics when applied to perfectly phased data (PH). Values greater than 0 indicate the unphased statistic had higher power. Applied to demographic histories Demo 4 (A and B), and Demo 5 (C and D) with n = 20 diploid samples from each population. $s$ is the selection coefficient, $f$ is the frequency of the adaptive allele at time of sampling, $g$ is the number of generations at time of sampling since fixation, $e$ is the frequency at which selection began, and $t_d = 2000$ is the time in generations since the two populations diverged.

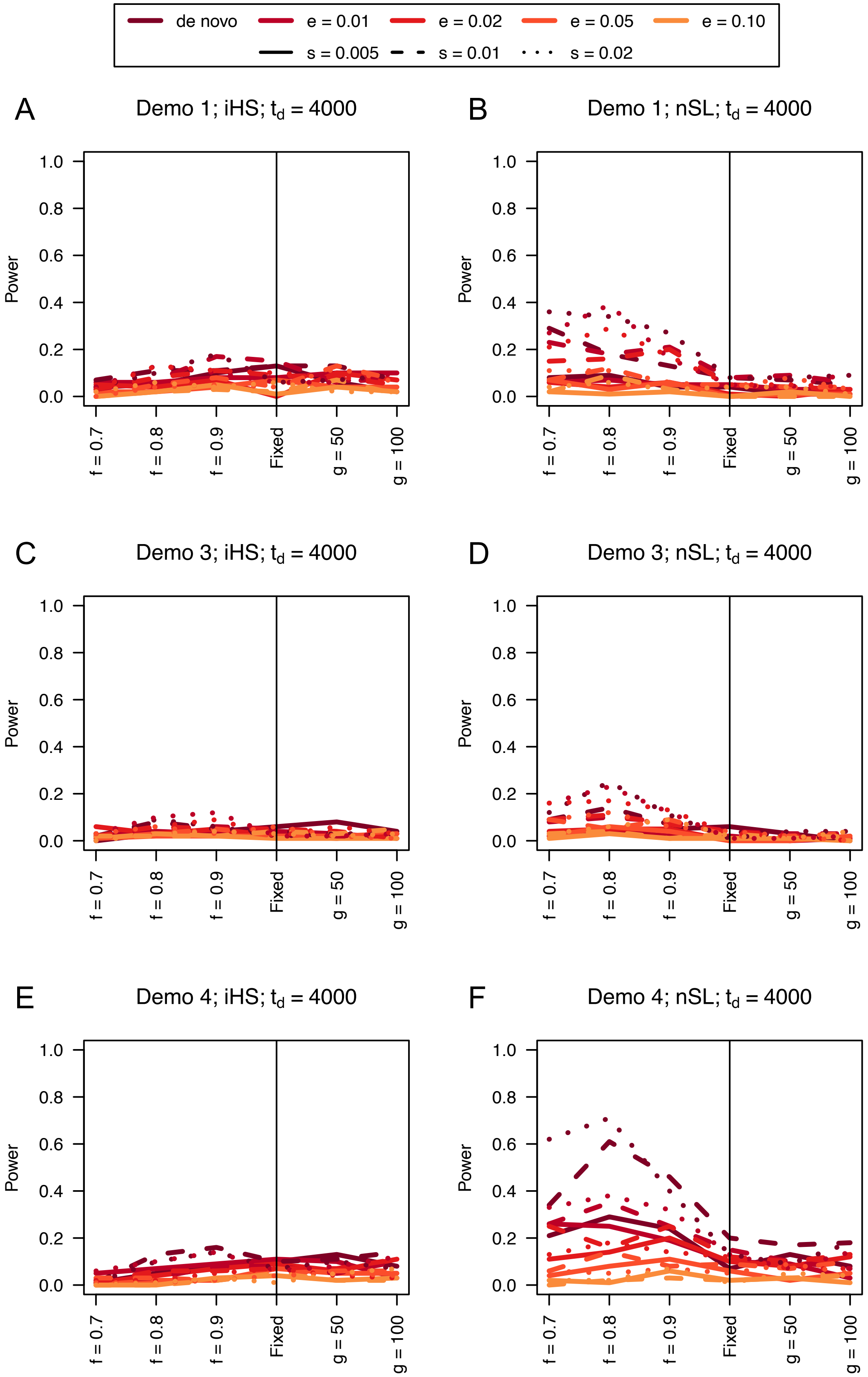

de novo
e = 0.01
e = 0.02
e = 0.05
e = 0.10
s = 0.005
s = 0.01
s = 0.02
A
Demo 1; iHS; $t_d$ = 4000
B
Demo 1; nSL; $t_d$ = 4000
C
Demo 3; iHS; $t_d$ = 4000
D
Demo 3; nSL; $t_d$ = 4000
E
Demo 4; iHS; $t_d$ = 4000
F
Demo 4; nSL; $t_d$ = 4000
Power
1.0
0.8
0.6
0.4
0.2
0.0
f = 0.7
f = 0.8
f = 0.9
Fixed
g = 50
g = 100


484

**Figure S43**. Power curves for unphased implementations of iHS (A, C, and E) and nSL (B, D, and F) under demographic histories Demo 1 (A and B), Demo 3 (C and D), and Demo 4 (E and F) with n = 20 diploid samples. $s$ is the selection coefficient, $f$ is the frequency of the adaptive allele at time of sampling, $g$ is the number of generations at time of sampling since fixation, $e$ is the frequency at which selection began, and $t_d = 4000$ is the time in generations since the two populations diverged.

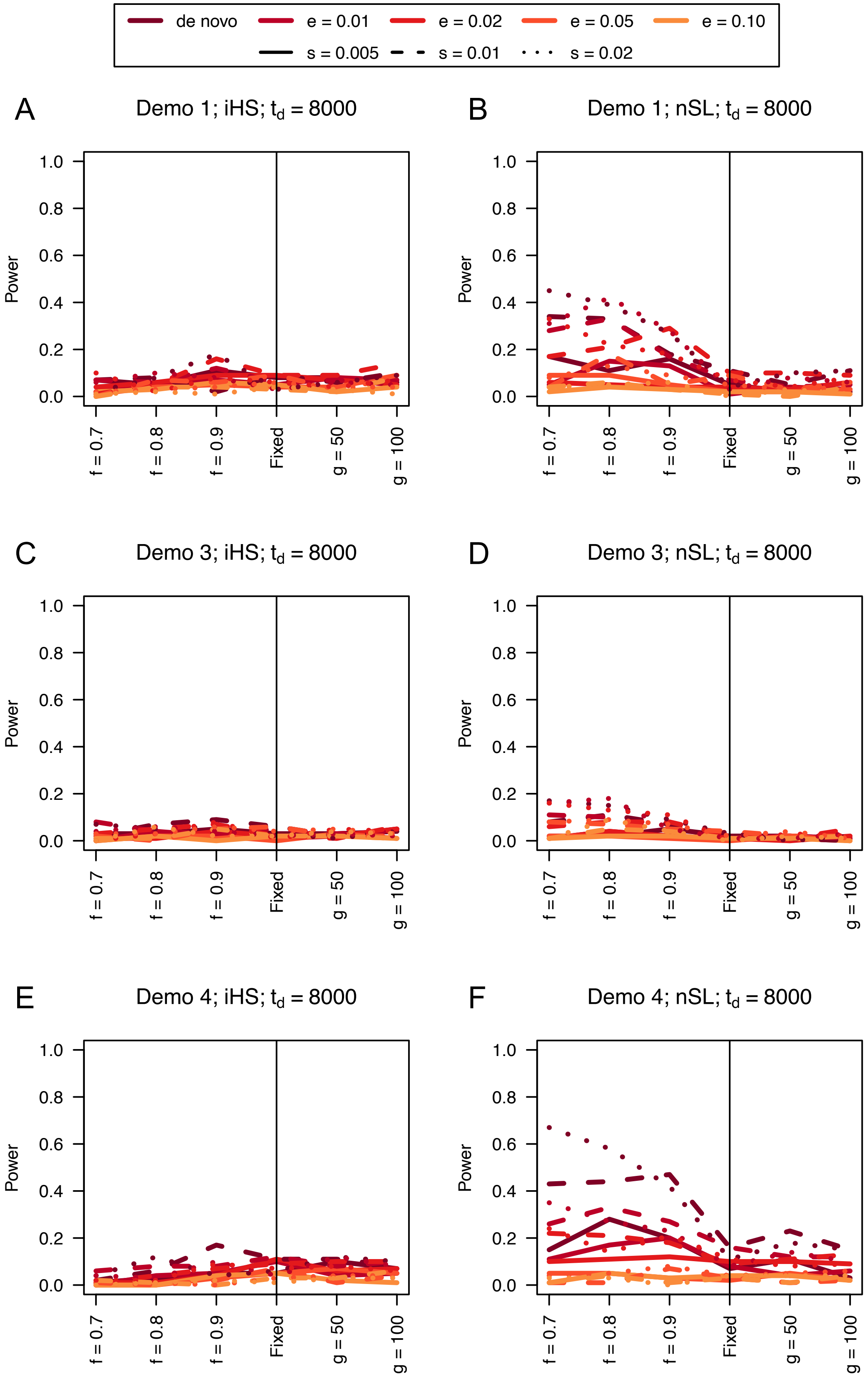

de novo
e = 0.01
e = 0.02
e = 0.05
e = 0.10
s = 0.005
s = 0.01
s = 0.02
A
Demo 1; iHS; $t_d$ = 8000
B
Demo 1; nSL; $t_d$ = 8000
C
Demo 3; iHS; $t_d$ = 8000
D
Demo 3; nSL; $t_d$ = 8000
E
Demo 4; iHS; $t_d$ = 8000
F
Demo 4; nSL; $t_d$ = 8000
Power
1.0
0.8
0.6
0.4
0.2
0.0
f = 0.7
f = 0.8
f = 0.9
Fixed
g = 50
g = 100

**Figure S44**. Power curves for unphased implementations of iHS (A, C, and E) and nSL (B, D, and F) under demographic histories Demo 1 (A and B), Demo 3 (C and D), and Demo 4 (E and F) with n = 20 diploid samples. $s$ is the selection coefficient, $f$ is the frequency of the adaptive allele at time of sampling, $g$ is the number of generations at time of sampling since fixation, $e$ is the frequency at which selection began, and $t_d = 8000$ is the time in generations since the two populations diverged.

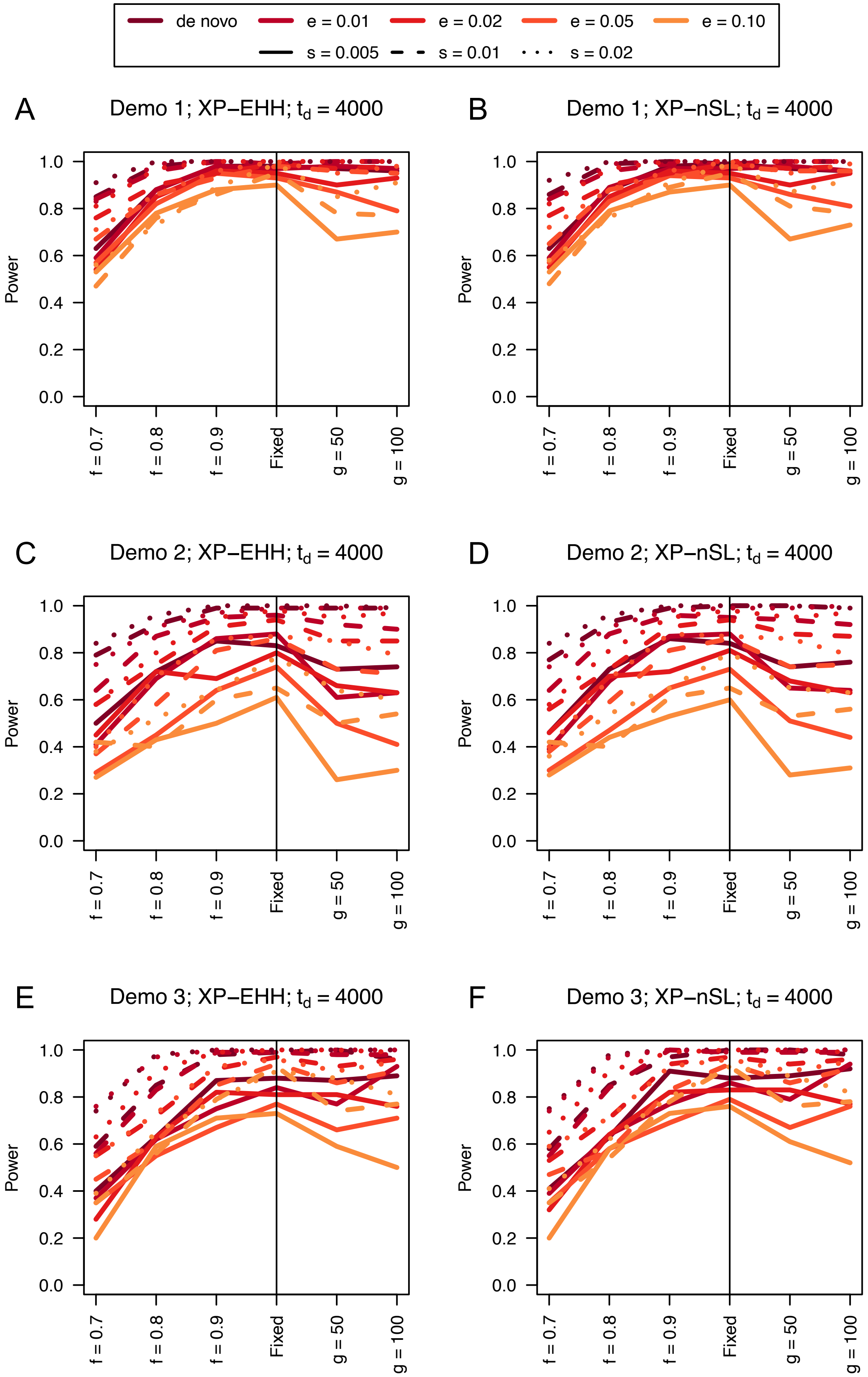
de novo
e = 0.01
e = 0.02
e = 0.05
e = 0.10
s = 0.005
s = 0.01
s = 0.02
A
Demo 1; XP−EHH; $t_d$ = 4000
B
Demo 1; XP−nSL; $t_d$ = 4000
C
Demo 2; XP−EHH; $t_d$ = 4000
D
Demo 2; XP−nSL; $t_d$ = 4000
E
Demo 3; XP−EHH; $t_d$ = 4000
F
Demo 3; XP−nSL; $t_d$ = 4000
Power
0.0
0.2
0.4
0.6
0.8
1.0
f = 0.7
f = 0.8
f = 0.9
Fixed
g = 50
g = 100

**Figure S45**. Power difference between unphased implementations of XP-EHH (A, C, and E) and XP-nSL (B, D, and F) and phased implementations. Blue curves represent the power difference between the unphased and phased statistics when applied to unphased data (UN). Red curves represent the power difference between the unphased and phased statistics when applied to perfectly phased data (PH). Values greater than 0 indicate the unphased statistic had higher power. Applied to demographic histories Demo 1 (A and B), Demo 2 (C and D), and Demo 3 (E and F) with n = 20 diploid samples from each population. $s$ is the selection coefficient, $f$ is the frequency of the adaptive allele at time of sampling, $g$ is the number of generations at time of sampling since fixation, $e$ is the frequency at which selection began, and $t_d = 4000$ is the time in generations since the two populations diverged.

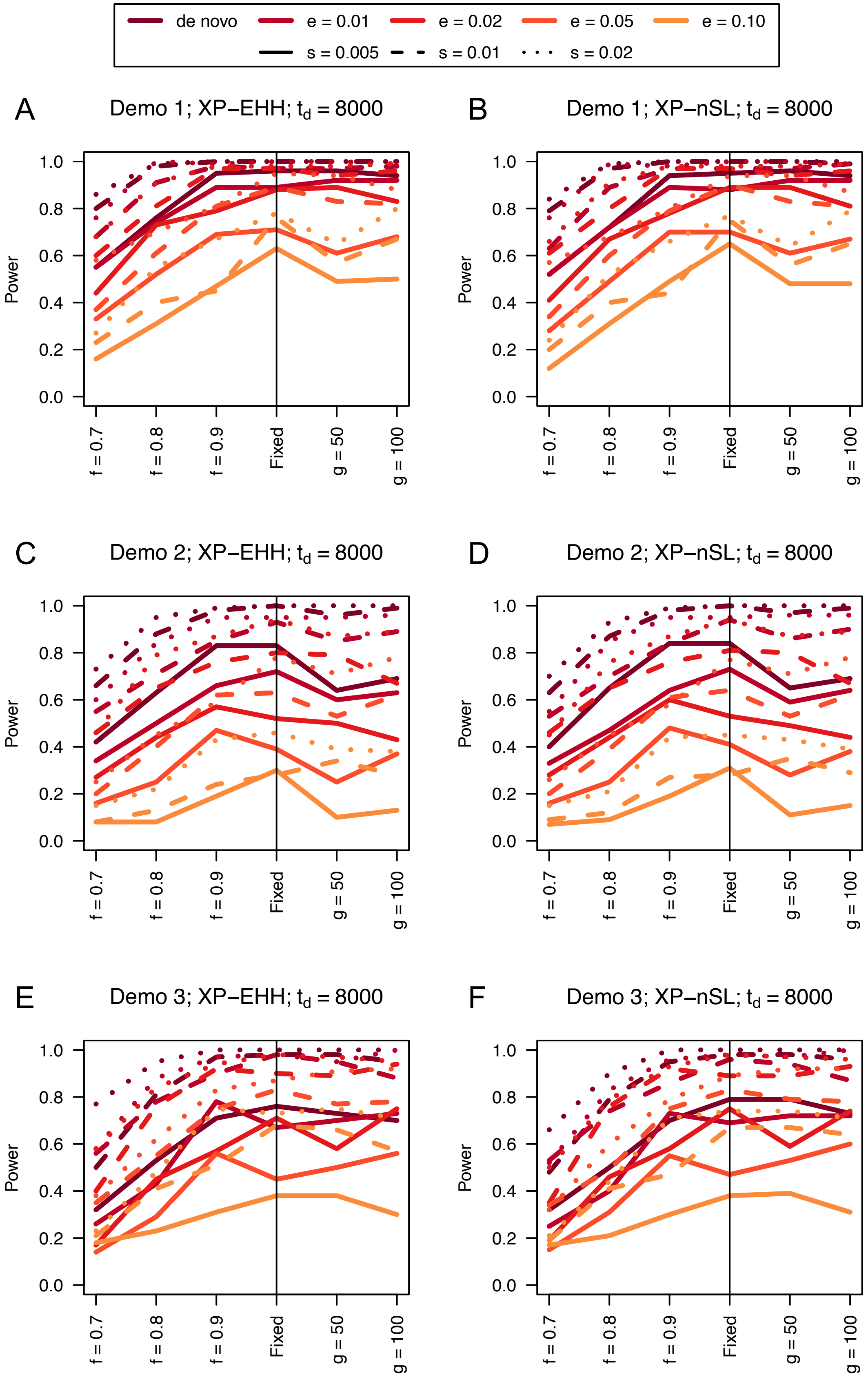

de novo
e = 0.01
e = 0.02
e = 0.05
e = 0.10
s = 0.005
s = 0.01
s = 0.02
A
Demo 1; XP−EHH; $t_d$ = 8000
B
Demo 1; XP−nSL; $t_d$ = 8000
C
Demo 2; XP−EHH; $t_d$ = 8000
D
Demo 2; XP−nSL; $t_d$ = 8000
E
Demo 3; XP−EHH; $t_d$ = 8000
F
Demo 3; XP−nSL; $t_d$ = 8000
Power
0.0
0.2
0.4
0.6
0.8
1.0
f = 0.7
f = 0.8
f = 0.9
Fixed
g = 50
g = 100

**Figure S46**. Power difference between unphased implementations of XP-EHH (A, C, and E) and XP-nSL (B, D, and F) and phased implementations. Blue curves represent the power difference between the unphased and phased statistics when applied to unphased data (UN). Red curves represent the power difference between the unphased and phased statistics when applied to perfectly phased data (PH). Values greater than 0 indicate the unphased statistic had higher power. Applied to demographic histories Demo 1 (A and B), Demo 2 (C and D), and Demo 3 (E and F) with n = 20 diploid samples from each population. $s$ is the selection coefficient, $f$ is the frequency of the adaptive allele at time of sampling, $g$ is the number of generations at time of sampling since fixation, $e$ is the frequency at which selection began, and $t_d = 8000$ is the time in generations since the two populations diverged.

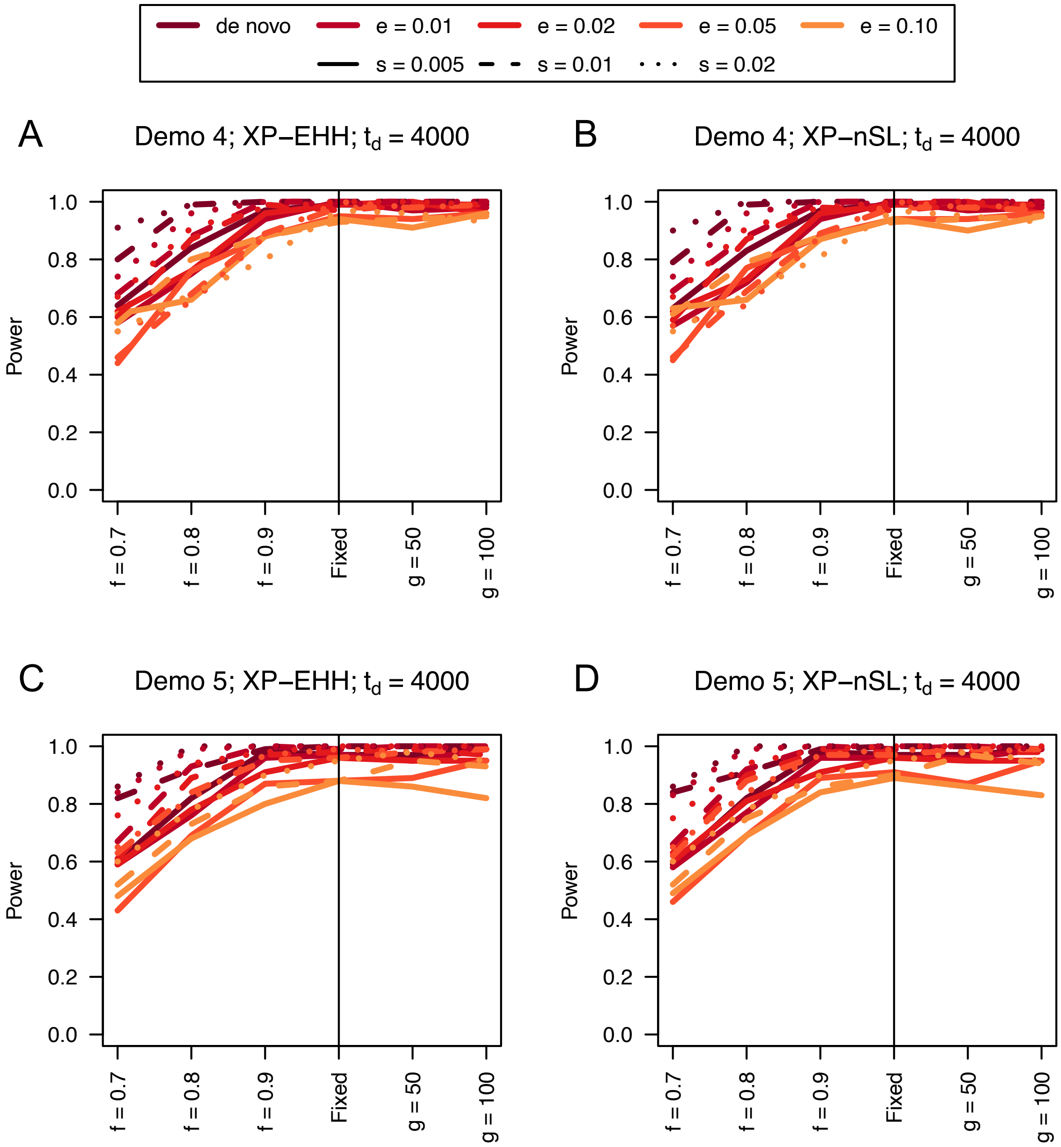


**Figure S47**. Power curves for unphased implementations of XP-EHH (A and C) and XP-nSL (B and D) under demographic histories Demo 4 (A and B), and Demo 5 (C and D) with n = 20 diploid samples from each population. $s$ is the selection coefficient, $f$ is the frequency of the adaptive allele at time of sampling, $g$ is the number of generations at time of sampling since fixation, $e$ is the frequency at which selection began, and $t_d = 4000$ is the time in generations since the two populations diverged.

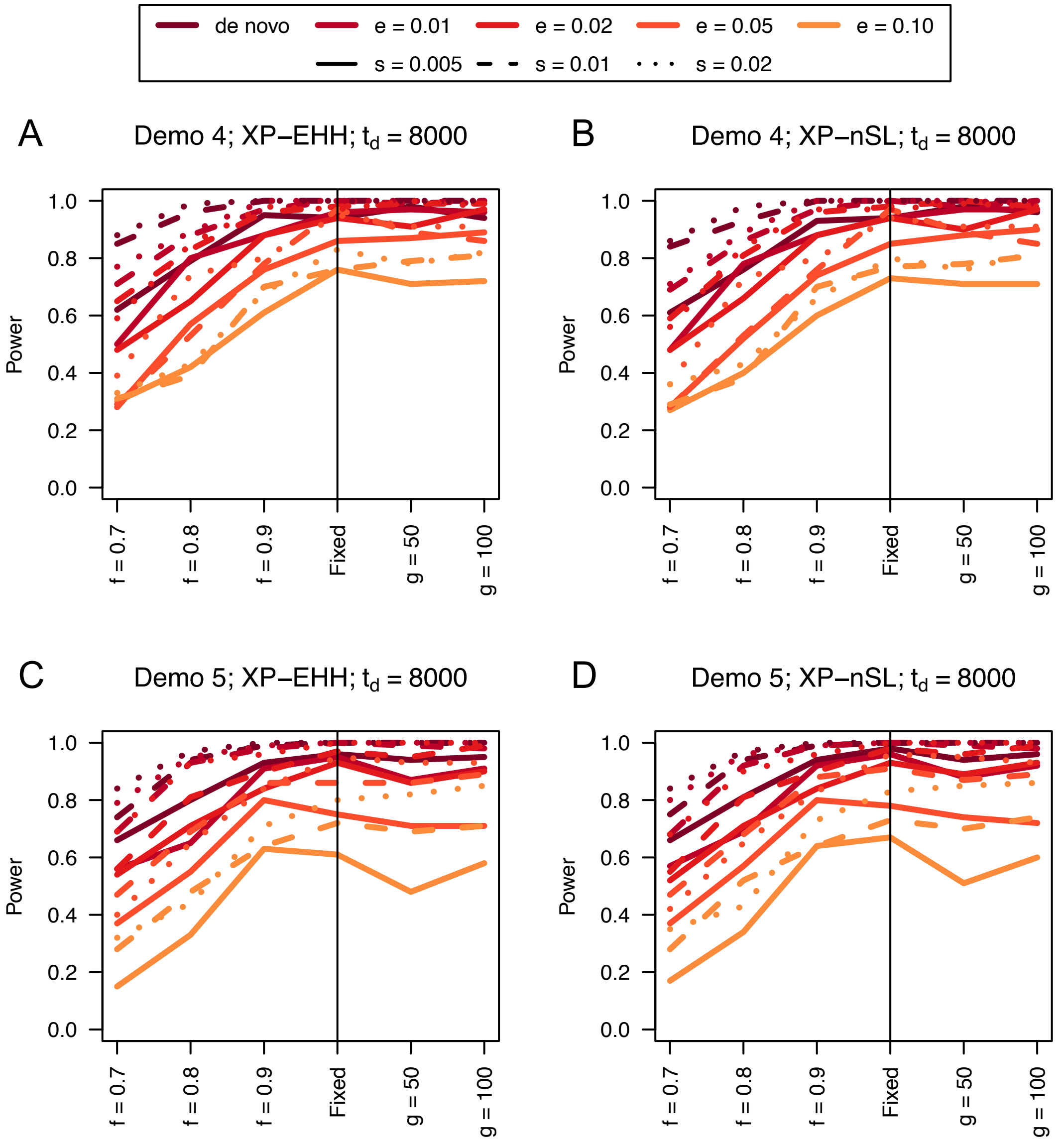


**Figure S48**. Power curves for unphased implementations of XP-EHH (A and C) and XP-nSL (B and D) under demographic histories Demo 4 (A and B), and Demo 5 (C and D) with n = 20 diploid samples from each population. $s$ is the selection coefficient, $f$ is the frequency of the adaptive allele at time of sampling, $g$ is the number of generations at time of sampling since fixation, $e$ is the frequency at which selection began, and $t_d = 8000$ is the time in generations since the two populations diverged.

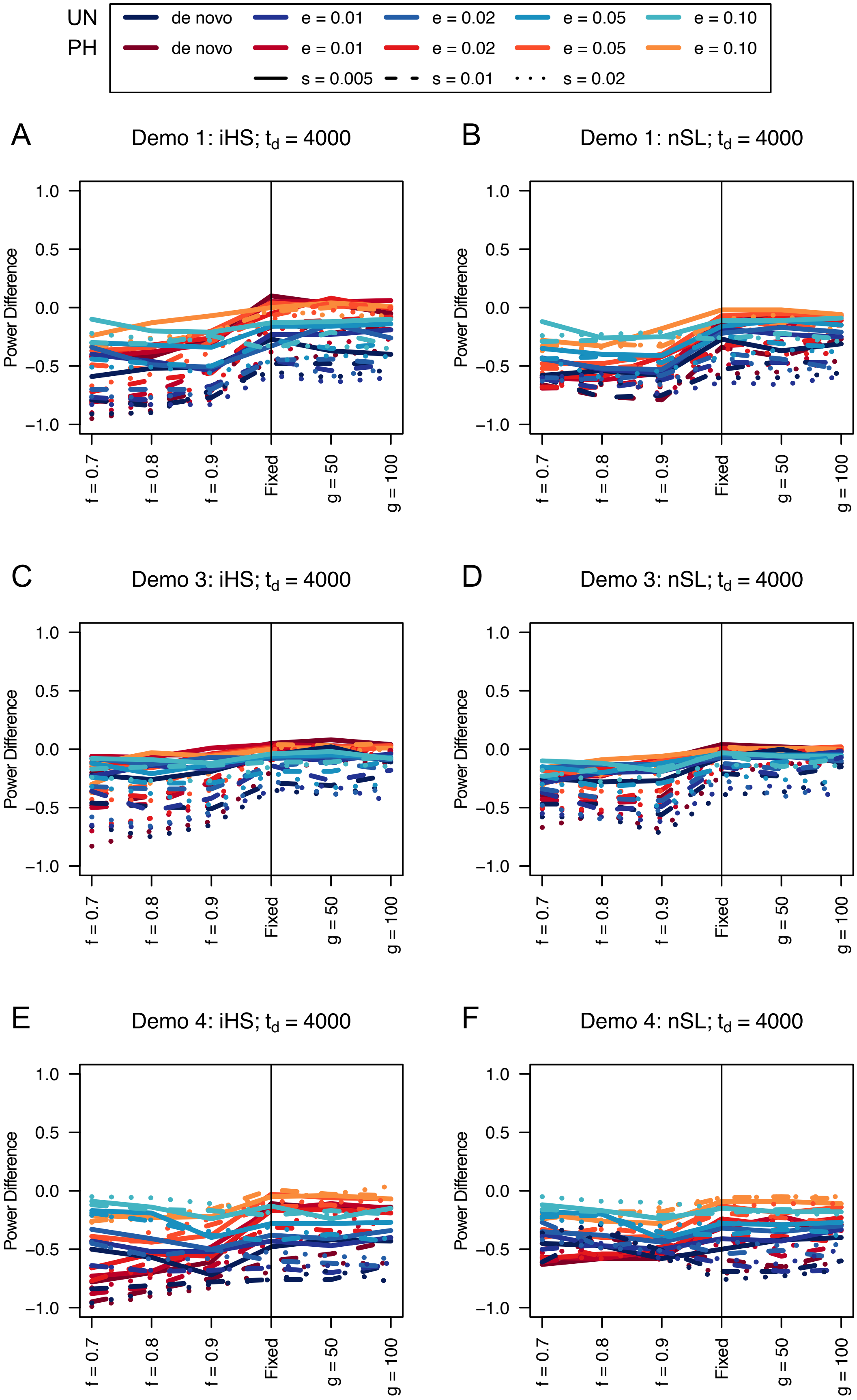
UN
PH
de novo
e = 0.01
e = 0.02
e = 0.05
e = 0.10
s = 0.005
s = 0.01
s = 0.02
A
Demo 1: iHS; $t_d$ = 4000
B
Demo 1: nSL; $t_d$ = 4000
C
Demo 3: iHS; $t_d$ = 4000
D
Demo 3: nSL; $t_d$ = 4000
E
Demo 4: iHS; $t_d$ = 4000
F
Demo 4: nSL; $t_d$ = 4000
Power Difference
f = 0.7
f = 0.8
f = 0.9
Fixed
g = 50
g = 100

**Figure S49**. Power difference between unphased implementations of iHS (A, C, and E) and nSL (B, D, and F) and phased implementations. Blue curves represent the power difference between the unphased and phased statistics when applied to unphased data (UN). Red curves represent the power difference between the unphased and phased statistics when applied to perfectly phased data (PH). Values greater than 0 indicate the unphased statistic had higher power. Applied to demographic histories Demo 1 (A and B), Demo 3 (C and D), and Demo 4 (E and F) with n = 20 diploid samples. $s$ is the selection coefficient, $f$ is the frequency of the adaptive allele at time of sampling, $g$ is the number of generations at time of sampling since fixation, $e$ is the frequency at which selection began, and $t_d = 4000$ is the time in generations since the two populations diverged.

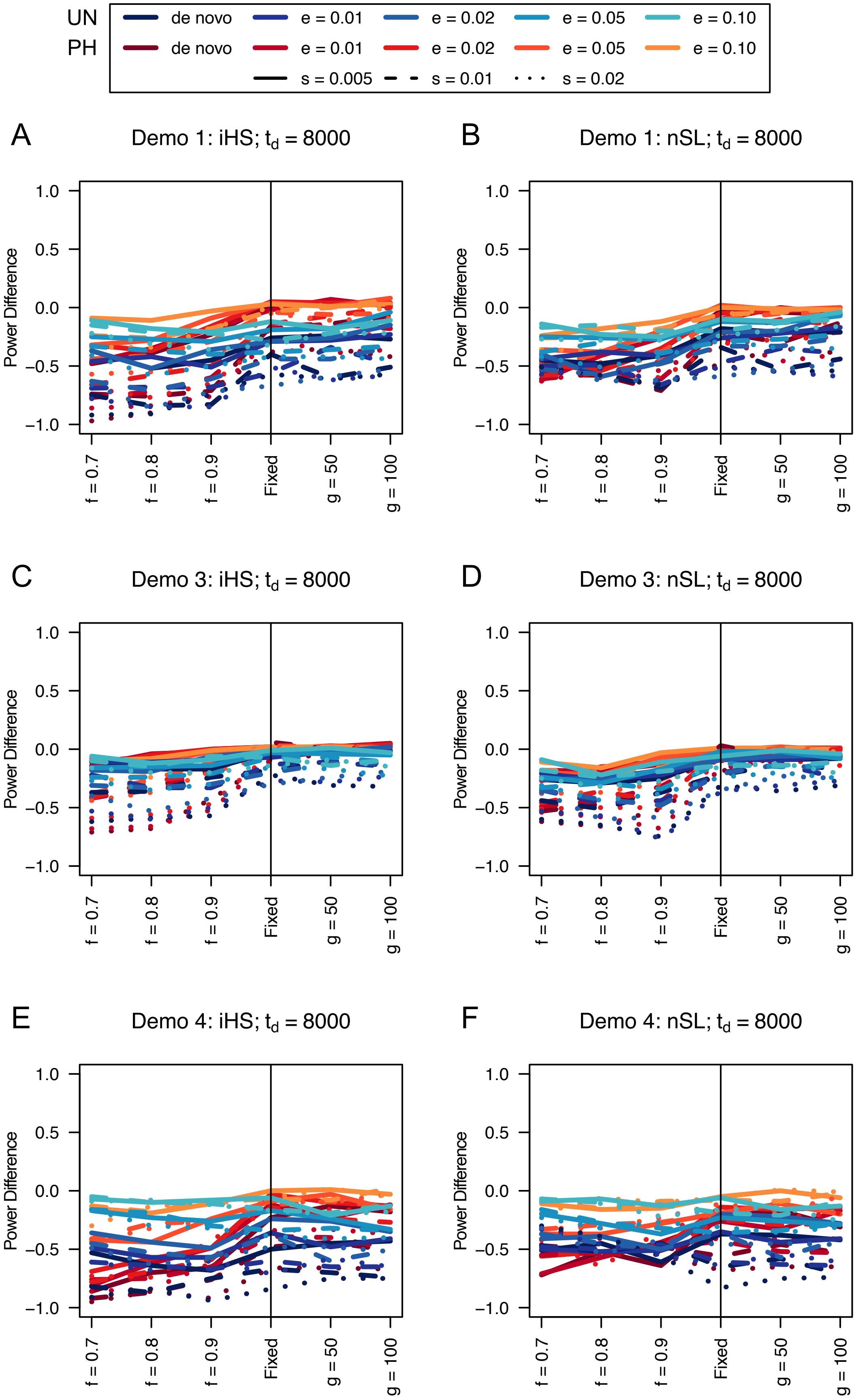

UN
PH
de novo
e = 0.01
e = 0.02
e = 0.05
e = 0.10
s = 0.005
s = 0.01
s = 0.02
A
Demo 1: iHS; $t_d$ = 8000
B
Demo 1: nSL; $t_d$ = 8000
C
Demo 3: iHS; $t_d$ = 8000
D
Demo 3: nSL; $t_d$ = 8000
E
Demo 4: iHS; $t_d$ = 8000
F
Demo 4: nSL; $t_d$ = 8000
Power Difference
1.0
0.5
0.0
−0.5
−1.0
f = 0.7
f = 0.8
f = 0.9
Fixed
g = 50
g = 100


547

**Figure S50**. Power difference between unphased implementations of iHS (A, C, and E) and nSL (B, D, and F) and phased implementations. Blue curves represent the power difference between the unphased and phased statistics when applied to unphased data (UN). Red curves represent the power difference between the unphased and phased statistics when applied to perfectly phased data (PH). Values greater than 0 indicate the unphased statistic had higher power. Applied to demographic histories Demo 1 (A and B), Demo 3 (C and D), and Demo 4 (E and F) with n = 20 diploid samples. $s$ is the selection coefficient, $f$ is the frequency of the adaptive allele at time of sampling, $g$ is the number of generations at time of sampling since fixation, $e$ is the frequency at which selection began, and $t_d = 8000$ is the time in generations since the two populations diverged.

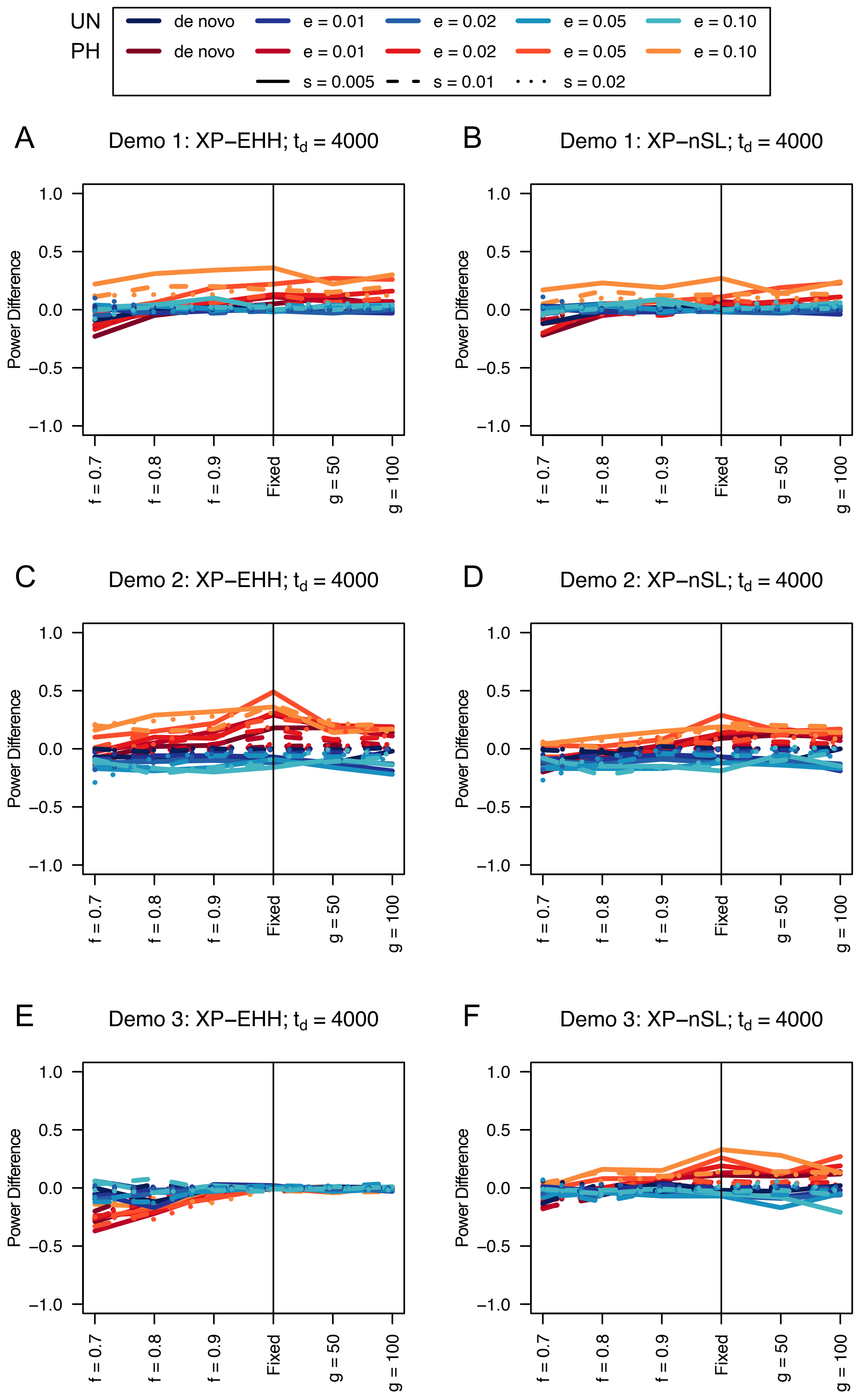
UN
PH
de novo
e = 0.01
e = 0.02
e = 0.05
e = 0.10
s = 0.005
s = 0.01
s = 0.02
A
Demo 1: XP−EHH; $t_d$ = 4000
B
Demo 1: XP−nSL; $t_d$ = 4000
C
Demo 2: XP−EHH; $t_d$ = 4000
D
Demo 2: XP−nSL; $t_d$ = 4000
E
Demo 3: XP−EHH; $t_d$ = 4000
F
Demo 3: XP−nSL; $t_d$ = 4000
Power Difference
1.0
0.5
0.0
−0.5
−1.0
f = 0.7
f = 0.8
f = 0.9
Fixed
g = 50
g = 100

557

**Figure S51**. Power difference between unphased implementations of XP-EHH (A, C, and E) and XP-nSL (B, D, and F) and phased implementations. Blue curves represent the power difference between the unphased and phased statistics when applied to unphased data (UN). Red curves represent the power difference between the unphased and phased statistics when applied to perfectly phased data (PH). Values greater than 0 indicate the unphased statistic had higher power. Applied to demographic histories Demo 1 (A and B), Demo 2 (C and D), and Demo 3 (E and F) with n = 20 diploid samples from each population. $s$ is the selection coefficient, $f$ is the frequency of the adaptive allele at time of sampling, $g$ is the number of generations at time of sampling since fixation, $e$ is the frequency at which selection began, and $t_d = 4000$ is the time in generations since the two populations diverged.

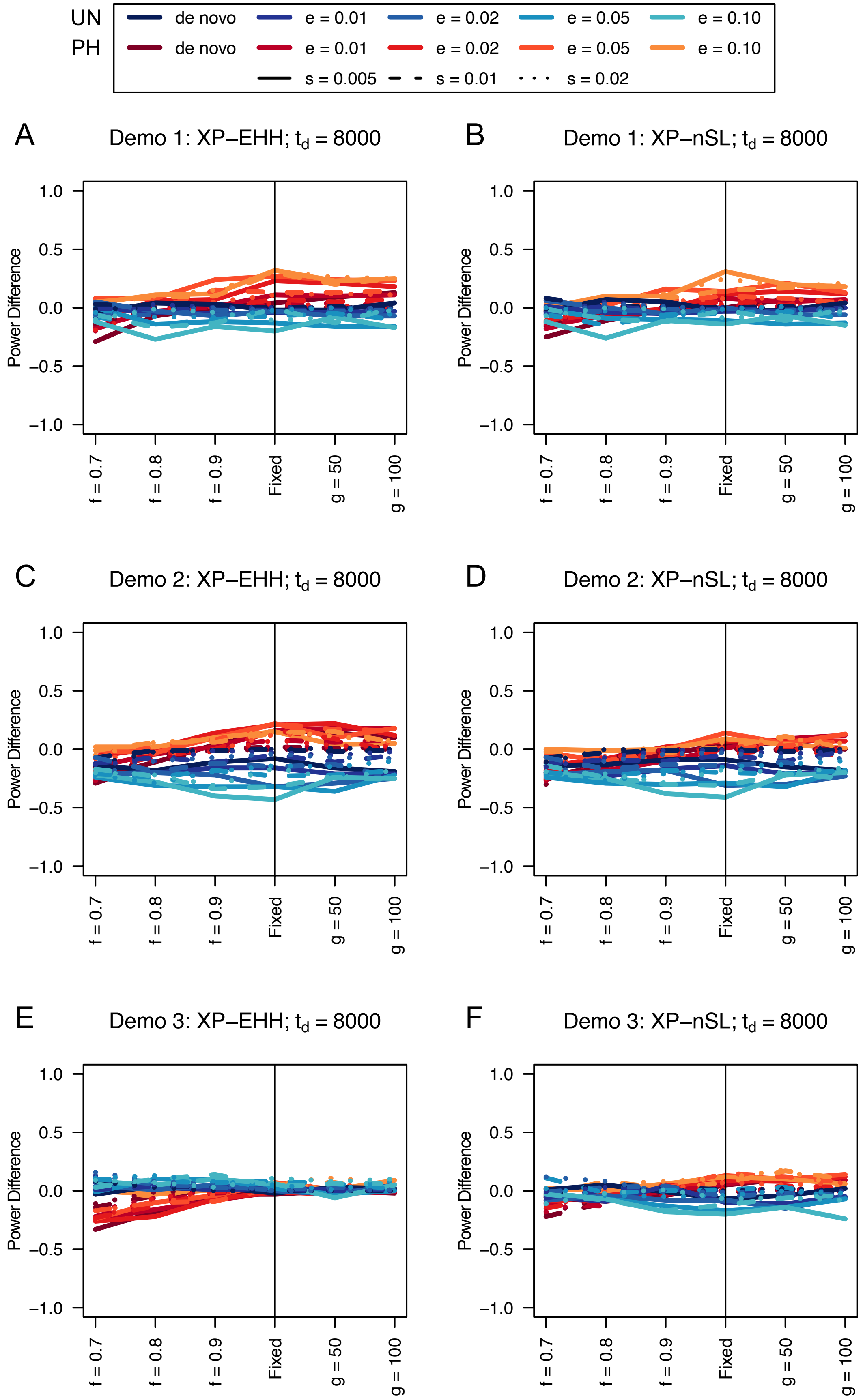

UN
PH
de novo
e = 0.01
e = 0.02
e = 0.05
e = 0.10
s = 0.005
s = 0.01
s = 0.02
A
Demo 1: XP−EHH; $t_d$ = 8000
B
Demo 1: XP−nSL; $t_d$ = 8000
C
Demo 2: XP−EHH; $t_d$ = 8000
D
Demo 2: XP−nSL; $t_d$ = 8000
E
Demo 3: XP−EHH; $t_d$ = 8000
F
Demo 3: XP−nSL; $t_d$ = 8000
Power Difference
1.0
0.5
0.0
−0.5
−1.0
f = 0.7
f = 0.8
f = 0.9
Fixed
g = 50
g = 100

**Figure S52**. Power difference between unphased implementations of XP-EHH (A, C, and E) and XP-nSL (B, D, and F) and phased implementations. Blue curves represent the power difference between the unphased and phased statistics when applied to unphased data (UN). Red curves represent the power difference between the unphased and phased statistics when applied to perfectly phased data (PH). Values greater than 0 indicate the unphased statistic had higher power. Applied to demographic histories Demo 1 (A and B), Demo 2 (C and D), and Demo 3 (E and F) with n = 20 diploid samples from each population. $s$ is the selection coefficient, $f$ is the frequency of the adaptive allele at time of sampling, $g$ is the number of generations at time of sampling since fixation, $e$ is the frequency at which selection began, and $t_d = 8000$ is the time in generations since the two populations diverged.

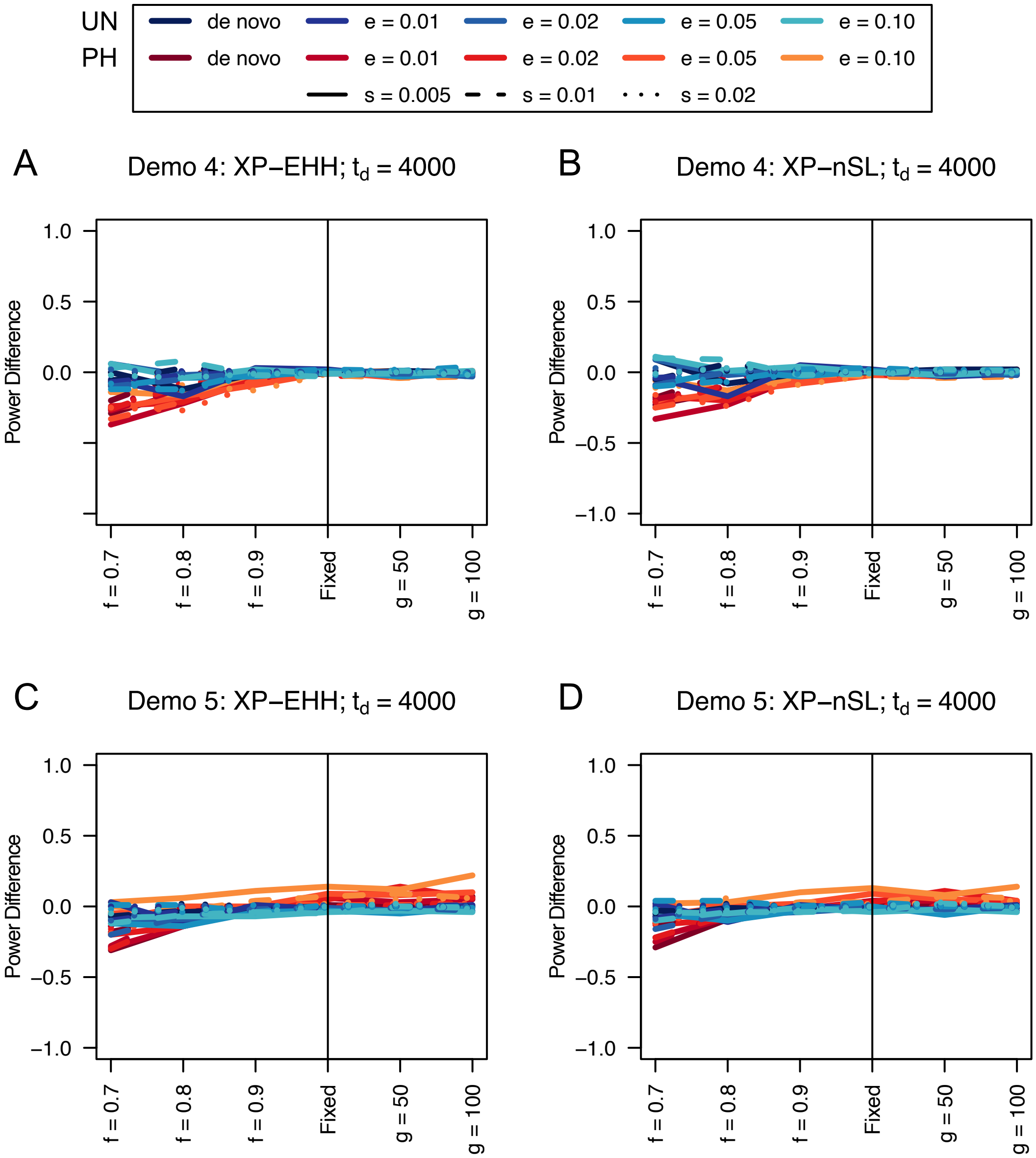


**Figure S53**. Power difference between unphased implementations of XP-EHH (A and C) and XP-nSL (B and D) and phased implementations. Blue curves represent the power difference between the unphased and phased statistics when applied to unphased data (UN). Red curves represent the power difference between the unphased and phased statistics when applied to perfectly phased data (PH). Values greater than 0 indicate the unphased statistic had higher power. Applied to demographic histories Demo 4 (A and B), and Demo 5 (C and D) with n = 20 diploid samples from each population. $s$ is the selection coefficient, $f$ is the frequency of the adaptive allele at time of sampling, $g$ is the number of generations at time of sampling since fixation, $e$ is the frequency at which selection began, and $t_d = 4000$ is the time in generations since the two populations diverged.

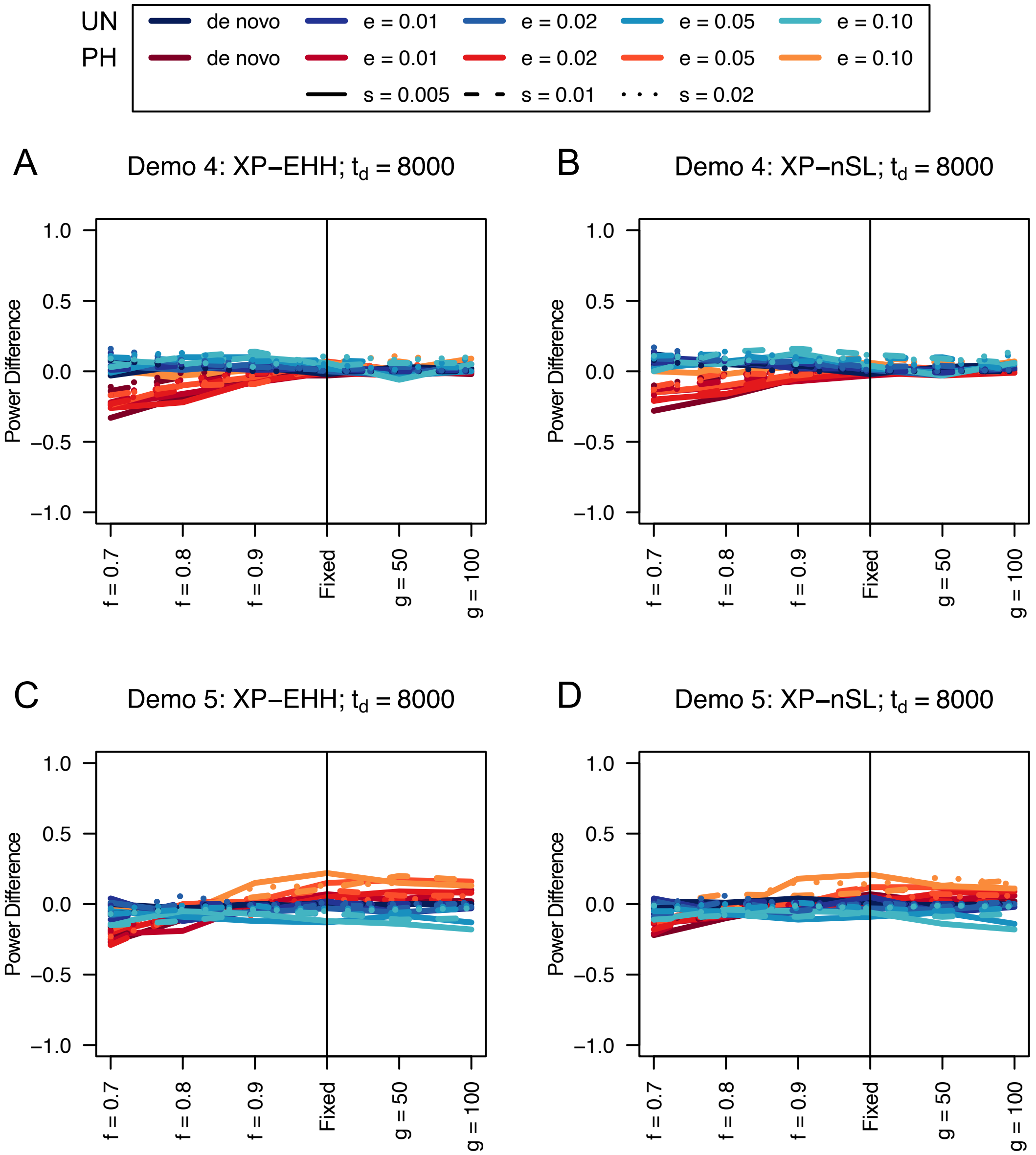


**Table S54**. Power difference between unphased implementations of XP-EHH (A and C) and XP-nSL (B and D) and phased implementations. Blue curves represent the power difference between the unphased and phased statistics when applied to unphased data (UN). Red curves represent the power difference between the unphased and phased statistics when applied to perfectly phased data (PH). Values greater than 0 indicate the unphased statistic had higher power. Applied to demographic histories Demo 4 (A and B), and Demo 5 (C and D) with n = 20 diploid samples from each population. $s$ is the selection coefficient, $f$ is the frequency of the adaptive allele at time of sampling, $g$ is the number of generations at time of sampling since fixation, $e$ is the frequency at which selection began, and $t_d = 8000$ is the time in generations since the two populations diverged.

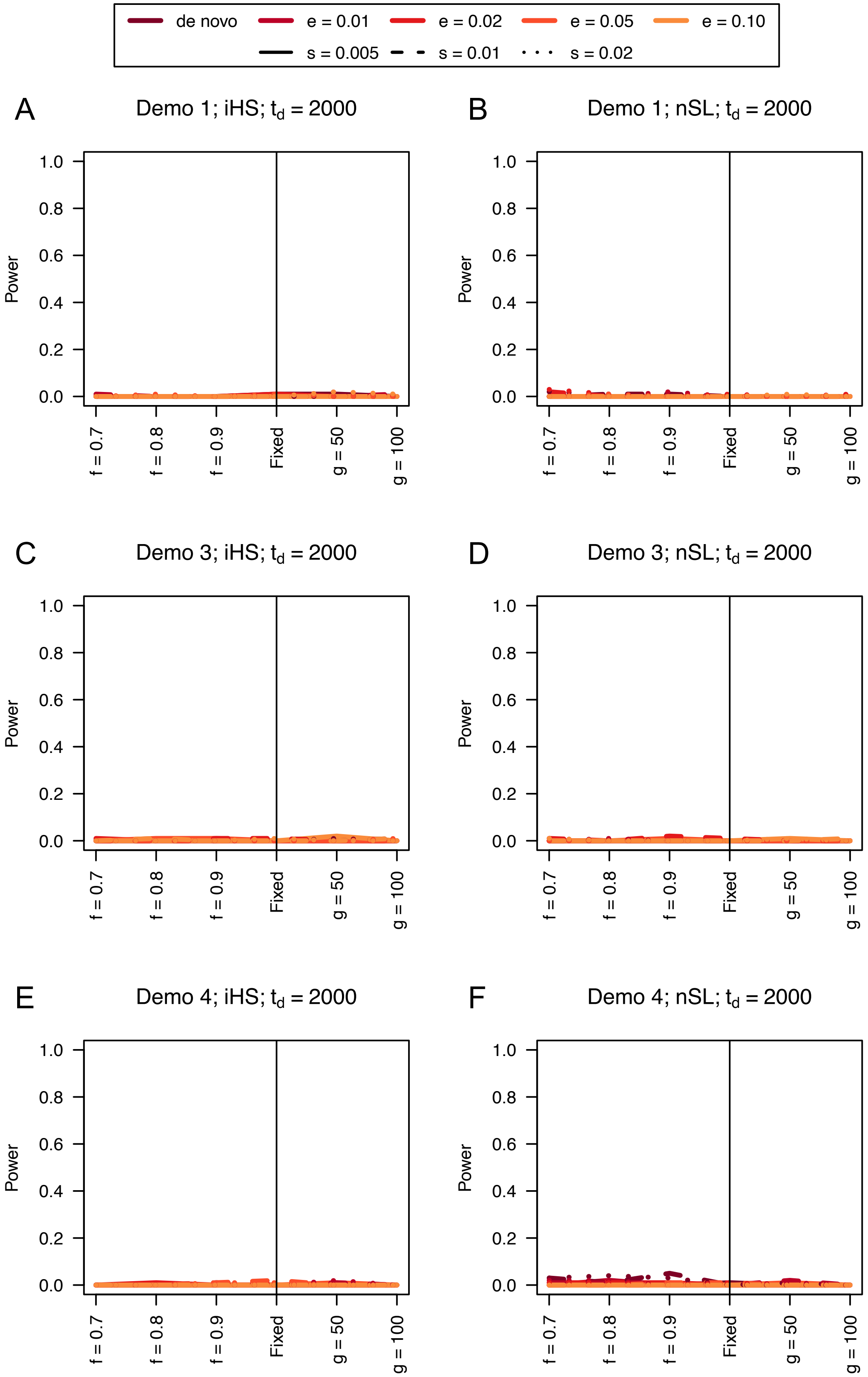

de novo
e = 0.01
e = 0.02
e = 0.05
e = 0.10
s = 0.005
s = 0.01
s = 0.02
A
Demo 1; iHS; $t_d$ = 2000
B
Demo 1; nSL; $t_d$ = 2000
C
Demo 3; iHS; $t_d$ = 2000
D
Demo 3; nSL; $t_d$ = 2000
E
Demo 4; iHS; $t_d$ = 2000
F
Demo 4; nSL; $t_d$ = 2000
Power
0.0
0.2
0.4
0.6
0.8
1.0
f = 0.7
f = 0.8
f = 0.9
Fixed
g = 50
g = 100

**Figure S55**. Power curves for unphased implementations of iHS (A, C, and E) and nSL (B, D, and F) under demographic histories Demo 1 (A and B), Demo 3 (C and D), and Demo 4 (E and F) with n = 10 diploid samples. $s$ is the selection coefficient, $f$ is the frequency of the adaptive allele at time of sampling, $g$ is the number of generations at time of sampling since fixation, $e$ is the frequency at which selection began, and $t_d = 2000$ is the time in generations since the two populations diverged.

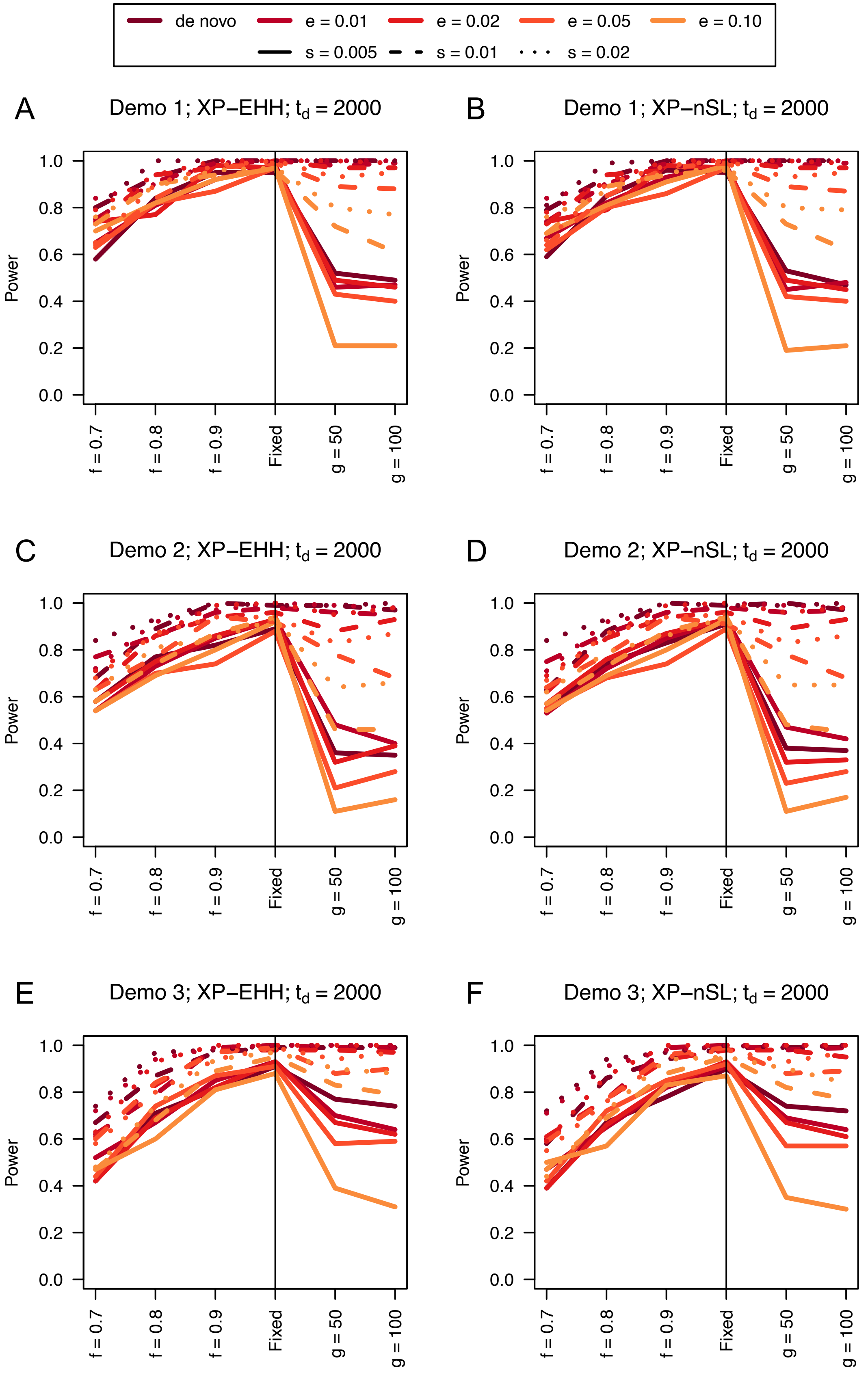
de novo
e = 0.01
e = 0.02
e = 0.05
e = 0.10
s = 0.005
s = 0.01
s = 0.02
A
Demo 1; XP−EHH; $t_d$ = 2000
B
Demo 1; XP−nSL; $t_d$ = 2000
C
Demo 2; XP−EHH; $t_d$ = 2000
D
Demo 2; XP−nSL; $t_d$ = 2000
E
Demo 3; XP−EHH; $t_d$ = 2000
F
Demo 3; XP−nSL; $t_d$ = 2000
Power
1.0
0.8
0.6
0.4
0.2
0.0
f = 0.7
f = 0.8
f = 0.9
Fixed
g = 50
g = 100

611

**Figure S56**. Power curves for unphased implementations of XP-EHH (A, C, and E) and XP-nSL (B, D, and F) under demographic histories Demo 1 (A and B), Demo 2 (C and D), and Demo 3 (E and F) with n = 10 diploid samples from each population. $s$ is the selection coefficient, $f$ is the frequency of the adaptive allele at time of sampling, $g$ is the number of generations at time of sampling since fixation, $e$ is the frequency at which selection began, and $t_d = 2000$ is the time in generations since the two populations diverged.

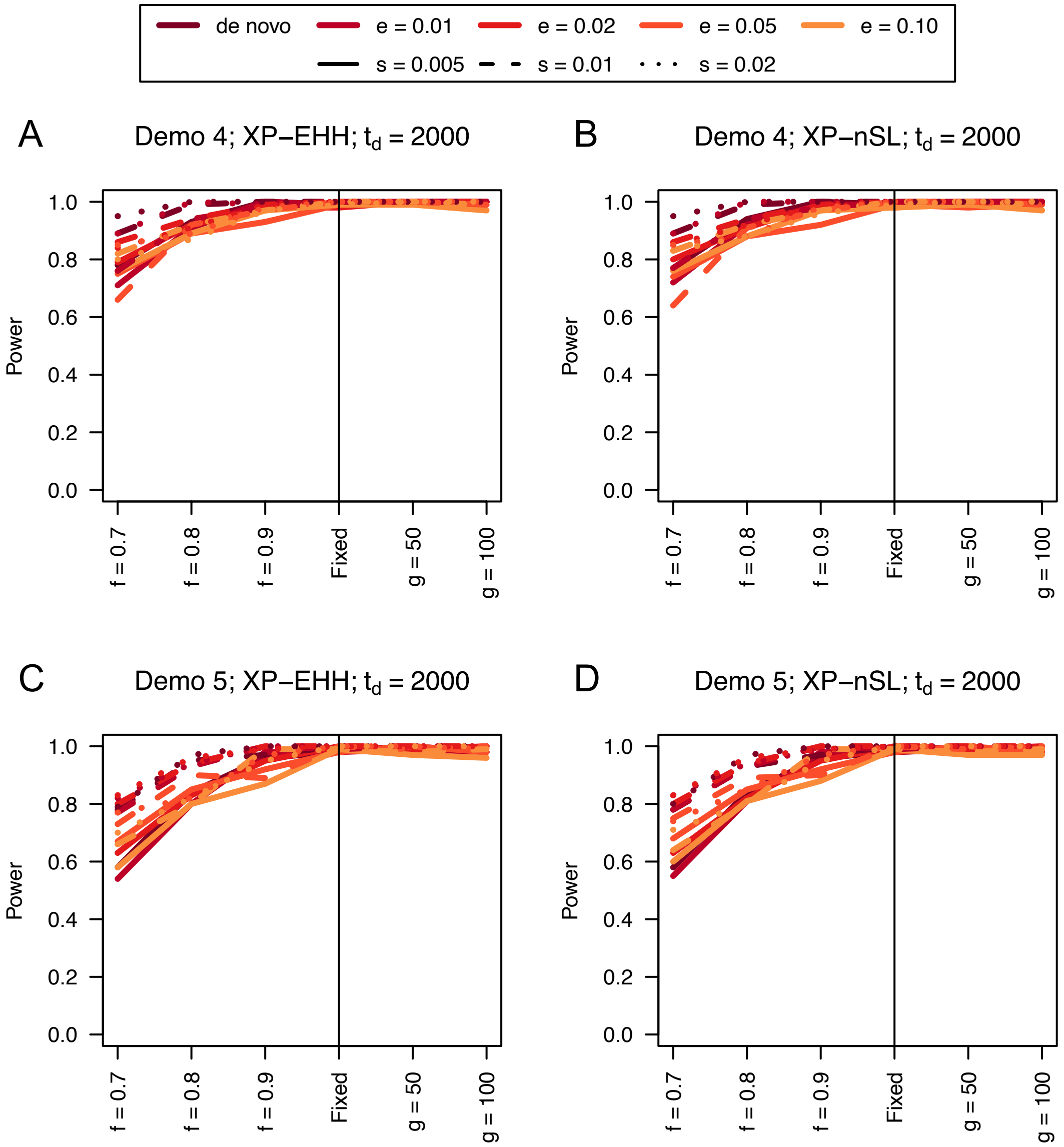


**Figure S57**. Power curves for unphased implementations of XP-EHH (A and C) and XP-nSL (B and D) under demographic histories Demo 4 (A and B), and Demo 5 (C and D) with n = 10 diploid samples from each population. $s$ is the selection coefficient, $f$ is the frequency of the adaptive allele at time of sampling, $g$ is the number of generations at time of sampling since fixation, $e$ is the frequency at which selection began, and $t_d = 2000$ is the time in generations since the two populations diverged.

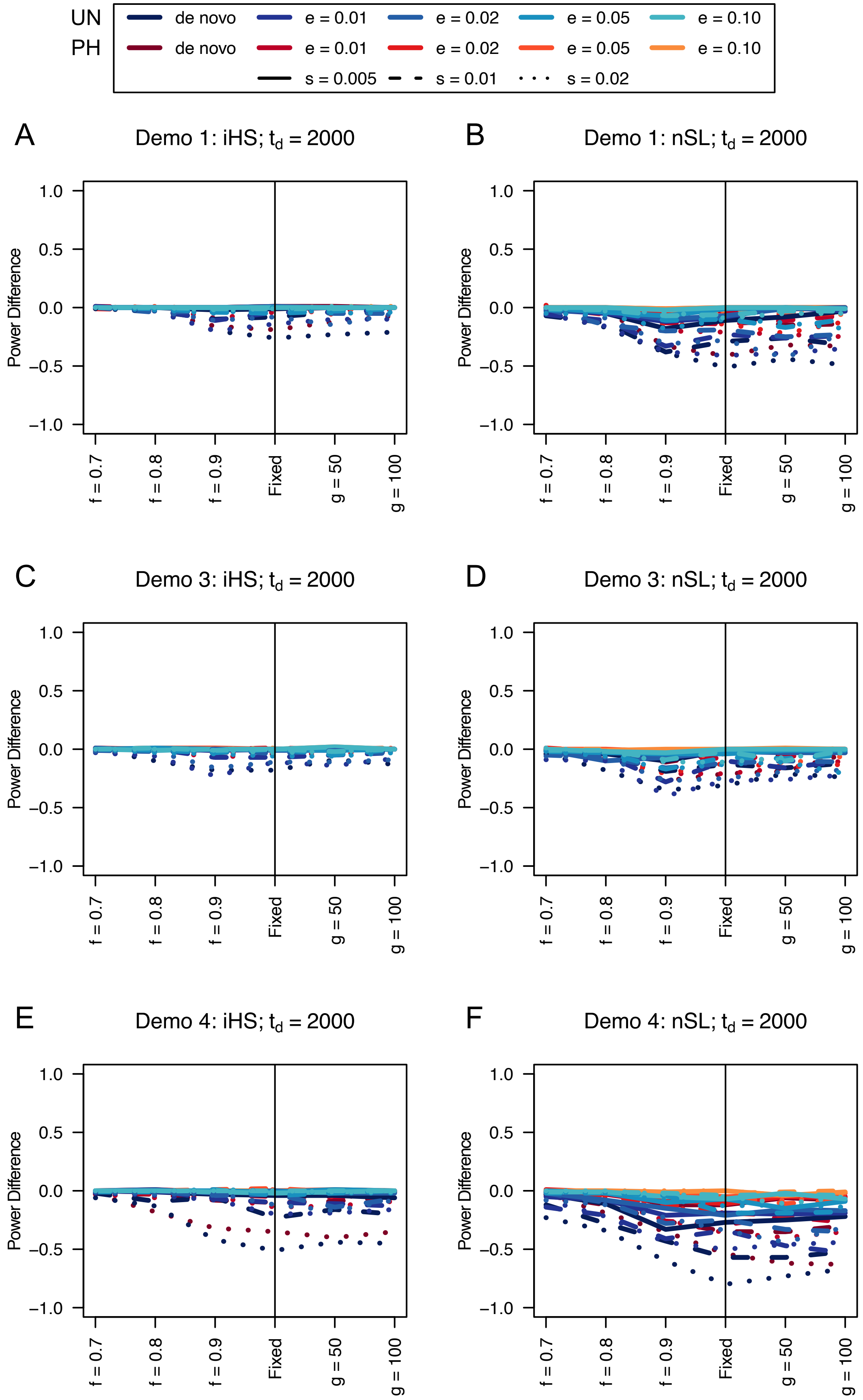

UN
PH
de novo
e = 0.01
e = 0.02
e = 0.05
e = 0.10
s = 0.005
s = 0.01
s = 0.02
A
Demo 1: iHS; $t_d$ = 2000
B
Demo 1: nSL; $t_d$ = 2000
C
Demo 3: iHS; $t_d$ = 2000
D
Demo 3: nSL; $t_d$ = 2000
E
Demo 4: iHS; $t_d$ = 2000
F
Demo 4: nSL; $t_d$ = 2000
Power Difference
1.0
0.5
0.0
−0.5
−1.0
f = 0.7
f = 0.8
f = 0.9
Fixed
g = 50
g = 100


636

**Figure S58**. Power difference between unphased implementations of iHS (A, C, and E) and nSL (B, D, and F) and phased implementations. Blue curves represent the power difference between the unphased and phased statistics when applied to unphased data (UN). Red curves represent the power difference between the unphased and phased statistics when applied to perfectly phased data (PH). Values greater than 0 indicate the unphased statistic had higher power. Applied to demographic histories Demo 1 (A and B), Demo 3 (C and D), and Demo 4 (E and F) with n = 10 diploid samples. $s$ is the selection coefficient, $f$ is the frequency of the adaptive allele at time of sampling, $g$ is the number of generations at time of sampling since fixation, $e$ is the frequency at which selection began, and $t_d = 2000$ is the time in generations since the two populations diverged.

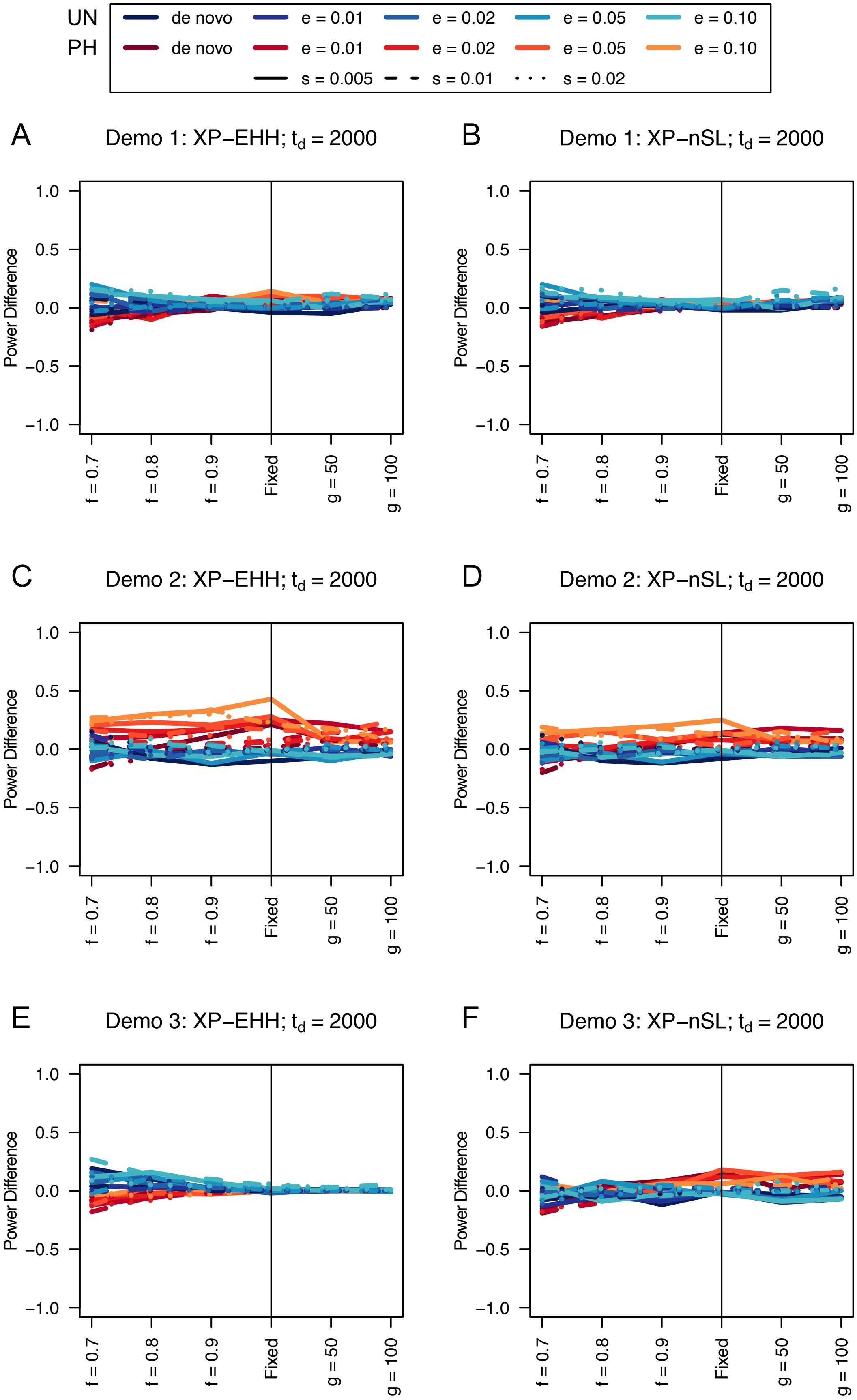

UN
PH
de novo
e = 0.01
e = 0.02
e = 0.05
e = 0.10
de novo
e = 0.01
e = 0.02
e = 0.05
e = 0.10
s = 0.005
s = 0.01
s = 0.02
A
Demo 1: XP−EHH; $t_d$ = 2000
B
Demo 1: XP−nSL; $t_d$ = 2000
C
Demo 2: XP−EHH; $t_d$ = 2000
D
Demo 2: XP−nSL; $t_d$ = 2000
E
Demo 3: XP−EHH; $t_d$ = 2000
F
Demo 3: XP−nSL; $t_d$ = 2000
Power Difference
1.0
0.5
0.0
−0.5
−1.0
f = 0.7
f = 0.8
f = 0.9
Fixed
g = 50
g = 100

**Figure S59**. Power difference between unphased implementations of XP-EHH (A, C, and E) and XP-nSL (B, D, and F) and phased implementations. Blue curves represent the power difference between the unphased and phased statistics when applied to unphased data (UN). Red curves represent the power difference between the unphased and phased statistics when applied to perfectly phased data (PH). Values greater than 0 indicate the unphased statistic had higher power. Applied to demographic histories Demo 1 (A and B), Demo 2 (C and D), and Demo 3 (E and F) with n = 10 diploid samples from each population. $s$ is the selection coefficient, $f$ is the frequency of the adaptive allele at time of sampling, $g$ is the number of generations at time of sampling since fixation, $e$ is the frequency at which selection began, and $t_d = 2000$ is the time in generations since the two populations diverged.

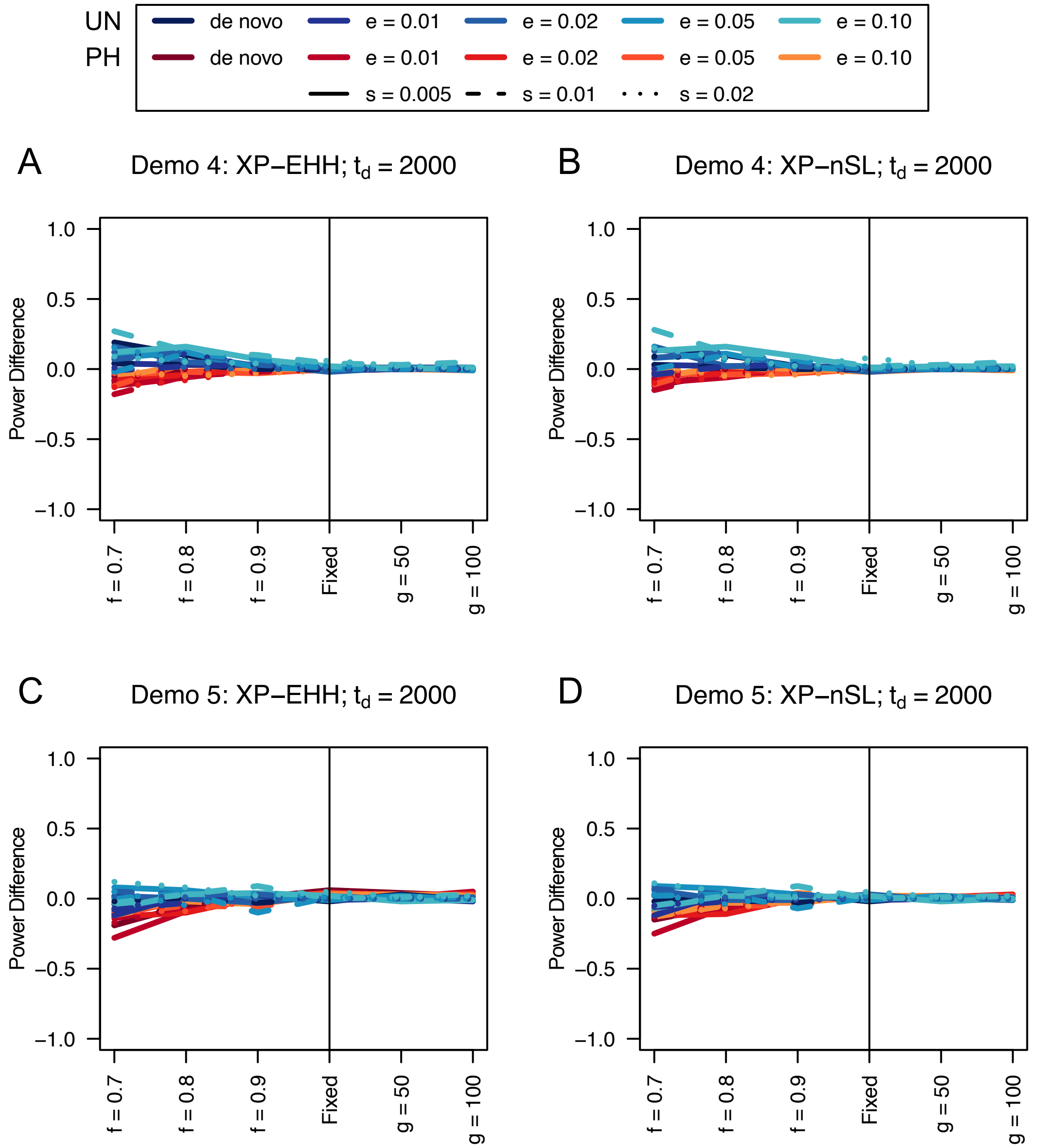


**Figure S60**. Power difference between unphased implementations of XP-EHH (A and C) and XP-nSL (B and D) and phased implementations. Blue curves represent the power difference between the unphased and phased statistics when applied to unphased data (UN). Red curves represent the power difference between the unphased and phased statistics when applied to perfectly phased data (PH). Values greater than 0 indicate the unphased statistic had higher power. Applied to demographic histories Demo 4 (A and B), and Demo 5 (C and D) with n = 10 diploid samples from each population. $s$ is the selection coefficient, $f$ is the frequency of the adaptive allele at time of sampling, $g$ is the number of generations at time of sampling since fixation, $e$ is the frequency at which selection began, and $t_d = 2000$ is the time in generations since the two populations diverged.

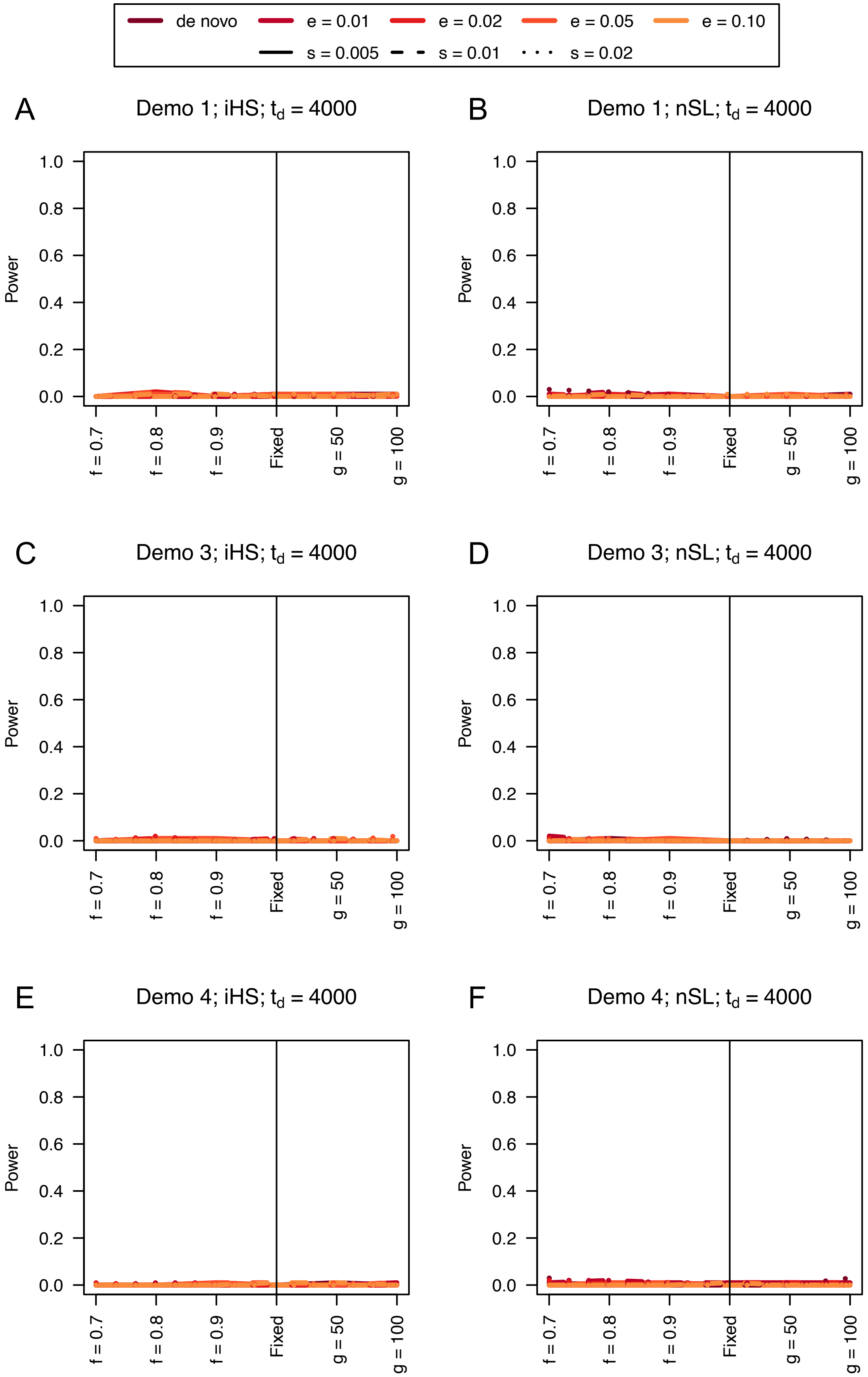
de novo
e = 0.01
e = 0.02
e = 0.05
e = 0.10
s = 0.005
s = 0.01
s = 0.02
A
Demo 1; iHS; t_d = 4000
B
Demo 1; nSL; t_d = 4000
C
Demo 3; iHS; t_d = 4000
D
Demo 3; nSL; t_d = 4000
E
Demo 4; iHS; t_d = 4000
F
Demo 4; nSL; t_d = 4000
Power
0.0
0.2
0.4
0.6
0.8
1.0
f = 0.7
f = 0.8
f = 0.9
Fixed
g = 50
g = 100

674

**Figure S61**. Power curves for unphased implementations of iHS (A, C, and E) and nSL (B, D, and F) under demographic histories Demo 1 (A and B), Demo 3 (C and D), and Demo 4 (E and F) with n = 10 diploid samples. $s$ is the selection coefficient, $f$ is the frequency of the adaptive allele at time of sampling, $g$ is the number of generations at time of sampling since fixation, $e$ is the frequency at which selection began, and $t_d = 4000$ is the time in generations since the two populations diverged.

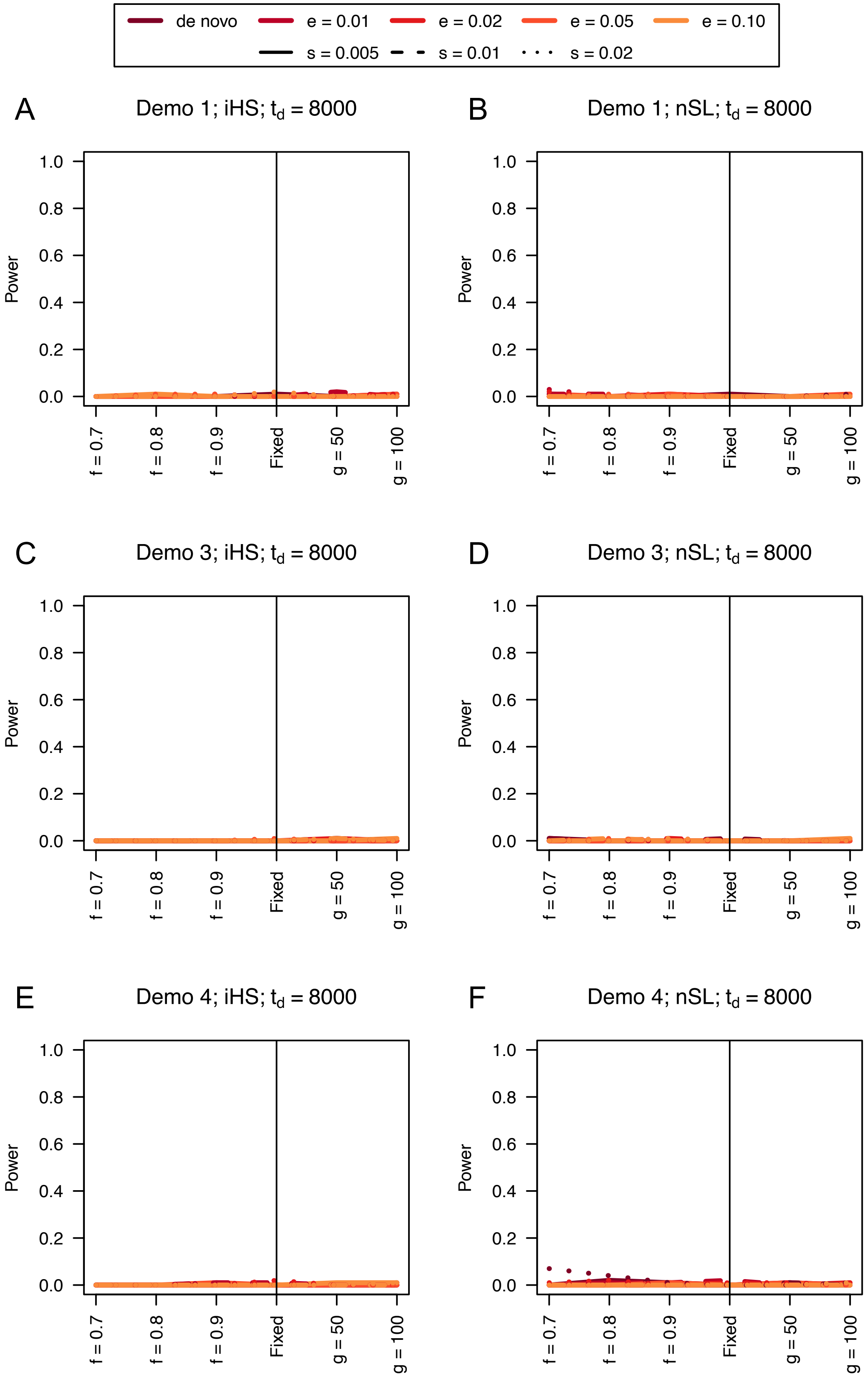
de novo
e = 0.01
e = 0.02
e = 0.05
e = 0.10
s = 0.005
s = 0.01
s = 0.02
A
Demo 1; iHS; $t_d$ = 8000
B
Demo 1; nSL; $t_d$ = 8000
C
Demo 3; iHS; $t_d$ = 8000
D
Demo 3; nSL; $t_d$ = 8000
E
Demo 4; iHS; $t_d$ = 8000
F
Demo 4; nSL; $t_d$ = 8000
Power
0.0
0.2
0.4
0.6
0.8
1.0
f = 0.7
f = 0.8
f = 0.9
Fixed
g = 50
g = 100

680

**Figure S62**. Power curves for unphased implementations of iHS (A, C, and E) and nSL (B, D, and F) under demographic histories Demo 1 (A and B), Demo 3 (C and D), and Demo 4 (E and F) with n = 10 diploid samples. $s$ is the selection coefficient, $f$ is the frequency of the adaptive allele at time of sampling, $g$ is the number of generations at time of sampling since fixation, $e$ is the frequency at which selection began, and $t_d = 8000$ is the time in generations since the two populations diverged.

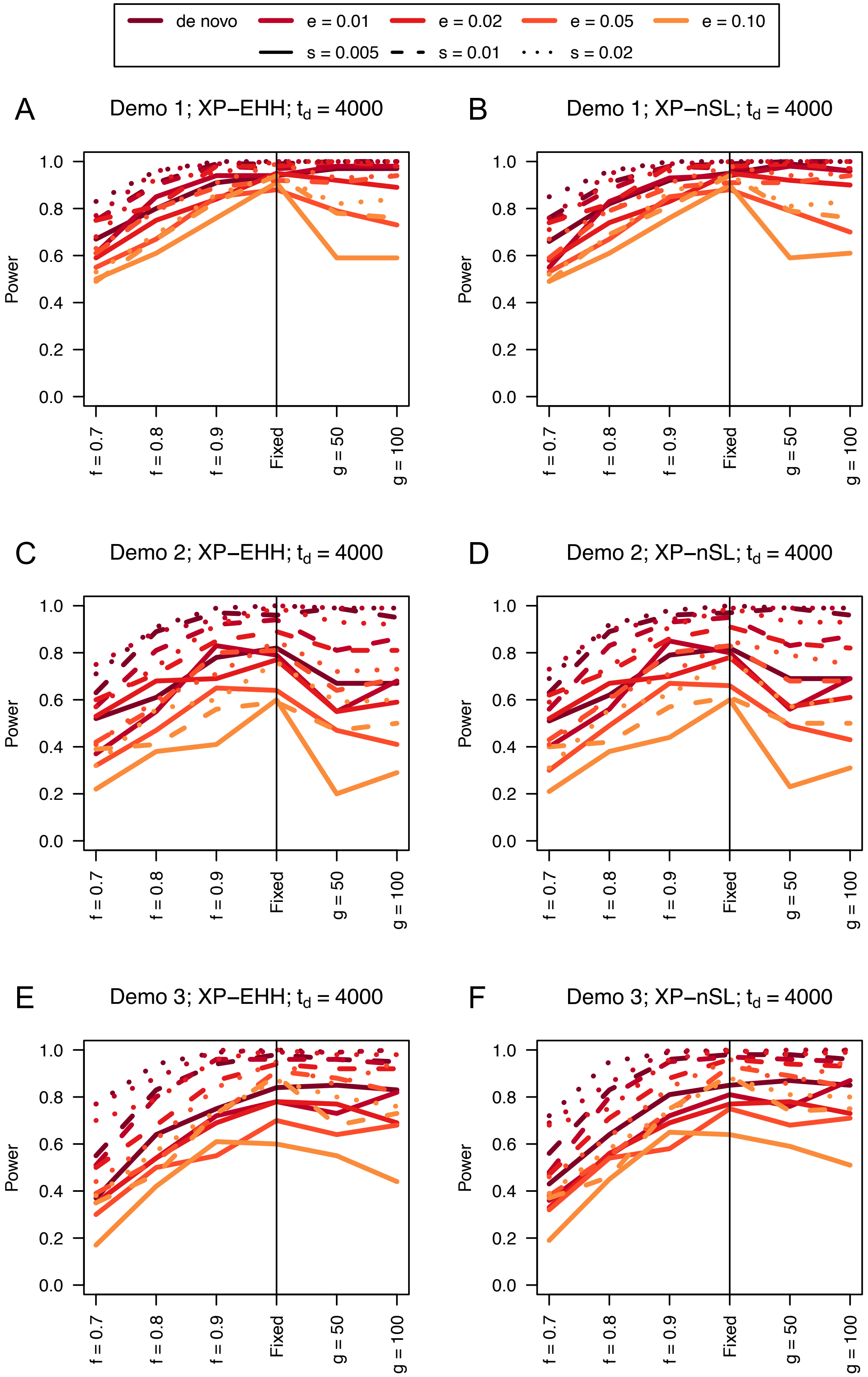

de novo
e = 0.01
e = 0.02
e = 0.05
e = 0.10
s = 0.005
s = 0.01
s = 0.02
A
Demo 1; XP−EHH; $t_d$ = 4000
B
Demo 1; XP−nSL; $t_d$ = 4000
C
Demo 2; XP−EHH; $t_d$ = 4000
D
Demo 2; XP−nSL; $t_d$ = 4000
E
Demo 3; XP−EHH; $t_d$ = 4000
F
Demo 3; XP−nSL; $t_d$ = 4000
Power
0.0
0.2
0.4
0.6
0.8
1.0
f = 0.7
f = 0.8
f = 0.9
Fixed
g = 50
g = 100

**Figure S63**. Power difference between unphased implementations of XP-EHH (A, C, and E) and XP-nSL (B, D, and F) and phased implementations. Blue curves represent the power difference between the unphased and phased statistics when applied to unphased data (UN). Red curves represent the power difference between the unphased and phased statistics when applied to perfectly phased data (PH). Values greater than 0 indicate the unphased statistic had higher power. Applied to demographic histories Demo 1 (A and B), Demo 2 (C and D), and Demo 3 (E and F) with n = 10 diploid samples from each population. $s$ is the selection coefficient, $f$ is the frequency of the adaptive allele at time of sampling, $g$ is the number of generations at time of sampling since fixation, $e$ is the frequency at which selection began, and $t_d = 4000$ is the time in generations since the two populations diverged.

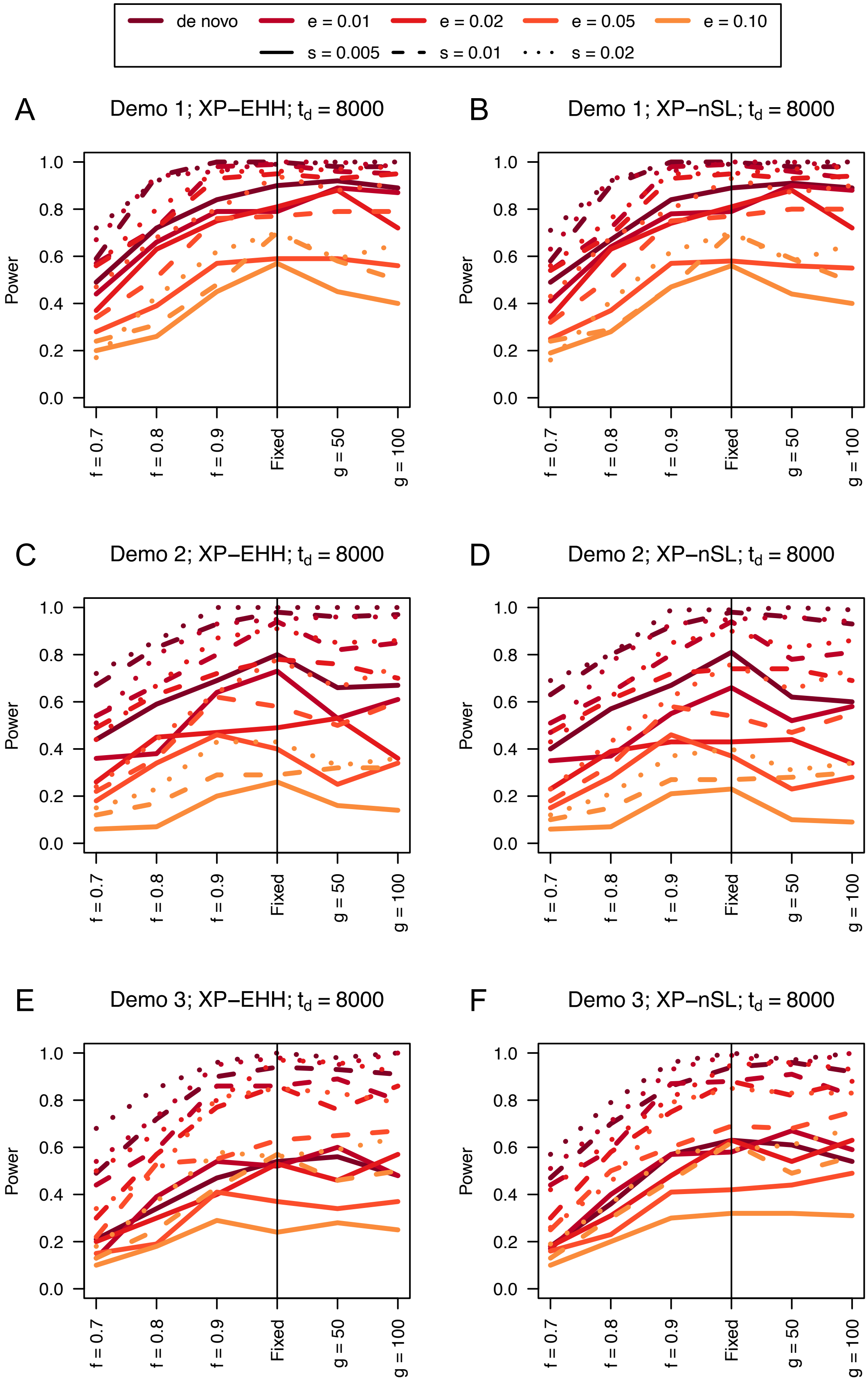

de novo
e = 0.01
e = 0.02
e = 0.05
e = 0.10
s = 0.005
s = 0.01
s = 0.02
A
Demo 1; XP−EHH; $t_d$ = 8000
B
Demo 1; XP−nSL; $t_d$ = 8000
C
Demo 2; XP−EHH; $t_d$ = 8000
D
Demo 2; XP−nSL; $t_d$ = 8000
E
Demo 3; XP−EHH; $t_d$ = 8000
F
Demo 3; XP−nSL; $t_d$ = 8000
Power
0.0
0.2
0.4
0.6
0.8
1.0
f = 0.7
f = 0.8
f = 0.9
Fixed
g = 50
g = 100

**Figure S64**. Power difference between unphased implementations of XP-EHH (A, C, and E) and XP-nSL (B, D, and F) and phased implementations. Blue curves represent the power difference between the unphased and phased statistics when applied to unphased data (UN). Red curves represent the power difference between the unphased and phased statistics when applied to perfectly phased data (PH). Values greater than 0 indicate the unphased statistic had higher power. Applied to demographic histories Demo 1 (A and B), Demo 2 (C and D), and Demo 3 (E and F) with n = 10 diploid samples from each population. $s$ is the selection coefficient, $f$ is the frequency of the adaptive allele at time of sampling, $g$ is the number of generations at time of sampling since fixation, $e$ is the frequency at which selection began, and $t_d = 8000$ is the time in generations since the two populations diverged.

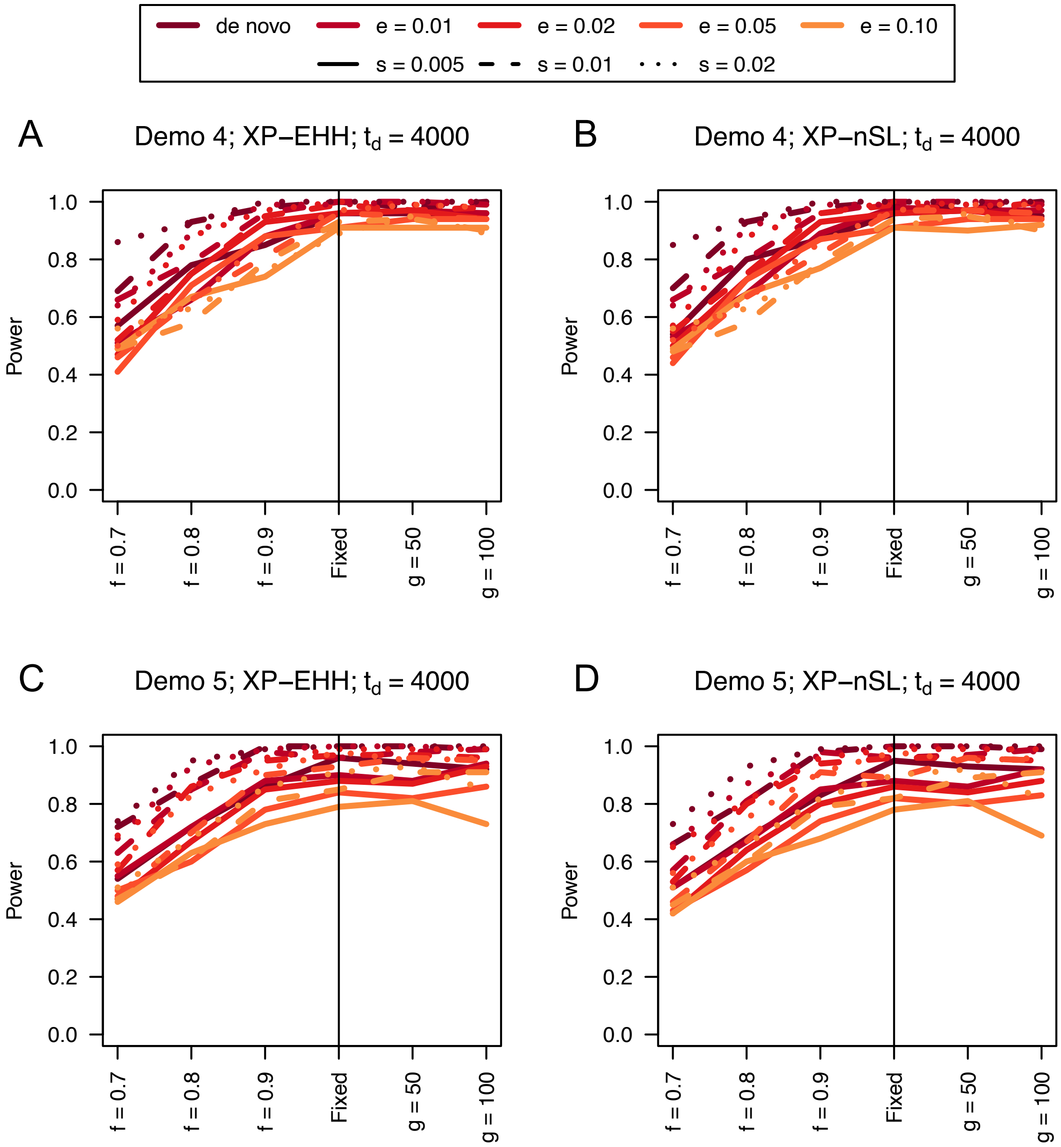


**Figure S65**. Power curves for unphased implementations of XP-EHH (A and C) and XP-nSL (B and D) under demographic histories Demo 4 (A and B), and Demo 5 (C and D) with n = 10 diploid samples from each population. $s$ is the selection coefficient, $f$ is the frequency of the adaptive allele at time of sampling, $g$ is the number of generations at time of sampling since fixation, $e$ is the frequency at which selection began, and $t_d = 4000$ is the time in generations since the two populations diverged.

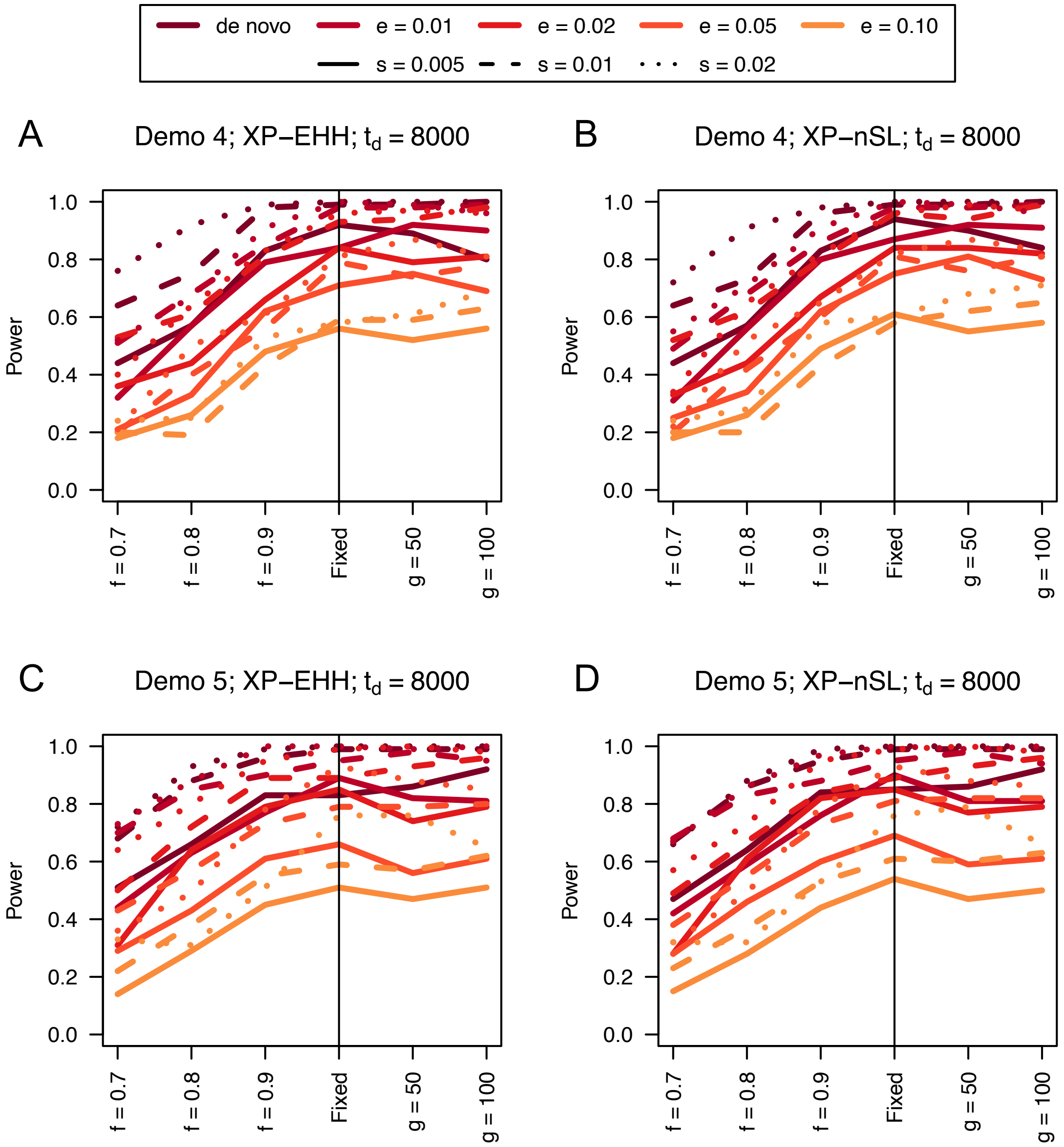


**Figure S66**. Power curves for unphased implementations of XP-EHH (A and C) and XP-nSL (B and D) under demographic histories Demo 4 (A and B), and Demo 5 (C and D) with n = 10 diploid samples from each population. $s$ is the selection coefficient, $f$ is the frequency of the adaptive allele at time of sampling, $g$ is the number of generations at time of sampling since fixation, $e$ is the frequency at which selection began, and $t_d = 8000$ is the time in generations since the two populations diverged.

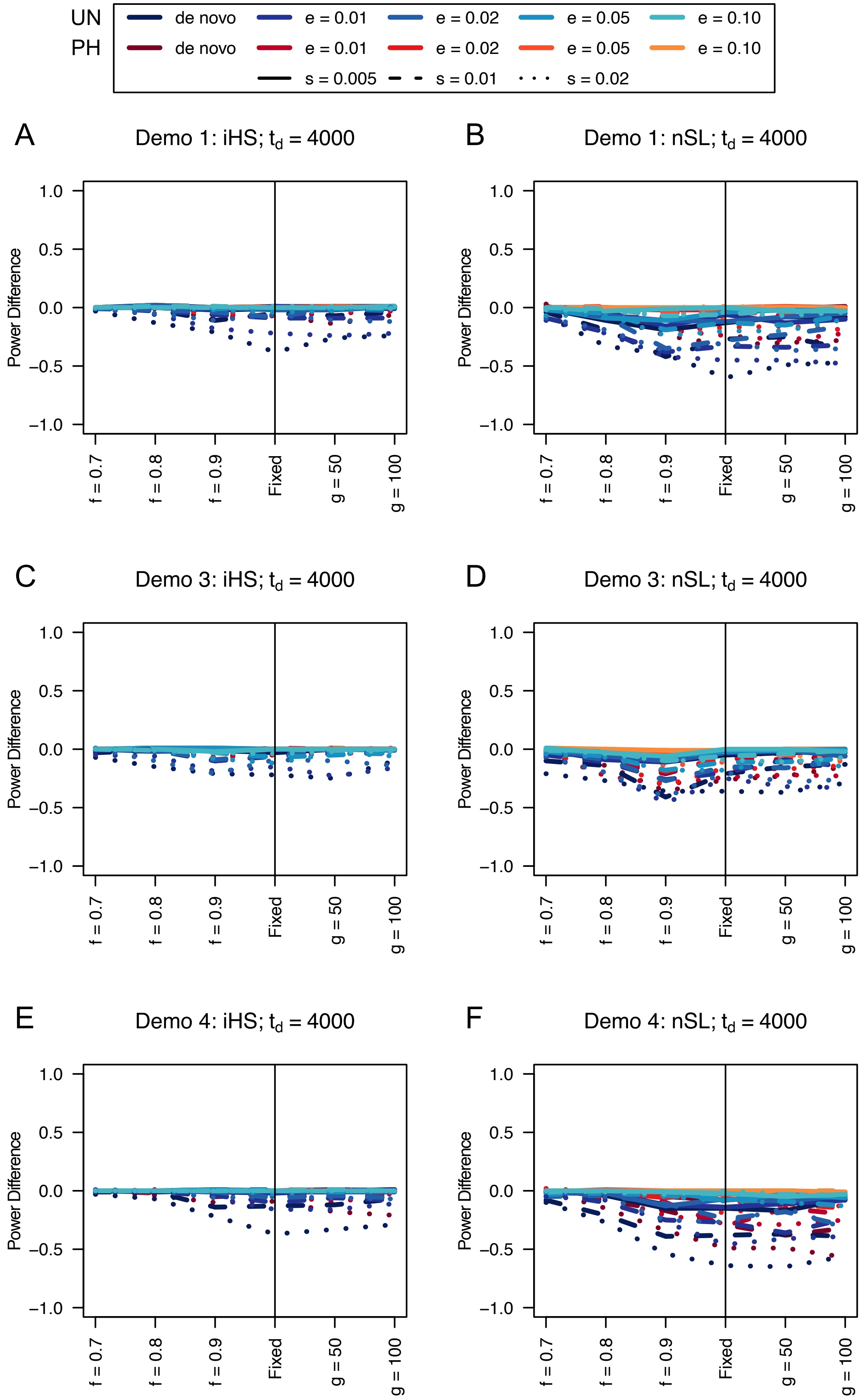

UN
PH
de novo
e = 0.01
e = 0.02
e = 0.05
e = 0.10
s = 0.005
s = 0.01
s = 0.02
A
Demo 1: iHS; t_d = 4000
B
Demo 1: nSL; t_d = 4000
C
Demo 3: iHS; t_d = 4000
D
Demo 3: nSL; t_d = 4000
E
Demo 4: iHS; t_d = 4000
F
Demo 4: nSL; t_d = 4000
Power Difference
1.0
0.5
0.0
−0.5
−1.0
f = 0.7
f = 0.8
f = 0.9
Fixed
g = 50
g = 100

**Figure S67**. Power difference between unphased implementations of iHS (A, C, and E) and nSL (B, D, and F) and phased implementations. Blue curves represent the power difference between the unphased and phased statistics when applied to unphased data (UN). Red curves represent the power difference between the unphased and phased statistics when applied to perfectly phased data (PH). Values greater than 0 indicate the unphased statistic had higher power. Applied to demographic histories Demo 1 (A and B), Demo 3 (C and D), and Demo 4 (E and F) with n = 10 diploid samples. $s$ is the selection coefficient, $f$ is the frequency of the adaptive allele at time of sampling, $g$ is the number of generations at time of sampling since fixation, $e$ is the frequency at which selection began, and $t_d = 4000$ is the time in generations since the two populations diverged.

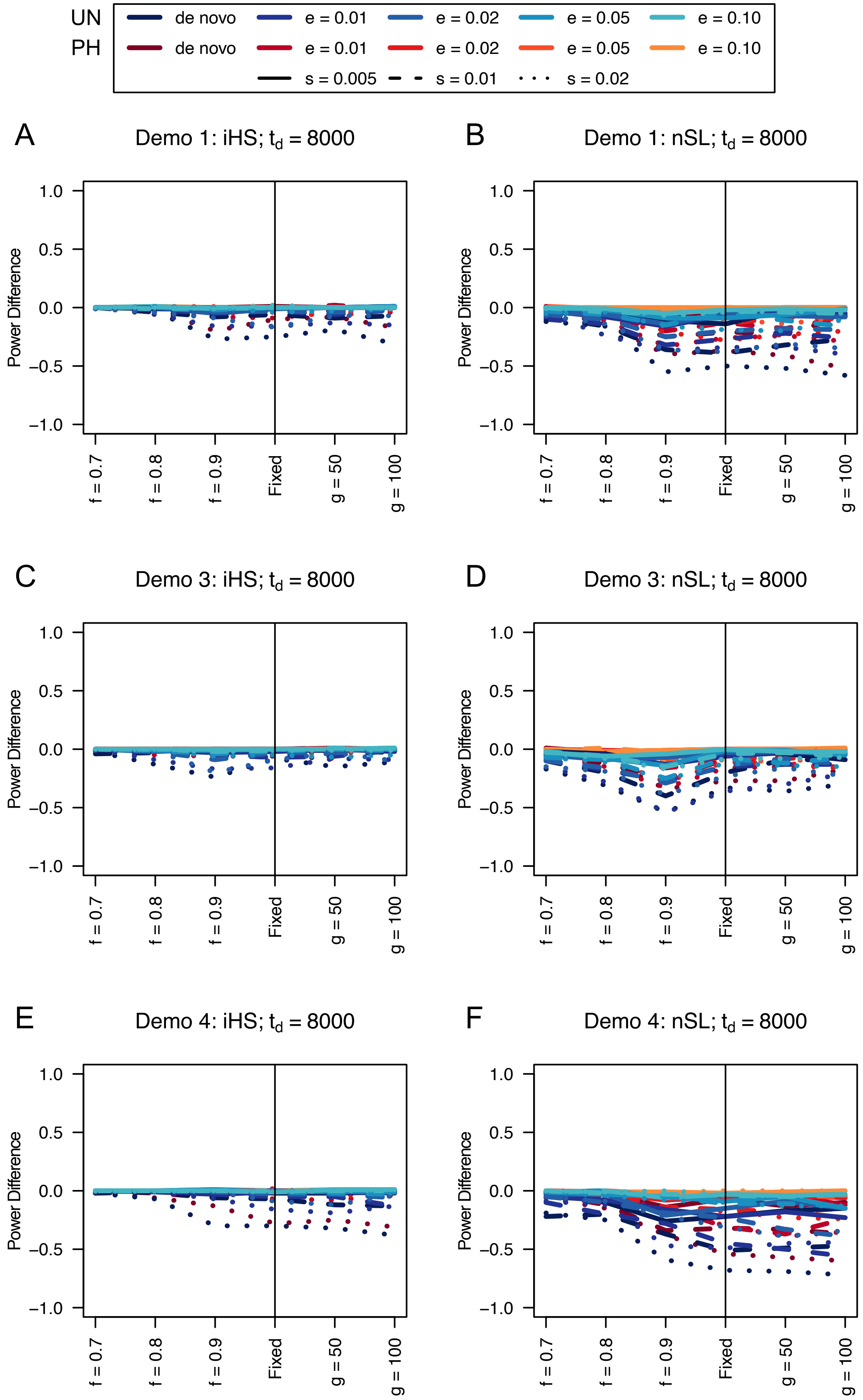
UN
PH
de novo
e = 0.01
e = 0.02
e = 0.05
e = 0.10
de novo
e = 0.01
e = 0.02
e = 0.05
e = 0.10
s = 0.005
s = 0.01
s = 0.02
A
Demo 1: iHS; t_d = 8000
B
Demo 1: nSL; t_d = 8000
C
Demo 3: iHS; t_d = 8000
D
Demo 3: nSL; t_d = 8000
E
Demo 4: iHS; t_d = 8000
F
Demo 4: nSL; t_d = 8000
Power Difference
1.0
0.5
0.0
−0.5
−1.0
f = 0.7
f = 0.8
f = 0.9
Fixed
g = 50
g = 100

737

**Figure S68**. Power difference between unphased implementations of iHS (A, C, and E) and nSL (B, D, and F) and phased implementations. Blue curves represent the power difference between the unphased and phased statistics when applied to unphased data (UN). Red curves represent the power difference between the unphased and phased statistics when applied to perfectly phased data (PH). Values greater than 0 indicate the unphased statistic had higher power. Applied to demographic histories Demo 1 (A and B), Demo 3 (C and D), and Demo 4 (E and F) with n = 10 diploid samples. $s$ is the selection coefficient, $f$ is the frequency of the adaptive allele at time of sampling, $g$ is the number of generations at time of sampling since fixation, $e$ is the frequency at which selection began, and $t_d = 8000$ is the time in generations since the two populations diverged.

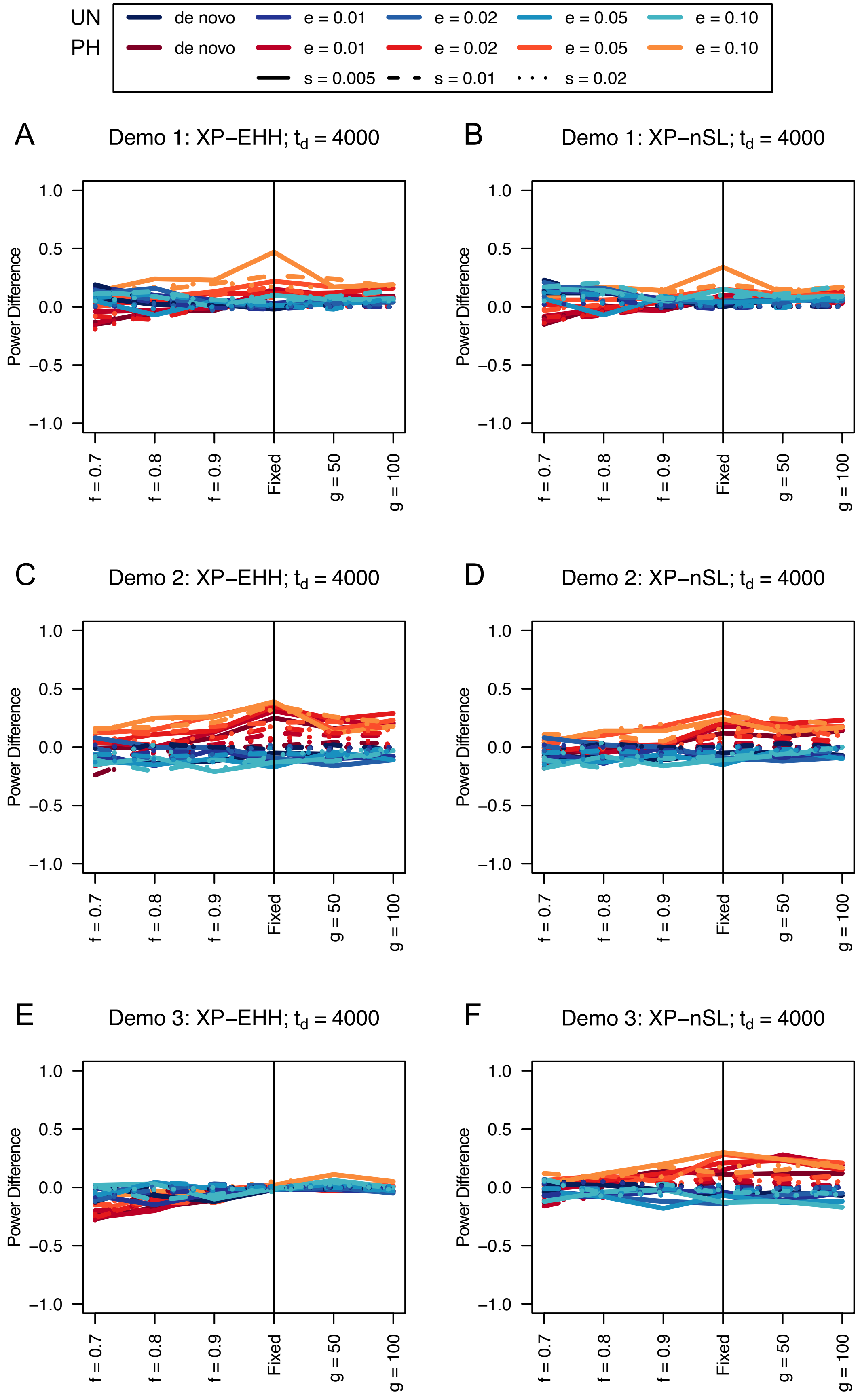
UN
PH
de novo
e = 0.01
e = 0.02
e = 0.05
e = 0.10
s = 0.005
s = 0.01
s = 0.02
A
Demo 1: XP−EHH; $t_d$ = 4000
B
Demo 1: XP−nSL; $t_d$ = 4000
C
Demo 2: XP−EHH; $t_d$ = 4000
D
Demo 2: XP−nSL; $t_d$ = 4000
E
Demo 3: XP−EHH; $t_d$ = 4000
F
Demo 3: XP−nSL; $t_d$ = 4000
Power Difference
1.0
0.5
0.0
−0.5
−1.0
f = 0.7
f = 0.8
f = 0.9
Fixed
g = 50
g = 100

747

**Figure S69**. Power difference between unphased implementations of XP-EHH (A, C, and E) and XP-nSL (B, D, and F) and phased implementations. Blue curves represent the power difference between the unphased and phased statistics when applied to unphased data (UN). Red curves represent the power difference between the unphased and phased statistics when applied to perfectly phased data (PH). Values greater than 0 indicate the unphased statistic had higher power. Applied to demographic histories Demo 1 (A and B), Demo 2 (C and D), and Demo 3 (E and F) with n = 10 diploid samples from each population. $s$ is the selection coefficient, $f$ is the frequency of the adaptive allele at time of sampling, $g$ is the number of generations at time of sampling since fixation, $e$ is the frequency at which selection began, and $t_d = 4000$ is the time in generations since the two populations diverged.

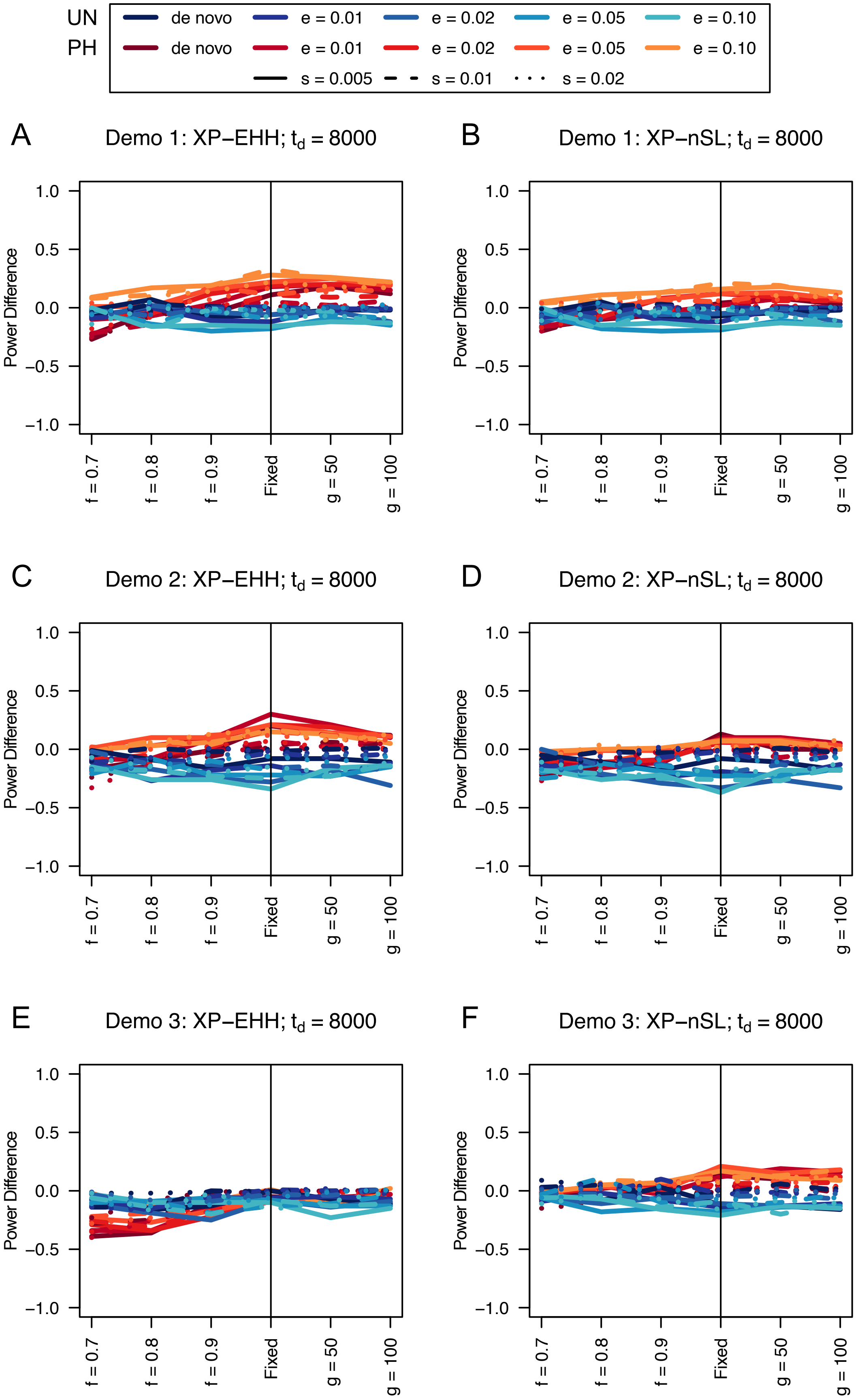

UN
PH
de novo
e = 0.01
e = 0.02
e = 0.05
e = 0.10
s = 0.005
s = 0.01
s = 0.02
A
Demo 1: XP−EHH; $t_d$ = 8000
B
Demo 1: XP−nSL; $t_d$ = 8000
C
Demo 2: XP−EHH; $t_d$ = 8000
D
Demo 2: XP−nSL; $t_d$ = 8000
E
Demo 3: XP−EHH; $t_d$ = 8000
F
Demo 3: XP−nSL; $t_d$ = 8000
Power Difference
1.0
0.5
0.0
−0.5
−1.0
f = 0.7
f = 0.8
f = 0.9
Fixed
g = 50
g = 100

**Figure S70**. Power difference between unphased implementations of XP-EHH (A, C, and E) and XP-nSL (B, D, and F) and phased implementations. Blue curves represent the power difference between the unphased and phased statistics when applied to unphased data (UN). Red curves represent the power difference between the unphased and phased statistics when applied to perfectly phased data (PH). Values greater than 0 indicate the unphased statistic had higher power. Applied to demographic histories Demo 1 (A and B), Demo 2 (C and D), and Demo 3 (E and F) with n = 10 diploid samples from each population. $s$ is the selection coefficient, $f$ is the frequency of the adaptive allele at time of sampling, $g$ is the number of generations at time of sampling since fixation, $e$ is the frequency at which selection began, and $t_d = 8000$ is the time in generations since the two populations diverged.

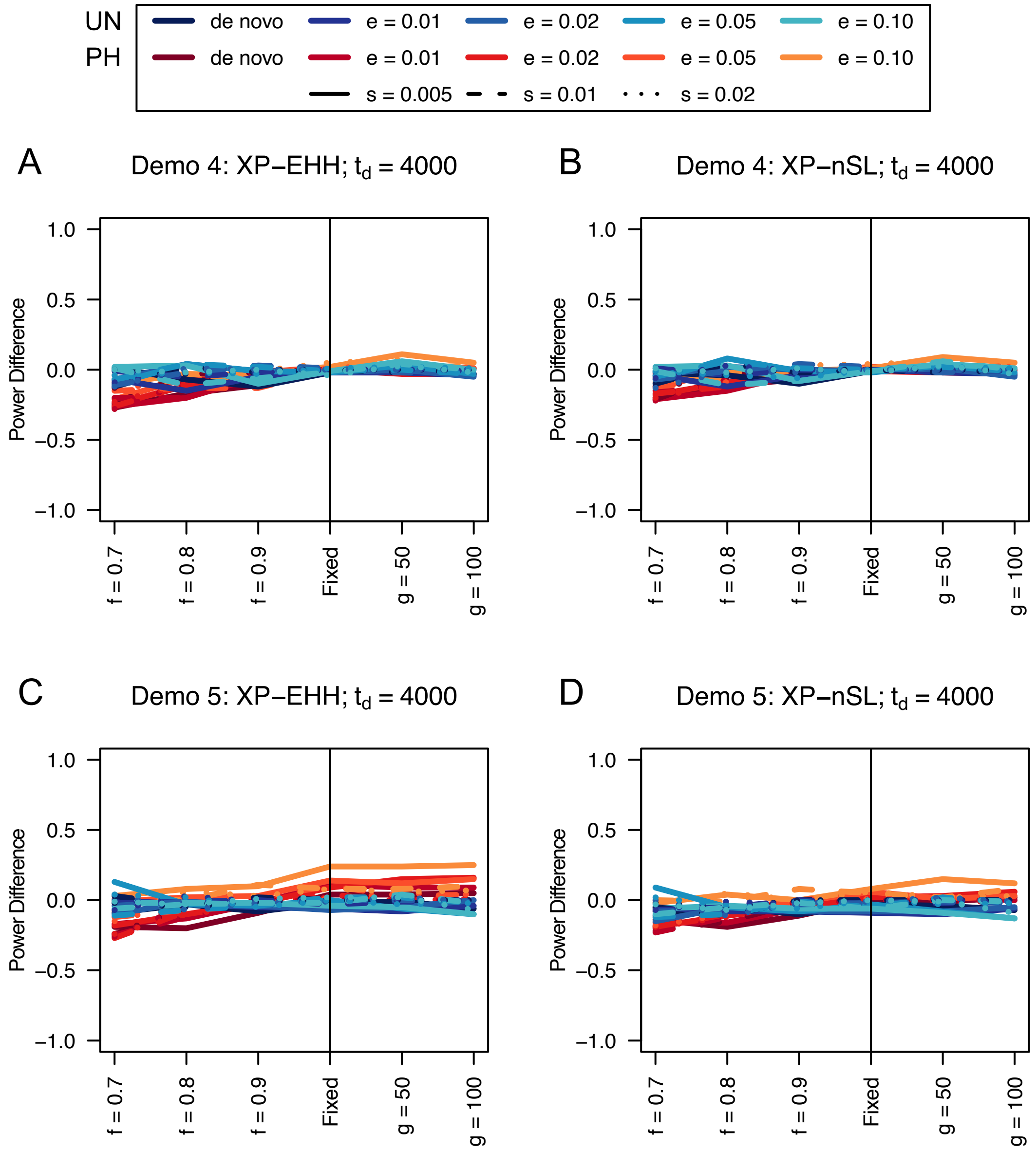


**Figure S71**. Power difference between unphased implementations of XP-EHH (A and C) and XP-nSL (B and D) and phased implementations. Blue curves represent the power difference between the unphased and phased statistics when applied to unphased data (UN). Red curves represent the power difference between the unphased and phased statistics when applied to perfectly phased data (PH). Values greater than 0 indicate the unphased statistic had higher power. Applied to demographic histories Demo 4 (A and B), and Demo 5 (C and D) with n = 10 diploid samples from each population. $s$ is the selection coefficient, $f$ is the frequency of the adaptive allele at time of sampling, $g$ is the number of generations at time of sampling since fixation, $e$ is the frequency at which selection began, and $t_d = 4000$ is the time in generations since the two populations diverged.

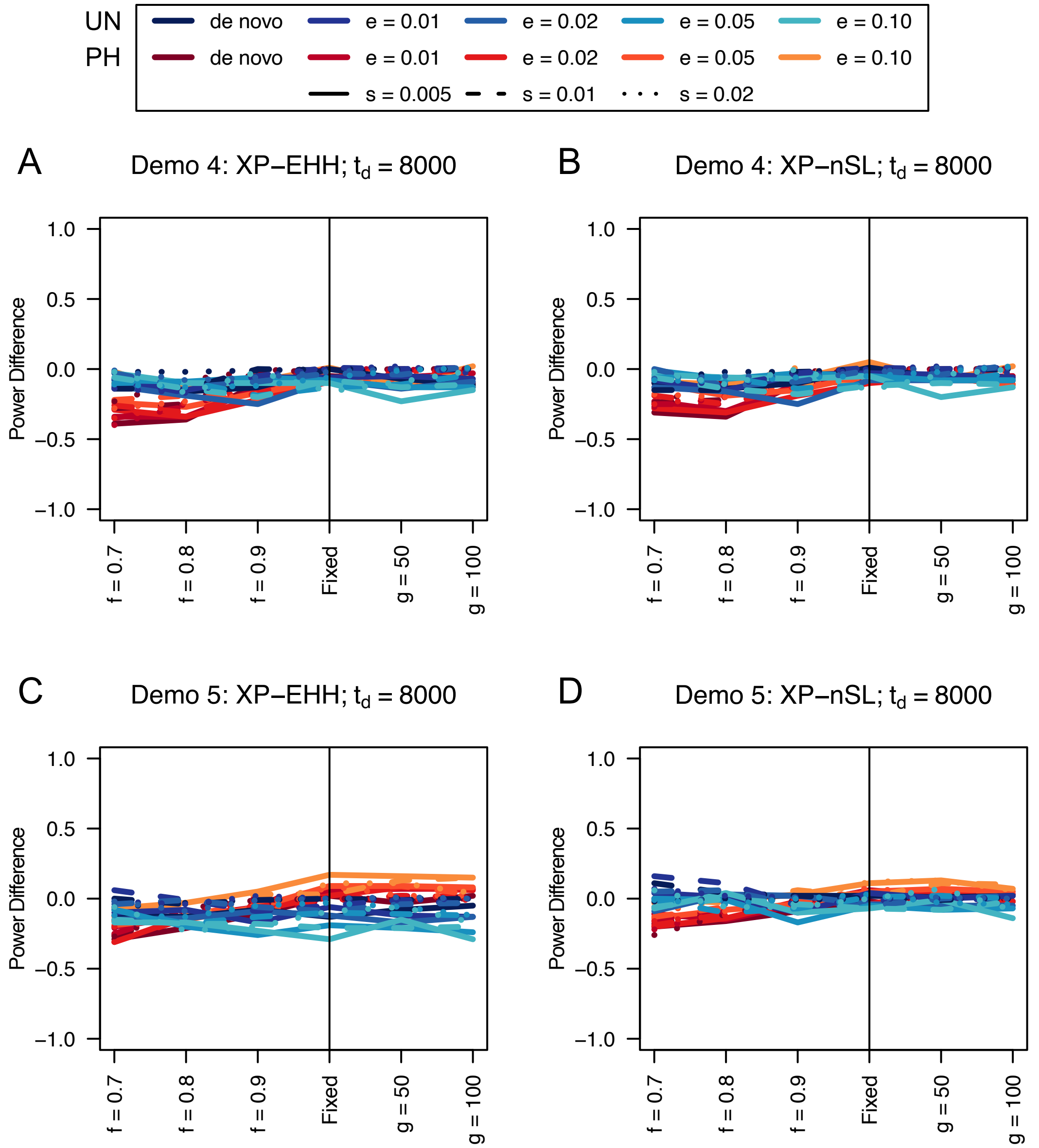


**Table S72**. Power difference between unphased implementations of XP-EHH (A and C) and XP-nSL (B and D) and phased implementations. Blue curves represent the power difference between the unphased and phased statistics when applied to unphased data (UN). Red curves represent the power difference between the unphased and phased statistics when applied to perfectly phased data (PH). Values greater than 0 indicate the unphased statistic had higher power. Applied to demographic histories Demo 4 (A and B), and Demo 5 (C and D) with n = 10 diploid samples from each population. $s$ is the selection coefficient, $f$ is the frequency of the adaptive allele at time of sampling, $g$ is the number of generations at time of sampling since fixation, $e$ is the frequency at which selection began, and $t_d = 8000$ is the time in generations since the two populations diverged.
